# Optimizing protein production in the One-Pot Pure system: insights into reaction composition and expression efficiency

**DOI:** 10.1101/2024.06.19.599772

**Authors:** Yan Zhang, Matas Deveikis, Yanping Qiu, Lovisa Björn, Zachary A. Martinez, Tsui-Fen Chou, Paul S. Freemont, Richard M. Murray

## Abstract

The One-Pot PURE (<u>P</u>rotein synthesis <u>U</u>sing <u>R</u>ecombinant <u>E</u>lements) system simplifies the preparation of traditional PURE systems by co-culturing and purifying 36 essential proteins for gene expression in a single step, thereby improving accessibility and affordability for widespread laboratory adoption and customization. However, replicating this protocol to match the productivity of traditional PURE systems can take considerable time and effort due to uncharacterized variability in the system’s biochemical composition. In this work, we observed unstable PURE protein expression in *E. coli* strains M15/pREP4 and BL21(DE3) and addressed this using glucose-mediated catabolite repression to minimize burdensome background expression. We also identified differences in optimal protein induction timing between these two strains, leading to growth incompatibility in co-culture, and observed proteolysis of PURE proteins expressed in M15/pREP4. We showed that consolidating all expression vectors into a protease-deficient BL21(DE3) strain could minimize proteolysis. This single-strain system also led to more uniform cell growth at the time of protein induction, improving the stoichiometry of critical translation initiation factors in the PURE reaction for efficient protein production. In addition to optimizing One-Pot PURE protein composition, we found that variations in commercial energy solution formulations could compensate for suboptimal PURE protein stoichiometry. Moreover, our study revealed significant differences in the expression capacity of commercially available *E. coli* tRNAs, suggesting the potential of optimizing tRNA codons to improve protein translation. Taken together, this work highlights the complex biochemical interplay influencing protein expression capacity in the One-Pot PURE system and presents strategies to improve its robustness and productivity.

## INTRODUCTION

Cell-free protein expression harnesses the core gene expression machinery in living cells to enable transcription and translation in a test tube reaction. The open nature of these expression platforms allows direct manipulation of the reaction environment. Applications of these platforms span high-throughput screening,^1–3^ novel protein modifications,^4,5^ interfacing biomolecules with synthetic materials,^6–9^ on-demand biosensing^10–12^ and biomanufacturing,^13,14^ and understanding the rules of life by building biological systems from scratch.^15–18^

Nearly all cell-free protein expression platforms can be categorized into two classes – crude lysate-based and reconstituted systems. Crude lysate-based expression systems use extracts from lysed cells, which retain most of the cell’s proteome and metabolic pathways.^19,20^ They can support gene expression from a wide range of promoters and enable higher protein expression due to effective energy regeneration.^21^ They can also be exceptionally affordable, costing approximately $0.03 per microliter of reaction.^22^ However, the active endogenous metabolism and partially intact proteome can divert critical energy sources and metabolites to side reactions, interfering with the central reactions implemented in these systems.^20^

In contrast, the PURE (<u>P</u>rotein synthesis <u>U</u>sing <u>R</u>ecombinant <u>E</u>lements) system isolates the necessary transcription, translation, and energy regeneration proteins to support gene expression by protein purification and reconstitution. This approach creates a more biochemically defined reaction environment, which enables the construction of predictive models for protein expression,^23,24^ allows facile removal and replacement of reaction enzymes for customization,^5^ and maintains a minimal reaction proteome toward a self-regenerating synthetic cell.^16^ However, traditional PURE systems are expensive to purchase ($0.64 – 1.00 per microliter of reaction) and laborious to prepare in-house, posing a significant bottleneck in the broad adoption of PURE systems.

To address these limitations, Lavickova and Maerkl developed the One-Pot PURE system, which offers a simplified alternative.^25^ Instead of growing and purifying each of the 36 PURE proteins, they grew a co-culture of the 36 *E. coli* strains, each expressing one of the PURE proteins, and purified the proteins in a single preparation. This approach significantly streamlines PURE system preparation and delivers protein expression yields comparable to commercial PURE systems but at a fraction of the cost ($0.10 per microliter of reaction).

Despite the promise of this method, replicating the One-Pot PURE protocol to achieve productivity comparable to conventional PURE systems can still take considerable time and effort. In this work, we identify sources of variability leading to reduced productivity in One-Pot PURE preparations, including expression instability, suboptimal PURE protein stoichiometry for effective expression, and uncharacterized protease activity during PURE protein purification. Moreover, we demonstrate that variations in the reaction’s biochemical composition can also significantly impact the protein production rate and the final protein yield, emphasizing the need for more precise system optimization. Taken together, our findings enhance the current understanding of factors influencing protein expression capacity in the One-Pot PURE system, providing strategies to enable robust and routinized implementation across diverse laboratories and applications.

## RESULTS AND DISCUSSION

### Enhancing protein expression stability for the One-Pot PURE system through glucose-mediated catabolite repression

We began by replicating the One-Pot PURE system using plasmids deposited on Addgene and transforming them into the respective *E. coli* expression strains M15/pREP4 and BL21(DE3). Plasmids containing PURE proteins encoded by pQE30 and pQE60 expression vectors were transformed into M15/pREP4 competent cells, with pREP4 plasmid coding for constitutive LacI expression. Plasmids containing PURE proteins encoded by pET21 and pET15 expression vectors were transformed into BL21(DE3) competent cells (**Figure 1A**).

**Figure 1:**
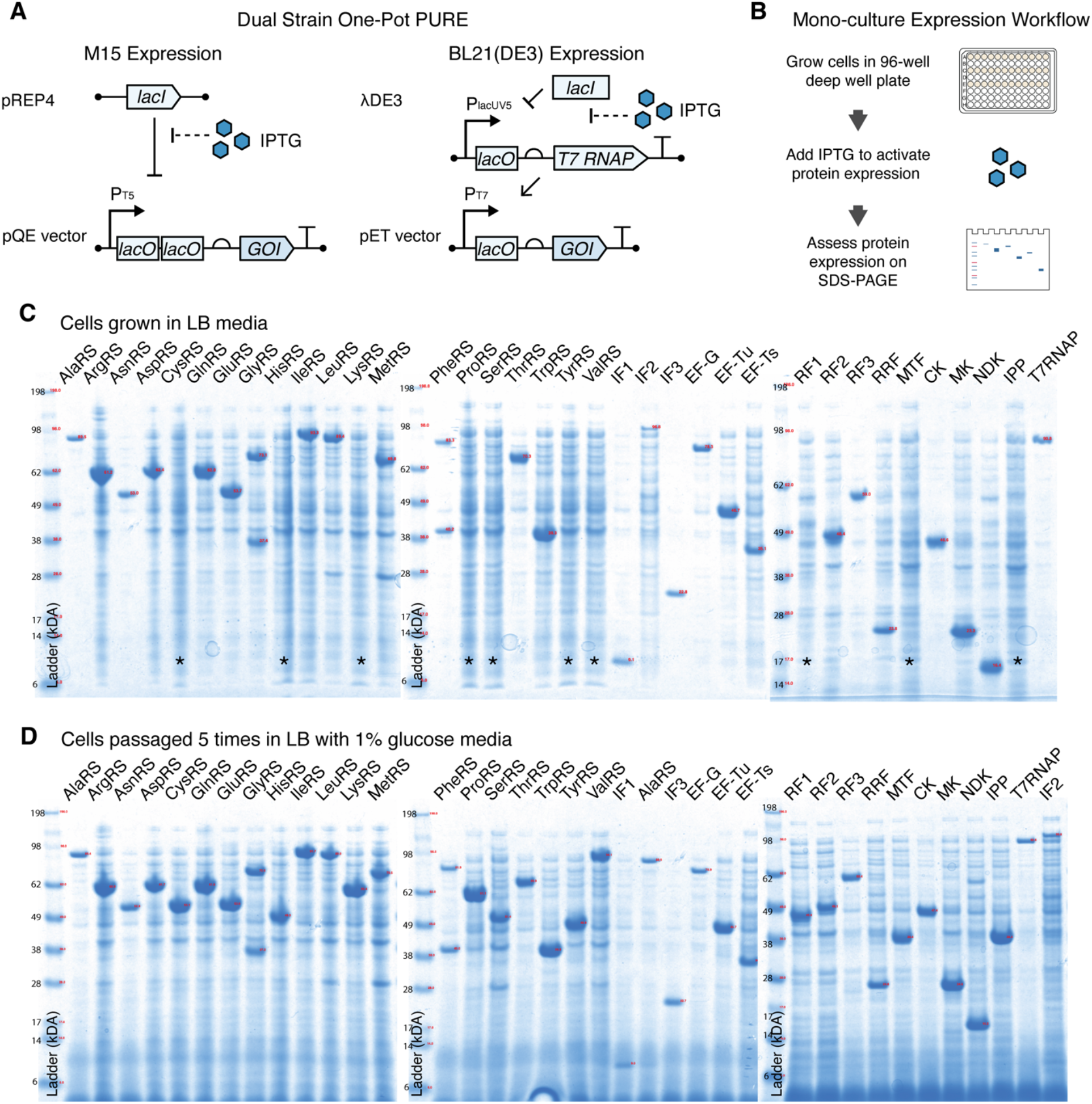
**(A)** Schematic of protein expression and regulation in *E. coli* strains M15/pREP4 and BL21(DE3) for Dual Strain One-Pot PURE systems. Each gene of interest is expressed under the control of either the P_T5_ or P_T7_ promoter followed by lacO operator site(s). In the M15/pREP4 expression strain, the pREP4 plasmid constitutively expresses the *lac* inhibitor, LacI, which represses protein expression by binding to the lacO sites. In BL21(DE3), endogenous LacI represses the production of T7 RNA polymerase (RNAP) and gene expression by binding to the lacO sites. Addition of the IPTG inducer de-represses LacI inhibition and activates gene expression. **(B)** Schematic of monoculture protein expression workflow. Cells were grown in 96-well deep-well plates, and protein expression was activated by adding 0.1 mM IPTG inducer. Following cell growth, protein expression was assessed via SDS-PAGE. **(C)** Monoculture protein expression assessment from cells grown in LB media. Ten strains (designated by *) were found to carry the protein expression plasmid but lacked the protein expression capability. **(D)** Stable protein expression from cells grown in LB with 1% w/v glucose after five serial passages, with all PURE proteins visibly expressed.

Our initial assessment of protein expression in monoculture assays revealed that several protein expressions were unstable. Despite confirming plasmid sequences through whole plasmid sequencing, we observed a loss of expression in ten proteins after growing and inducing protein expression from saved glycerol stocks (**Figure 1B, C**). No growth defect was observed for these non-expressing cell cultures (**Figure SI.III.1**). This phenomenon was also observed in several expression strains directly obtained from our collaborator, with the loss of protein expression appearing in other strains. This indicates that expression vector instability is a stochastic yet frequent event in PURE protein expression (**Figure SI.III.2**).

The expression vector instability is problematic because previously validated PURE protein expression strains can spontaneously lose protein expression after being saved to and grown from glycerol stocks. Without addressing this issue, subsequent preparations of the One-Pot PURE system may exhibit protein dropouts, leading to unproductive batches. Alternatively, each One-Pot PURE preparation would require fresh transformations of the 36 protein expression vectors, which could be labor-intensive and inefficient. Stabilizing the growth and expression of PURE proteins from glycerol stocks could significantly simplify the preparation workflow and facilitate the dissemination of these strains to other laboratories.

We hypothesize that leaky background protein expression from the P_T7-lacO_ promoter in BL21(DE3) and P_T5-lacO-lacO_ promoters in M15/pREP4 strains can create growth burdens, leading to the loss of protein expression. This phenomenon has been observed by others, in which background protein expression burden led to promoter mutations, resulting in the loss of target protein expression.^5^ To reduce background protein expression, we investigated whether catabolite repression – through the addition of a preferred carbon source (glucose) – could upregulate the intracellular concentration of repressors (LacI).^26^ This upregulation of LacI repressor could reduce background expression from P_T7-lacO_ and P_T5-lacO-lacO_ promoters.

Indeed, our results showed that by supplementing with growth media with 1% w/v glucose, PURE protein expression was stably maintained even after five serial passages from the seed glycerol stock (**Figure 1D**). For strains grown in catabolite repression media, we also observed improved cell growth and higher protein expression levels in ten PURE proteins previously found to lose expression ability (**Figure SI.III.3**). Based on these findings, all subsequent One-Pot PURE batches were grown with LB media supplemented with 1% w/v glucose.

### Offsets in the Dual Strain One-Pot PURE protein composition contribute to low protein production

After resolving the instability in protein expression, we proceeded to replicate the original One-Pot PURE system with BL21(DE3) and M15/pREP4 expression strains (referred to as Dual Strain One-Pot PURE) following the established protocol.^27^ To assess the system’s transcription and translation capacity, we used a reporter plasmid with a malachite green aptamer (MGA) to approximate mRNA concentration and a green fluorescent protein (deGFP) to measure protein yield (**Figure 2A**). Surprisingly, across two batches of the Dual Strain One-Pot PURE system, while the system produced high mRNA transcripts up to 6 *μ*M, protein expression was capped at approximately 1.5 *μ*M (**Figure 2B**). This protein yield is significantly lower than the yield we observed from commercial PURE systems (4-6 *μ*M, **Figure 3B, Figure 5D**). We also note that the high initial malachite green mRNA concentration at the start of the reaction may have come from rapid mRNA transcription that had begun before the microwell plate was placed into the plate reader.

**Figure 2:**
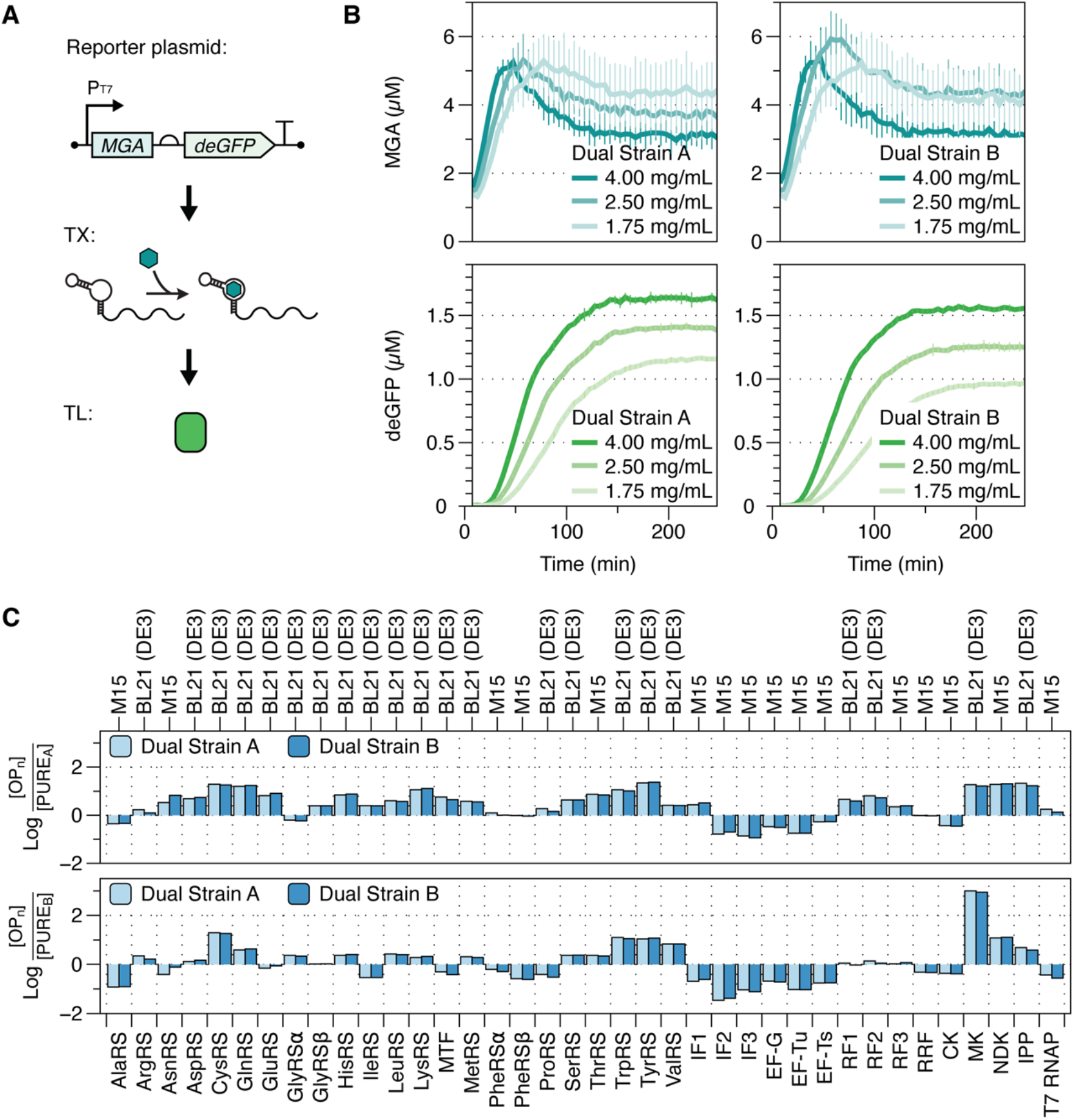
**(A)** Schematic of the reporter plasmid used to assess transcription and translation activity. The plasmid contains an RNA malachite green aptamer (MGA) to assess mRNA transcription and a green fluorescent protein reporter (deGFP) to measure protein expression. **(B)** Productivity of two batches of Dual Strain One-Pot PURE systems (Dual Strain A and Dual Strain B) in transcription and translation, measured via MGA and deGFP, respectively. The One-Pot PURE protein concentration in reactions was varied from 1.75 to 4 mg/mL to optimize protein yield. Plots represent the averages of reaction triplicates, with error bars indicating the standard deviations of reaction triplicates. **(C)** Comparison of protein abundances in Dual Strain One-Pot PURE systems to those in two commercial PURE systems, after scaling by concentrations in the reaction.

**Figure 3:**
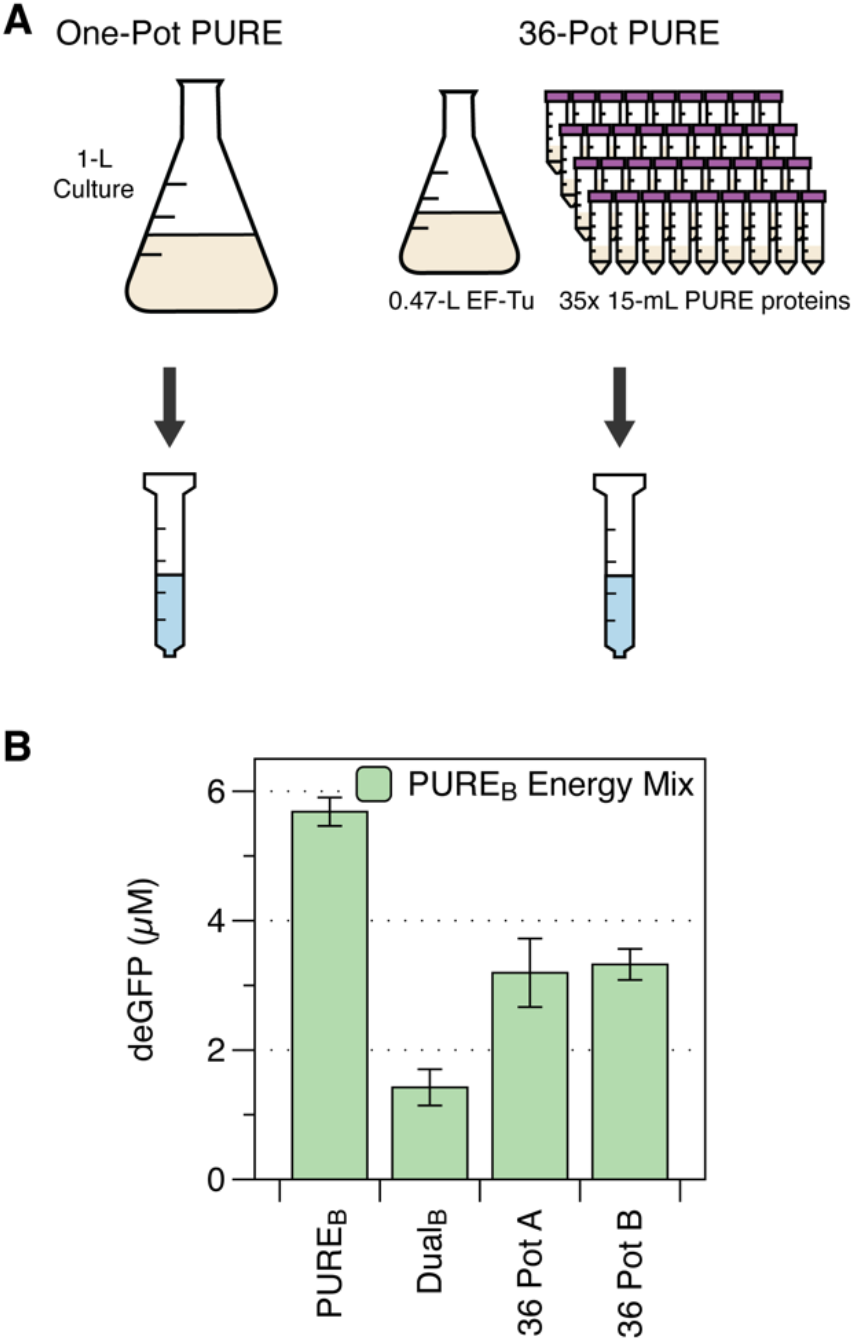
**(A)** Schematic showing the differences in preparation between the Dual Strain One-Pot PURE and 36-Pot PURE system. **(B)** PURE_B_ is a commercial PURE reaction assembled according to the manufacturer’s protocol. Dual Strain One-Pot PURE and 36-Pot PURE reactions were assembled with 5 mg/mL of PURE proteins. All reactions were prepared using a commercial energy solution (supplier B) and contained 5 nM of P_T7_-UTR1-deGFP plasmid. The bar graphs represent the average terminal protein yield of reaction triplicates, with error bars indicating the standard deviations of reaction triplicates.

Given that we had confirmed all strains were expressing the One-Pot PURE proteins in mono-culture expression assays, we hypothesized that there might be differences in cell growth or protein expression in the co-culture conditions not captured by the monoculture expression assays. However, using gel electrophoresis to resolve purified One-Pot PURE protein mixture, as done in the monoculture assays, would be challenging due to the similarity in molecular weight of many proteins. We, therefore, used a targeted proteomics approach to characterize the relative abundance of these proteins.

Using liquid chromatography with tandem mass spectrometry (LC-MS/MS), we mapped the detected peptides to protein abundances. After scaling for their concentrations in reactions, we compared the relative abundance of individual proteins in the Dual Strain One-Pot PURE mixture to those in commercial PURE systems (**Figure 2C**). Our analysis revealed a consistent deficiency in seven proteins in Dual Strain One-Pot PURE systems compared to the two commercial PURE systems (PURE_A_ and PURE_B_). These proteins are alanyl-tRNA synthetase (AlaRS), initiation factor 2 (IF2), initiation factor 3 (IF3), elongation factor-G (EF-G), elongation factor-Tu (EF-Tu), elongation factor-Ts (EF-Ts), and creatine kinase (CK).

### Bypassing Dual Strain One-Pot induction challenge in co-culture with 36-Pot PURE

Since all seven consistently deficient proteins were expressed in the *E. coli* M15/pREP4 expression strain, we investigated whether these cells had growth deficiencies. Monoculture growth data revealed that M15/pREP4 cells exhibited slower growth rates than BL21(DE3) cells, and most of them stopped growing after inducing protein expression (**Figures SI.III. 1 and 2**). This mismatch in growth rates created an incompatible induction window when co-culturing BL21(DE3) and M15/pREP4 cells for protein expression. To address this induction incompatibility, we explored growing the strains independently, followed by pooling them together at harvest for purification in a single preparation. We note here that a similar approach has been pursued as part of an open-source PURE protocol.^28^

This strategy, named the 36-Pot PURE, involved growing and expressing each of the 36 PURE proteins separately while maintaining the original inoculum composition for One-Pot PURE co-culture. For a 1-L scale preparation, the original One-Pot PURE method requires a 10-mL inoculum consisting of 47% v/v for elongation factor-thermal unstable (EF-Tu) culture and 1.5% v/v of each of the other proteins from saturated overnight cultures. In 36-Pot PURE, thirty-six cultures were grown: one 470 mL for EF-Tu growth culture and thirty-five 15 mL cultures for all other proteins (**Figure 3A**). Each culture was inoculated using a 1:100 dilution of a saturated overnight culture, grown to an induction OD above 0.6, induced for protein expression, and incubated for an additional 3 hours for protein expression.

Using this approach, the 36-Pot PURE system showed a twofold increase in deGFP yield compared to the Dual Strain One-Pot PURE system (**Figure 3B**). The improvement may be attributed to the mitigation of growth competition, as growing the strains separately prevented the faster-growing BL21(DE3) strains from outcompeting the M15/pREP4 cells in co-culture. The 36-Pot format also provided more flexibility in selecting optimal induction times for each protein, thus improving the final PURE protein composition. However, despite these improvements, the gene expression capacity still does not match the levels achieved by commercial PURE systems.

### T7 RNAP proteolysis in M15/pREP4 expression strain

In a separate attempt to purify T7 RNAP from the M15/pREP4 strain, we discovered protease activity in this expression strain that had not been reported previously. We found that, despite adding a broad-spectrum protease inhibitor (cOMPLETE™) in the lysis buffer, proteolysis of T7 RNAP into 80- and 20-kDa fragments was observed after protein purification, suggesting proteolysis occurred during cell lysis (**Figure 4**). This protease activity was not detected in earlier protein gel analyses, where cultures were heat-denatured immediately after protein expression (**Figure 1C, D**).

**Figure 4:**
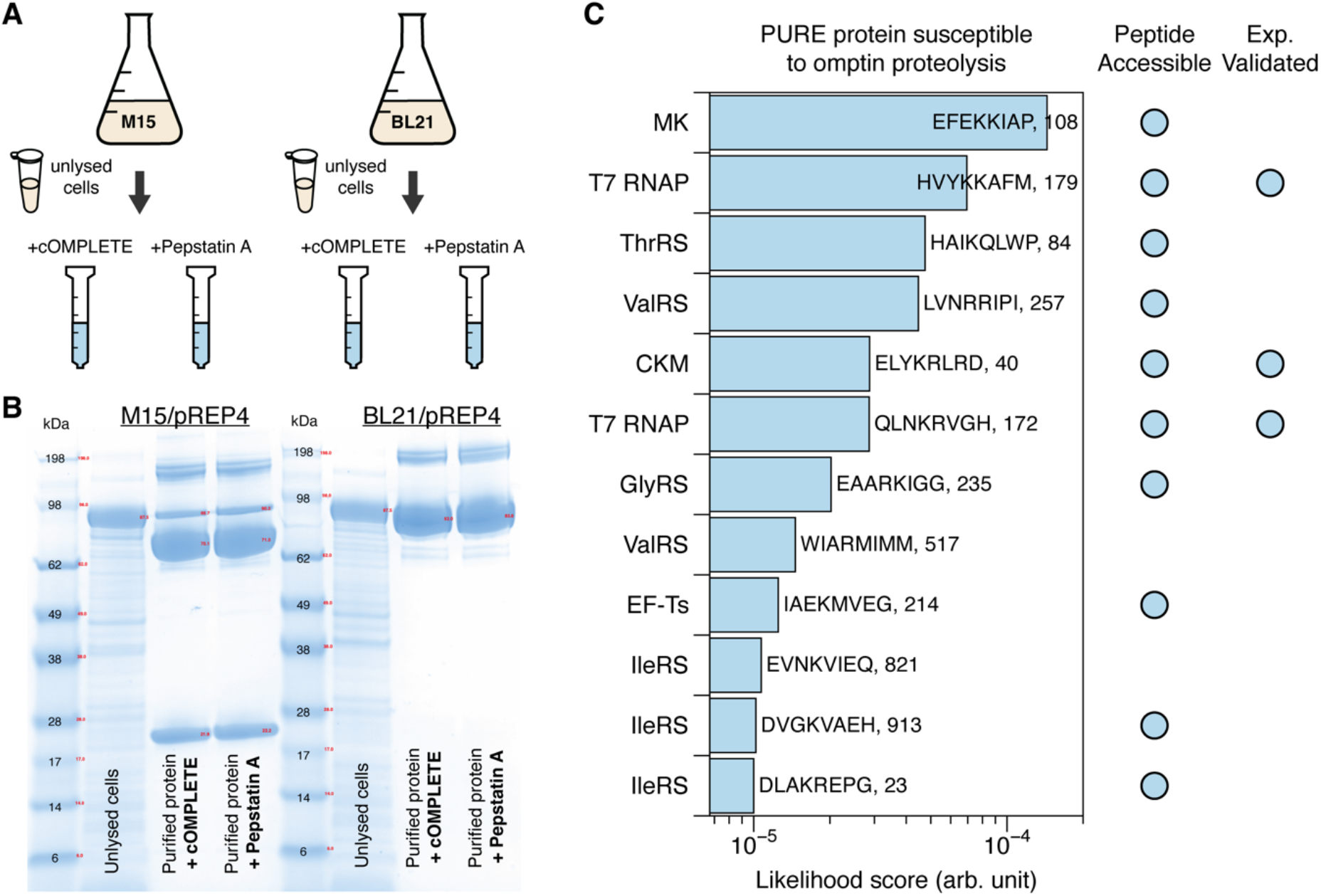
**(A)** Schematic of T7 RNAP preparation to assess outer membrane omptin family protease activity present in M15/pREP4 and BL21/pREP4 strains. A single culture of cells was grown, induced for T7 RNAP expression, and harvested, following a procedure similar to One-Pot PURE co-culture preparations. An un-lysed aliquot of cells expressing T7 RNAP was saved for SDS-PAGE analysis. The cultures were then split into two lysis and purification groups, each treated with a different protease inhibitor (cOMPLETE™ and Pepstatin A). **(B)** SDS-PAGE gel result showed T7 RNAP proteolysis when grown and purified in M15/pREP4 strains. Omptin proteases cleaved full-length T7 RNAP into truncated 80- and 20-kDa fragments. Proteolysis of T7 RNAP is absent in the BL21/pREP4 strain, which is deficient in the OmpT protease. **(C)** Simulated likelihood score of PURE proteins with peptide-subsequences susceptible to omptin proteolysis. The identified peptide sequences were assessed for their accessibility for omptin proteolysis (**SI. III. Figure 4**) and compared with existing literature. Only peptides for T7 RNAP and creatine kinase (CK) matched literature findings.^32,34^

Since M15/pREP4 is not a protease-deficient strain like BL21(DE3), we hypothesize that outer membrane omptin family proteases present in M15/pREP4 were responsible for cleaving T7 RNAP into truncated fragments. This is consistent with literature reports that omptins recognize and cleave T7 RNAP into 80- and 20-kDa fragments.^29^ Because omptins are aspartyl proteases unaffected by common protease inhibitors,^30^ no inhibition of protease activity was observed when lysis buffer was supplemented with Pepstatin A, an aspartyl protease inhibitor. This is consistent with reports of weak inhibition of omptins.^31^ In contrast, no proteolysis was observed when T7 RNAP was expressed in the BL21/pREP4 strain, which lacks one variant of the omptin family proteases, OmpT (**Figure 4A, B**).

It is also possible for omptins to cleave other PURE proteins with similar recognition motifs.^32^ Using established omptin cleavage motifs,^33^ we identified 12 possible omptin cleavage sites in eight PURE proteins (**Figure 4C**). The susceptible peptides identified for T7 RNAP and creatine kinase (CK) matched literature findings.^32,34^ We could not find published results confirming omptin-mediated proteolysis for the other six PURE proteins: myokinase (MK), elongation factor – thermal stable (EF-Ts), and threonyl-(ThrRS), valyl-(ValRS), glycyl-(GlyRS), isoleucyl-tRNA synthetases (IleRS). Using AlphaFold 3 simulations,^35^ we found that all but two peptides – one for ValRS and one for IleRS – are located on the external surface of the proteins and are accessible for protease binding (**Figure SI.III. 4**).

These simulations suggest that proteolysis of PURE proteins caused by omptins may contribute to the lower gene expression capacity observed during our optimization of the Dual Strain One-Pot PURE system. Since omptins can cleave full-length proteins into truncated fragments, even if these fragments can reassemble and be recovered from affinity column purification, their activities are likely to be reduced, as reported in the case of T7 RNAP.^36^

However, we expect the extent of proteolysis to be less significant in the One-Pot PURE and 36-Pot PURE preparation compared to what was observed in this T7 RNAP purification. This expectation arises from the high mRNA transcription activity observed with the Dual Strain One-Pot PURE system in an earlier experiment (**Figure 2A**). We hypothesize that the apparent impact of omptin proteolysis on the protein expression capacity of the Dual Strain One-Pot PURE and 36-Pot systems may be reduced by extra copies of T7 RNAP produced by BL21(DE3) cells and a lower abundance of omptins in the cell mixture with BL21(DE3) cells.

We also note that the M15/pREP4 strain was not used in the original development of the PURE system. All proteins were expressed in BL21/pREP4 and BL21(DE3) strains lacking the OmpT protease.^37^ As a result, we find it is critical to transfer all PURE protein expressions to a protease-deficient strain to ensure the production of full-length, active PURE proteins.

### Enhancing One-Pot PURE protein composition and expression capacity with a Single-Strain One-Pot PURE system

To prevent unwanted proteolysis and attain more uniform cell growth in the co-culture environment, we transferred all PURE proteins encoded in the pQE30 and pQE60 vectors for M15/pREP4 expression to pET21 vectors for BL21(DE3) expression. However, a caveat of this Single Strain One-Pot PURE system is that the T7 RNAP needs to be removed from the One-Pot PURE co-culture due to the genetic instability of having a P_T7_ promoter expressing T7 RNAP. As a result, we opted to purify and add T7 RNAP into One-Pot PURE reactions separately (see **MATERIALS AND METHODS**). Alternatively, T7 RNAP could be co-transformed with pREP4 plasmid carrying constitutive LacI expression into BL21 cells and grown in the One-Pot PURE co-culture.

Our results showed that five out of the 16 proteins previously expressed in M15/pREP4 cells exhibited at least a 30% improvement in expression levels when grown in BL21(DE3) cells, as determined by monoculture growth and SDS-PAGE analysis (**Figure SI.III. 5**). This enhancement is also evident in the Single Strain One-Pot PURE co-culture. Compared to the Dual Strain system, the purified PURE protein mixture from the Single Strain system has a slightly higher composition of translation factors (24% and 26% for Dual Strain and Single Strain, respectively, **Figure 5B**). Using mass spectrometry to resolve individual protein composition, we found that translation initiation factors 2 and 3 (IF2 and IF3) and creatine kinase (CK) involved in energy regeneration (**Figure 5A**) are present at significantly higher concentrations in Single Strain One-Pot PURE. An Individual protein composition comparison of Single Strain One-Pot PURE to commercial PURE systems is also provided in **Figure SI.III. 6**.

**Figure 5:**
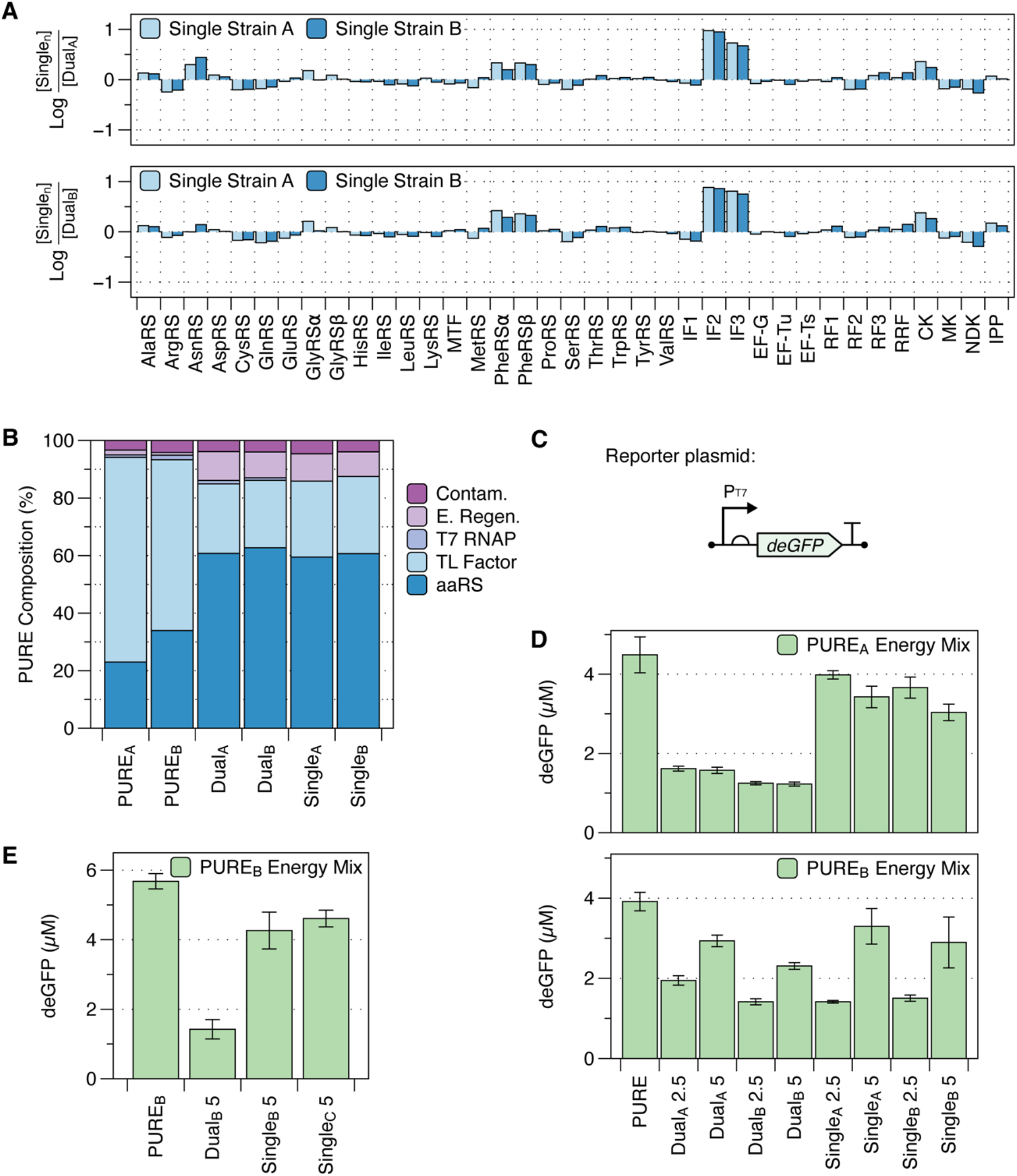
**(A)** Comparison of the relative abundance of individual One-Pot PURE proteins in the Single Strain and Dual Strain systems. Initiation factors 2 and 3 (IF2, IF3) show the most notable increase in composition in the Single Strain One-Pot PURE system. Previously under-expressed aminoacyl-tRNA synthetases (AlaRS and PheRS) and creatine kinase (CK) also increased in composition in the Single Strain One-Pot PURE system. **(B)** Composition of commercial PURE proteins, Dual Strain One-Pot PURE proteins, and Single Strain One-Pot PURE proteins grouped by protein classes: aminoacyl-tRNA synthetase (aaRS), translation factors (TL factors), T7 RNAP, energy regeneration (E Regen.). Detected proteins not part of the PURE proteins are categorized as contaminants (Contam.). Ribosomal proteins are excluded from the analysis. **(C)** Schematic of the reporter plasmid P_T7_-UTR1-deGFP expressed in PURE reactions to assess system productivity. **(D)** Productivity of Dual Strain and Single Strain One-Pot PURE reactions assembled with two commercial energy solutions. PURE_A_ and PURE_B_ are control reactions with 5 nM P_T7_-UTR1-deGFP plasmids assembled according to the manufacturer’s protocol. “Dual_n_ 2.5” and “Dual_n_ 5” designate varying batches of Dual Strain One-Pot PURE proteins added at 2.5 and 5 mg/mL. Likewise, “Single_n_ 2.5” and “Single_n_ 5” designate varying batches of Single Strain One-Pot PURE proteins added at 2.5 and 5 mg/mL. The bar graphs represent the average terminal protein yield of reaction triplicates, with error bars indicating the standard deviations of reaction triplicates. **(E)** Productivity comparison of the collaborator’s batch of Single Strain One-Pot PURE (Single_C_). PURE_B_ is the control reaction assembled according to the manufacturer’s protocol. Single Strain and Dual Strain One-Pot PURE reactions were assembled using 5 mg/mL of PURE protein, using the commercial energy solution from supplier B and 5 nM of P_T7_-UTR1-deGFP plasmid. The bar graphs represent the average terminal protein yield of reaction triplicates, with error bars indicating the standard deviations of reaction triplicates.

We next compared the productivity of the Dual Strain and Single Strain versions of the One-Pot PURE system. Using only the energy solution from commercial vendors and replacing commercial PURE proteins with One-Pot PURE proteins, we observed consistently higher protein production with the Single Strain One-Pot PURE system (**Figure 5C, D**). The robustness of the Single-Strain system was further validated through a reproducibility study conducted in our collaborator’s lab. A new batch of Single Strain One-Pot PURE system was prepared following the same protocol (referred to as Single_C_), and its productivity was compared to shipped aliquots of previous batches, Single_B_ and Dual_B_. Even in a different laboratory setting, using separate equipment and reaction consumables, the higher yield of the Single Strain system was reproduced on the first attempt. This was observed both with commercial energy systems (**Figure 5E**) and with homemade energy solutions (**Figure SI.III. 7**)

### High-yield protein expression from One-Pot PURE requires a balance between the protein production rate and energy solution formulation

The differences in productivity between the Dual Strain One-Pot PURE system when using two commercial PURE energy solutions, referred to as PURE_A_ and PURE_B_, are particularly interesting. It appears that higher amounts of One-Pot PURE proteins can be added in reactions with PURE_B_ energy solution as a strategy to compensate for a sub-optimal stoichiometry of translation factors in the purified protein mixture (**Figure 5D**). In reactions with the PURE_A_ and homemade energy solution, increasing the concentration of One-Pot PURE proteins from 2.5 mg/mL to 5 mg/mL decreased protein production.

Although we do not know the exact formulation of these energy solutions, comparing the protein production rates (coupled transcription-translation rates) of One-Pot PURE systems to commercial PURE systems provides valuable insights (**Figure 6** and **Figure SI.III 8-9**). In the PURE_A_ energy solution, the protein production rates of the Dual Strain and Single Strain One-Pot PURE systems reflected the anticipated PURE protein differences based on their translation factor content (**Figure 6**). Deficiencies in translation factors in the Dual Strain One-Pot PURE system contributed to a lower production rate over the course of the reaction. With more protein translation machinery, the Single Strain One-Pot PURE system could match or even exceed the production rate of commercial PURE_A_ proteins. However, higher production rate peaks led to steeper rate declines shortly after, as observed for the Single Strain One-Pot PURE system with the PURE_A_ energy solution.

**Figure 6:**
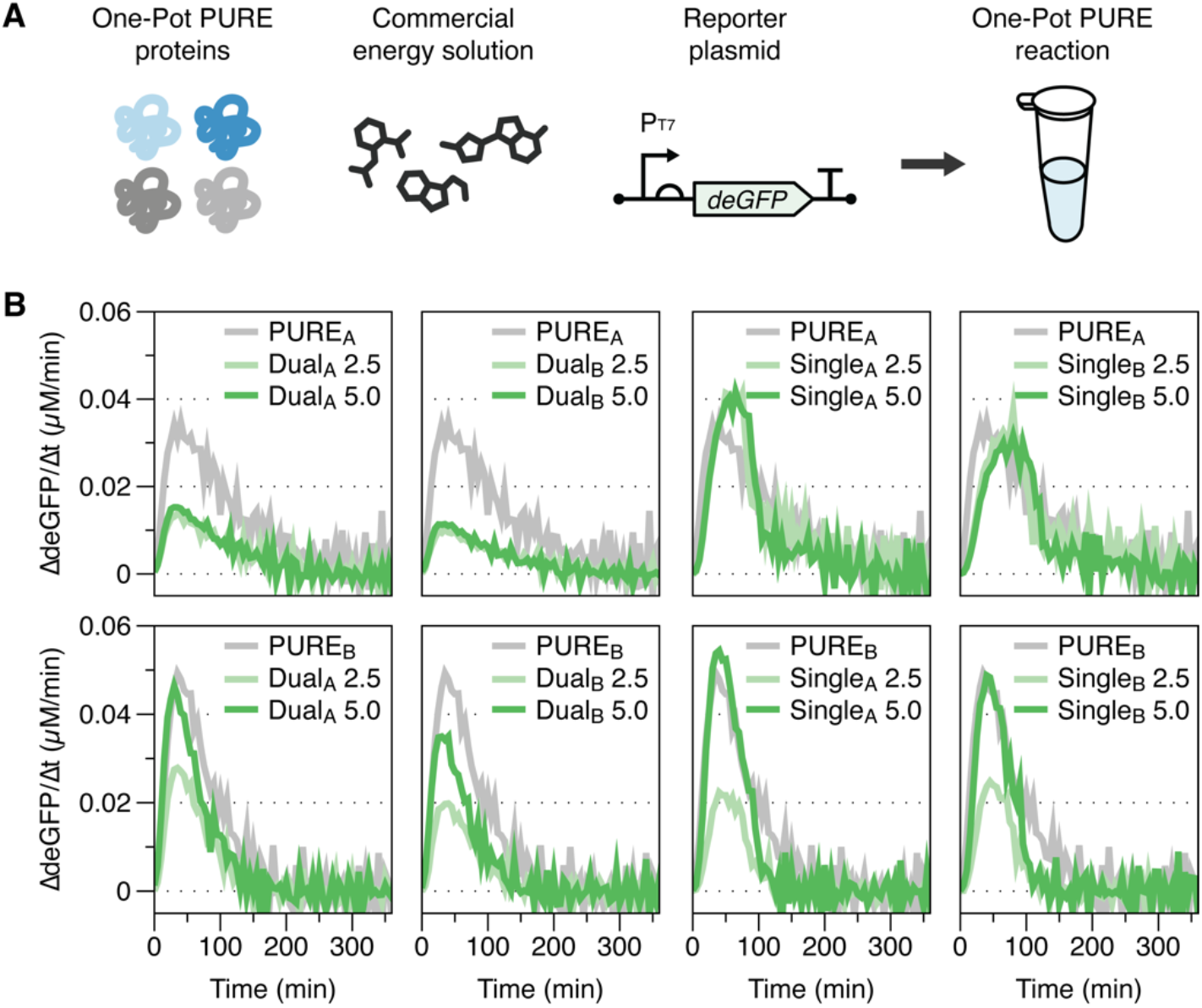
**(A)** Schematic of One-Pot PURE reaction composition using Dual Strain and Single Strain One-Pot PURE proteins with commercial energy solutions, assembled according to manufacturers’ protocols. All reactions used 5 nM of P_T7_-UTR1-deGFP reporter plasmid. In the case of Single Strain One-Pot PURE reactions, 0.5 *μ*M of purified T7 RNAP was added. **(B)** Comparison of protein translation rate for various preparations of One-Pot PURE proteins (added to the reaction at 2.5 or 5.0 mg/mL) using two commercial energy solutions. Plots represent the average of reaction triplicates. Time traces of deGFP production are provided in the **Supplementary Figures SI.III. 8-9**

In the PURE_B_ energy solution, when both the Dual Strain and Single Strain One-Pot PURE proteins are added to 5 mg/mL, both systems exhibited comparable protein expression levels despite the deficiency in translation factors found in the Dual Strain One-Pot PURE proteins (**Figure 6**). It is important to note that the contribution to these differences in translation rate comes from the energy solution alone (amino acids, NTPs, tRNA, creatine phosphate, and other co-factors). All reactions were assembled with the same reporter plasmids and ribosomes. This result suggests that an optimal preparation of the energy solution could compensate for an otherwise sub-optimal One-Pot PURE protein stoichiometry in reactions.

We found it intriguing that when the production rate from the One-Pot PURE system exceeded that of commercial PURE proteins, a steeper drop in production rate soon followed (**Figure 6**). We first attempted to explain the phenomenon from the perspective of the system’s energy regeneration capability. While fast protein production can occur, it can also significantly deplete the available energy molecules (ATP and GTP) critical for powering translation initiation and elongation reactions. A fast production rate surpassing the system’s energy regeneration capacity can negatively impact the final protein yield by stalling protein translation, potentially resulting in premature translation termination and lower apparent protein production. This corroborates the finding that supplementing an additional energy regeneration system to the PURE reaction can help attain a higher maximum protein production rate.^38^

However, the energy regeneration bottleneck can only explain the protein production behavior observed in reaction with PURE_A_ energy solution and Single Strain One-Pot PURE proteins. If energy regeneration was the only bottleneck in protein production, then reactions with suboptimal PURE protein stoichiometry (Dual Strain One-Pot PURE) should have a lower yet longer-lasting maximal protein production rate in the PURE_A_ energy solution. Similarly, energy regeneration alone cannot explain the discrepancy between the Dual Strain and Single Strain One-Pot PURE expression behaviors in PURE_A_ and PURE_B_ energy solutions.

These observations lead us to consider additional factors contributing to differences in protein expression rates. One component often overlooked in optimizing cell-free expression capacity is the tRNA composition and codon usage. Because the exact tRNA composition for each codon in the PURE reaction is not known, and protein translation is influenced by codon usage, it is often challenging to optimize the expressed protein’s codon usage to the tRNA composition in the reaction.

Since the same reporter plasmid was used for Dual Strain and Single Strain One-Pot PURE reactions in different PURE energy solutions, the differences in final yield and maximal production rate may be influenced by the tRNA codon composition in different energy formulations. We hypothesize that a sub-optimal tRNA composition in the reaction, not matching well with the codon usage of the reporter protein, may also contribute to ribosome stalling on the mRNA transcript when encountering sub-optimal codons.^39^ This could be exacerbated by an mRNA transcript with a high translation initiation rate, such as the one used in the reporter plasmid (UTR1 ribosomal binding site),^40^ which increases the frequency of ribosomal collisions. Colliding ribosomes could lead to ribosomal fall-off, resulting in a low apparent yield of protein translation. This also corroborates with reports of C-terminal truncated protein produced in PURE reactions.^41^

### E. coli tRNA composition critically influences One-Pot PURE reaction productivity

We further demonstrated the important influence of suboptimal tRNA composition on protein production by preparing homemade energy solutions using two different sources of commercial *E. coli* tRNA mixtures: one from *E. coli* strain MRE600 and one from *E. coli* strain W. In these experiments, the same concentrations of the Single Strain One-Pot PURE proteins, reporter plasmids, ribosomes, and T7 RNAP were used. The only difference between the reactions was the source – and possibly the composition – of *E. coli* tRNA in the energy solution.

It was observed that the energy solution prepared using *E. coli* MRE600 tRNA exhibited a high protein production rate and final protein yield (**Figure 7B**). When we replaced tRNA in the energy solution with *E. coli* Strain W tRNA, we observed a significant decrease in protein production rate and yield (**Figure 7C**). Increasing the concentration of *E. coli* Strain W tRNA did not improve the rate and yield of protein expression.

**Figure 7:**
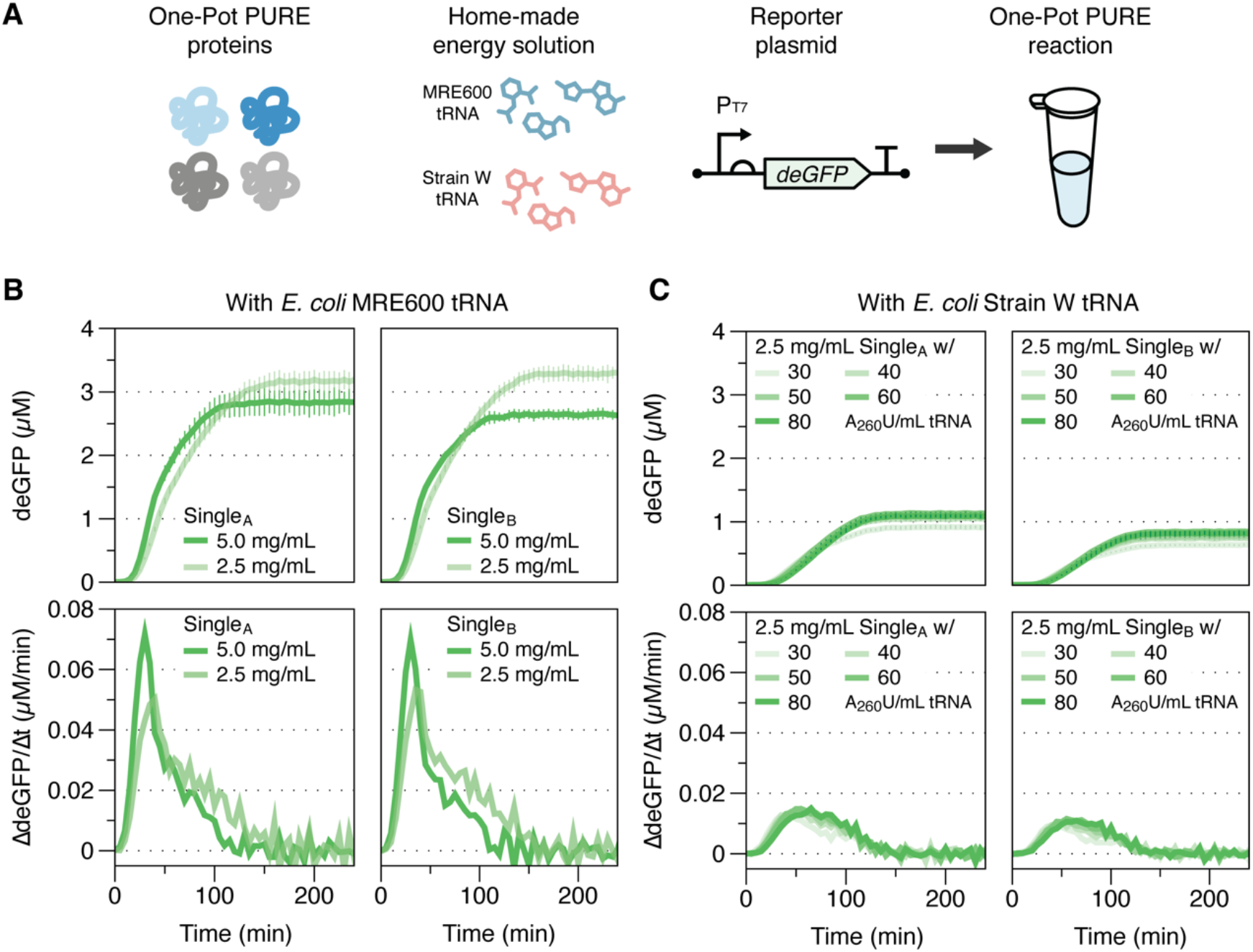
**(A)** Schematic of One-Pot PURE reaction composition using in-house prepared energy solutions with commercial tRNAs from *E. coli* MRE600 and *E. coli* Strain W. Each reaction contained 5 nM P_T7_-UTR1-deGFP to assess protein production. **(B)** Comparison of deGFP yield and production rate for two batches of Single Strain One-Pot PURE proteins (added to the reaction either at 2.5 or 5.0 mg/mL) in energy solution with 22 A_260_U/mL *E. coli* MRE600 tRNA. Error bars represent standard deviations of reaction triplicates. Protein production rate plots represent the average of reaction triplicates. **(C)** Comparison of deGFP yield and production rate for two batches of Single Strain One-Pot PURE proteins (added to the reaction at 2.5 mg/mL) in energy solution with increasing *E. coli* Strain W tRNA activity units (30 – 80 A_260_U/mL). The plots represent the average of reaction triplicates, and the error bars represent standard deviations of reaction triplicates. Protein production rate plots represent the average of reaction triplicates.

This finding is significant because the complexity and composition of tRNAs can be overlooked when preparing the energy solution for PURE reactions. Differences in codon usage among various *E. coli* strains can lead to unexpected deficiency in tRNAs associated with the codons used. Without prior knowledge of these tRNA deficiencies, it can be difficult to codon-optimize the protein for effective expression, resulting in low apparent protein yield despite an otherwise productive system.

## CONCLUSION

In this work, we identified several important factors that affect the expression capacity of the One-Pot PURE system. We improved the stability of One-Pot PURE protein expression through glucose-mediated catabolite repression and mitigated potential protease degradation from M15/pREP4 expression strains by consolidating all protein expression vectors to one *E. coli* BL21(DE3) strain. This new single-strain approach also increased the expression levels of critical translation initiation and energy regeneration proteins and, consequently, the productivity of the One-Pot PURE reaction. Our characterizations of both dual and single-strain One-Pot PURE proteins in two commercial PURE energy solutions revealed that optimizing biochemical compositions could overcome deficiencies in translation capacity. Lastly, our in-house energy solution preparation revealed the critical role of tRNA composition in determining the translation capacity of PURE reactions.

These findings showcased that the gene expression capacity of the PURE system is influenced by both PURE protein stoichiometry and biochemical factors. At the protein level, we found that when translation initiation factors (initiation factors 2 and 3, IF2 and IF3) and energy regeneration protein, creatine kinase (CK) are present at higher concentrations in the PURE protein mixture, higher expression capacity is observed (**Figure 5D, E**).

At the biochemical level, we found that energy solution composition significantly impacts the One-Pot PURE production rate and the final protein yield. Using the PURE_A_ energy solution, the reaction proceeded with a slower protein production rate and took over 6 hours to stabilize to the final deGFP concentration (**Figure SI.III 8**). This suggests that there may be a reaction rate-limiting step imposed by the PURE_A_ energy solution, and this limitation can make One-Pot PURE proteins with sub-optimal protein composition proceed even more slowly, resulting in lower protein yield. On the other hand, the PURE_B_ energy solution supports a faster protein production rate overall, with reactions completed in under 3 hours (**Figure SI.III 9**). This suggests the rate-limiting step observed in the PURE_A_ energy solution may be removed in PURE_B_ to deliver a high protein yield with a shorter reaction lifetime. We hypothesize this rate-limiting step can be the reaction’s tRNA composition. We also note that PURE_A_ and PURE_B_ energy solutions are proprietary information from commercial vendors. The exact compositions of these energy solutions are not known to us, which limits the conclusions we can draw.

Lastly, we show that commercial preparations of *E. coli* tRNA sourced from strain MRE600 and strain W significantly impact effective protein translation (**Figure 7**). Although MRE600 is a highly preferred source for *E. coli* tRNA due to the strain’s low RNase I activity,^42^ there is no apparent reason behind tRNA obtained from strain W (a safe and fast-growing strain utilizing sucrose as an industrially preferred carbon source)^43^ exhibiting sub-optimal translation capacity. This highlights a critical limitation and source of variability in most cell-free systems (lysate-based and PURE), which is that *E. coli* tRNA is also a complex mixture and exhibits differential concentration profiles across growth stages and host strains.^44,45^ For robust control over both One-Pot PURE reaction and potentially lysate-based cell-free expression systems, a promising approach can be developing an optimal tRNA composition.^46^

Taken together, this work seeks to obtain a comprehensive understanding of the factors influencing the gene expression capacity of the One-Pot PURE system. By addressing both protein and biochemical composition, our findings offer valuable insights into optimizing not only One-Pot PURE systems but also extending to lysate-based cell-free protein synthesis, paving the way for more efficient and reliable applications in synthetic biology and biotechnology.

## MATERIALS AND METHODS

### Escherichia coli strains and plasmids

*E. coli* BL21(DE3) (Thermo Scientific) and M15/pREP4 (Qiagen) strains were used for protein expression. **Supplementary Information I** contains the Addgene catalog numbers and plasmid sequences for Dual-Strain One-Pot. To establish the Single Strain One-Pot, all proteins were amplified via Gibson Assembly and placed onto the pET21a backbone. All sequences of cloned proteins transplanted onto pET21a are provided in **Supplementary Information I**.

Sequence-verified plasmids (Primordium Labs, now Plasmidsaurus) were transformed into their respective BL21(DE3) and M15/pREP4 expression strains and plated on Luria Broth (LB) Agar plates with 100 *μ*g/mL carbenicillin and 1% glucose. An inoculation loop was used to scrape multiple colonies to inoculate overnight culture in LB with 100 *μ*g/mL carbenicillin and 1% glucose to obtain the population expression average. To facilitate inoculation of subsequent final cultures, 250 *μ*L of overnight cultures were saved into frozen glycerol stocks with 250 *μ*L of 50% glycerol in 1.3 mL 96 well plates (NEST Biotech). The glycerol stock plate was sealed with an aluminum film and stored at -80 °C.

Sequences of plasmid DNA used for reporter protein expression are provided in **Supplementary Information I**. All plasmid DNA was prepared using the NucleoBond Xtra Midi kit (Macherey-Nagel) and eluted in nuclease-free water.

### Monoculture Expression assay for One-Pot PURE proteins

The protocol was adapted from Grasemann *et al*.27 with a few modifications. An overnight culture of One-Pot PURE protein expression strains was inoculated either from freshly transformed cells grown on LB agar plates with carbenicillin (without or with 1% w/v glucose to exert catabolite repression) or from frozen glycerol stocks using a cryo-replicator (Enzyscreen, CR1000). The overnight culture was grown in 96-well deep-well plates sealed with a Breathe-Easier membrane (Diversified Biotech) at 37 °C and 1000 rpm.

The next morning, a fresh starter culture was prepared using 4 *μ*L of the overnight cells and 396 *μ*L of LB with carbenicillin. OD600 was monitored at the time of 0.1 mM IPTG induction to initiate protein expression after 2-2.5 hrs of growth and 3 hours after protein expression. 10 *μ*L of cells expressing One-Pot PURE proteins were mixed with Lammeli Buffer (Bio-Rad) and heat denatured at 95 °C for 10 minutes, and 10 *μ*L of the mixture was loaded onto Bolt™ 4-12% Bis-Tris Protein Gels (Invitrogen) along with protein ladders (SeeBlue™ Plus2 Pre-stained Protein Standard, Invitrogen). Samples were run at 150 kV, 400 mAmp for 30 minutes with NuPAGE MES SDS buffer (Invitrogen).

The gel was stained with SimplyBlue Safe Stain (Invitrogen) for 1 hour and de-stained in water overnight before imaging on a ChemiDoc Imager (Bio-Rad) or a GelDoc Go Imager (Bio-Rad). The obtained protein gel images were analyzed using the accompanying software Image Lab 6.1 for Mac (Bio-Rad).

### One-Pot PURE reaction assembly and data acquisition

Detailed methods describing the preparation of One-Pot PURE protein mixture and energy solution are provided in **Supplementary Information II**.

All One-Pot reactions were assembled on ice using low protein-binding microcentrifuge tubes (Eppendorf). Each reaction is 5 *μ*L in total volume and consists of 2 *μ*M *E. coli* Ribosome (NEB), 5 nM reporter protein expression plasmids (either P_T7_-MGA-UTR1-deGFP or P_T7_-UTR1-deGFP), 2.5x Energy Solution (125 mM HEPES-KOH pH 7.6, 250 mM Potassium L-Glutamate, 5 mM ATP and GTP at pH 7.5, 2.5 mM UTP and CTP at pH 7.5, either 56 A_260_U/mL for *E. coli* MRE600 tRNA or 75 – 200 A_260_U/mL for *E. coli* W tRNA, 50 mM Creatine Phosphate, 2.5 mM TCEP, 50 *μ*M Folinic Acid, 5 mM Spermidine, and 0.75 mM amino acids mixture at pH 7.5), 1.75 – 5 mg/mL of One-Pot PURE proteins and 0.5 – 1 *μ*M T7 RNAP (only for Single Strain One-Pot). For reactions using malachite green aptamer (MGA) fluorescence as a proxy for mRNA transcripts, 10 *μ*M of malachite green dye was added to the reaction.

Following assembly, 5 *μ*L of each reaction was pipetted onto a low-volume 384-well microplate with a non-binding surface (Corning), and the reaction trajectory was measured every 5 minutes using the BioTek Synergy Plate Reader at 37 °C. Fluorescence for deGFP was measured at 485/515 nm excitation/emission wavelengths at Gain 61, and fluorescence for malachite green aptamer was measured at 610/650 nm excitation/emission wavelengths at Gain 150. The fluorescence data for MGA and deGFP were calibrated to protein concentration data using synthesized MGA RNA and deGFP protein standards, provided in **Supplementary Information III, Figure SI.III.10**.

### Acquisition and analysis of proteomics data

A brief summary of LC-MS/MS workflow is provided below. Detailed methods are provided in

## Supporting information

Supplementary Information I. DNA Sequences

Supplementary Inforation III. Supplementary Figures

## Supplementary Information II

LC-MS/MS analyses of the digested peptides were performed on an EASY-nLC 1200 (Thermo Fisher Scientific) coupled to a Q Exactive HF hybrid quadrupole-Orbitrap mass spectrometer (Thermo Fisher Scientific). MS2 fragmentation spectra were searched with Proteome Discoverer SEQUEST (version 2.5; Thermo Scientific, Waltham, MA) against an *in silico* tryptic digested *E. coli* BL21 (DE3) and *E. coli* M15 database. The relative abundance of parental peptides was calculated by integrating the area under the curve of the MS1 peaks.

The proteomics data presented in this manuscript are from the same batch of LC-MS/MS run for results to be comparable. The relative protein abundances for each sample are first normalized by scaling the total sample protein abundance with the average total protein abundance for all samples. The proteins involved in the PURE system were extracted from the entire proteome dataset and scaled by their composition in PURE reactions (manufacturer’s recommended compositions for PURE_A_ and PURE_B_, 4 mg/mL for Dual Strain One-Pot PURE batches A and B in **Figure 2**, and 2.5 mg/mL for all Dual- and Single Strain One-Pot PURE batches A and B in **Figure 5**)

### Characterization of protease activity in M15/pREP4 and BL21/pREP4 strain

Plasmid expressing T7 RNAP (Addgene# 124128) is co-transformed into M15 and BL21 cells with pREP4 plasmid coding for constitutive LacI expression. Transformants were grown in a 5 mL overnight culture with LB with antibiotics and 1% glucose. 1 mL of the overnight was used to start the 150 mL final culture with LB and antibiotics. When culture OD exceeded 0.8, 1 mM IPTG was used to induce T7 RNAP expression, and the culture was grown for an additional 3 hours. At harvest, the cultures were divided into 50 mL aliquots, centrifuged at 5000 x g for 20 minutes to obtain cell pellets, and the pellets were stored at -80 °C until use. One 100 *μ*L aliquot of the M15 and BL21 cultures were saved as un-lysed controls.

To characterize M15 proteolysis during protein purification, each of the cell pellets was resuspended in 1 mL of the Lysis Buffer (50 mM HEPES-KOH pH 7.6, 100 mM NH_4_Cl, 500 mM NaCl, 10 mM MgCl_2_, and 1 mM TCEP). Two variations of the lysis buffer were prepared: one Pepstatin A (1 *μ*g/mL), and another with cOmplete™ Mini, EDTA-free protease inhibitor at the recommended working concentration. The resuspended cells were lysed using the freeze-thaw method by alternating between flash freezing in liquid nitrogen and thawing in a room-temperature water bath 8 times. The lysate was centrifuged at 15,000 rpm at 4°C for 20 minutes to remove cellular debris before His-tag purification.

A 2 mL column (Thermo Scientific) was prepared with 200 *μ*L of Ni-Sepharose resin (Cytiva) and equilibrated using 15 mL of Wash Buffer (50 mM HEPES-KOH pH 7.6, 100 mM NH_4_Cl, 500 mM NaCl, 10 mM MgCl_2_, 1 mM TCEP, and 20 mM Imidazole-HCl pH 7). Following equilibration, the lysate-resin mixture was rocked at 4 °C for 1 hour to enhance protein binding to the resin, washed with 20 mL of Wash Buffer, and eluted with 250 *μ*L of Elution Buffer (50 mM HEPES-KOH pH 7.6, 100 mM NH_4_Cl, 500 mM NaCl, 10 mM MgCl_2_, 1 mM TCEP, and 500 mM Imidazole-HCl pH 7). The un-lysed cell culture aliquots, as well as purified proteins from M15 and BL21 strains, were then prepared for SDS-PAGE analysis using the same setting described in the previous sections.

## AUTHOR CONTRIBUTIONS

*Conceptualization:* YZ, RMM, PSF, TC.; *Investigation:* YZ, MD, YQ, LB, ZAM; *Analysis:* YZ, MD, YQ, LB, ZAM; *Writing – manuscript:* YZ, MD, LB, ZAM; *Writing – review and editing:* YZ, MD, RMM, PSF, LB, ZAM, YQ, TC; *Visualization:* YZ, MD, LB, ZAM; *Supervision:* RMM, PSF, TC.

## ACKNOWLEDGEMENT

We sincerely thank the Murray Lab members, particularly Dr. Zoila Jurado and Miryong (Miki) Yun, who contributed to the project’s early setup and discussions. We also thank b.next, a company working on open-source synthetic cell protocols (https://bnext.bio), for insightful discussions on the 36-Pot PURE preparation. We also thank Dr. Jurado, Manisha Kapasiawala, and Dr. John Marken for their review and feedback on this manuscript.

Dr. Jurado prepared the P_T7_-MGA-UTR1-deGFP plasmid ^47^ used in this work. We thank Prof. Lulu Qian for access to the ChemiDoc Imager used for protein gel image acquisition. We thank the Caltech Protein Expression Center (PEC) for preparing the purified T7 RNAP used in this work.

