## Supplementary Information I. DNA Sequences for "Optimizing protein production in the One-Pot Pure system: insights into reaction composition and expression efficiency"

#### *Dual Strain One-Pot PURE plasmids*

**Table SI-I 1.** Addgene accession number for all proteins used in Dual Strain One-Pot PURE

| Protein | Vector | Expression Strain | Addgene Plasmid # |
|---|---|---|---|
| AlaRS | pQE30-His-AlaRS | M15 | 124103 |
| ArgRS | pET16b-His-ArgRS | BL21(DE3) | 124104 |
| AsnRS | pQE30-His-AsnRS | M15 | 124105 |
| AspRS | pET21a-AspRS-His | BL21(DE3) | 124106 |
| CysRS | pET21a-CysRS-His | BL21(DE3) | 124107 |
| GlnRS | pET21a-GlnRS-His | BL21(DE3) | 124108 |
| GluRS | pET21a-GluRS-His | BL21(DE3) | 124109 |
| GlyRS | pET21a-GlyRS-His | BL21(DE3) | 124110 |
| HisRS | pET21a-HisRS-His | BL21(DE3) | 124111 |
| IleRS | pET21a-His-IleRS | BL21(DE3) | 124112 |
| LeuRS | pET21a-LeuRS-His | BL21(DE3) | 124113 |
| LysRS | pET21a-LysRS-His | BL21(DE3) | 124114 |
| MetRS | pET21a-MetRS-His | BL21(DE3) | 124115 |
| PheRS | pQE30-His-PheRS | M15 | 124116 |
| ProRS | pET21a-ProRS-His | BL21(DE3) | 124117 |
| SerRS | pET21a-SerRS-His | BL21(DE3) | 124118 |
| ThrRS | pQE30-His-ThrRS | M15 | 124119 |
| TrpRS | pET21a-TrpRS-His | BL21(DE3) | 124120 |
| TyrRS | pET21a-TyrRS-His | BL21(DE3) | 124121 |
| ValRS | pET21a-ValRS-His | BL21(DE3) | 124122 |
| IF1 | pQE30-His-IF1 | M15 | 124123 |
| IF2 | pQE30-His-IF2 | M15 | 124124 |
| IF3 | pQE30-His-IF3 | M15 | 124125 |
| EF-G | pQE60-EFG-His | M15 | 124126 |
| EF-Tu | pQE60-EFTu-His | M15 | 124127 |
| EF-Ts | pQE60-EFTs-His | M15 | 124128 |
| RF1 | pET21a-RF1-His | BL21(DE3) | 224372 |
| RF2 | pET15b-His-RF2 | BL21(DE3) | 124130 |
| RF3 | pQE30-His-RF3 | M15 | 124131 |
| RRF | pQE60-RRF-His | M15 | 124132 |
| MTF | pET21a-MTF-His | BL21(DE3) | 124133 |
| CKM | pQE30-His-CKM | M15 | 124134 |
| AK1 | pET29b-AK1-His | BL21(DE3) | 118977 |
| NDK | pQE30-His-NDK | M15 | 124136 |
| IPP | pET29b-IPP1-His | BL21(DE3) | 118978 |
| T7 RNAP | pQE30-His-T7RNAP | M15 | 124138 |

#### *Single Strain One-Pot PURE plasmids*

**Table SI-I 2.** Addgene accession number for 36 proteins used in Single-Strain One-Pot PURE

| Protein | Vector | Expression Strain | Addgene Plasmid # |
|---|---|---|---|
| AlaRS | pET21a-His-AlaRS | BL21(DE3) | 224362 |
| ArgRS | pET16b-His-ArgRS | BL21(DE3) | 124104 |
| AsnRS | pET21a-His-AsnRS | BL21(DE3) | 224363 |
| AspRS | pET21a-AspRS-His | BL21(DE3) | 124106 |
| CysRS | pET21a-CysRS-His | BL21(DE3) | 124107 |
| GlnRS | pET21a-GlnRS-His | BL21(DE3) | 124108 |
| GluRS | pET21a-GluRS-His | BL21(DE3) | 124109 |
| GlyRS | pET21a-GlyRS-His | BL21(DE3) | 124110 |
| HisRS | pET21a-HisRS-His | BL21(DE3) | 124111 |
| IleRS | pET21a-His-IleRS | BL21(DE3) | 124112 |
| LeuRS | pET21a-LeuRS-His | BL21(DE3) | 124113 |
| LysRS | pET21a-LysRS-His | BL21(DE3) | 124114 |
| MetRS | pET21a-MetRS-His | BL21(DE3) | 124115 |
| PheRS | pET21a-His-PheRS | BL21(DE3) | 224364 |
| ProRS | pET21a-ProRS-His | BL21(DE3) | 124117 |
| SerRS | pET21a-SerRS-His | BL21(DE3) | 124118 |
| ThrRS | pET21a-His-ThrRS | BL21(DE3) | 224365 |
| TrpRS | pET21a-TrpRS-His | BL21(DE3) | 124120 |
| TyrRS | pET21a-TyrRS-His | BL21(DE3) | 124121 |
| ValRS | pET21a-ValRS-His | BL21(DE3) | 124122 |
| IF1 | pET21a-His-IF1 | BL21(DE3) | 224366 |
| IF2 | pET21a-His-IF2 | BL21(DE3) | 224367 |
| IF3 | pET21a-His-IF3 | BL21(DE3) | 224368 |
| EF-G | pET21a-EFG-His | BL21(DE3) | 224369 |
| EF-Tu | pET21a-EFTu-His | BL21(DE3) | 224370 |
| EF-Ts | pET21a-EFTs-His | BL21(DE3) | 224371 |
| RF1 | pET21a-RF1-His | BL21(DE3) | 224372 |
| RF2 | pET15b-His-RF2 | BL21(DE3) | 124130 |
| RF3 | pET21a-His-RF3 | BL21(DE3) | 224373 |
| RRF | pET21a-RRF-His | BL21(DE3) | 224374 |
| MTF | pET21a-MTF-His | BL21(DE3) | 124133 |
| CKM | pET21a-His-CKM | BL21(DE3) | 224375 |
| AK1 | pET29b-AK1-His | BL21(DE3) | 118977 |
| NDK | pET21a-His-NDK | BL21(DE3) | 224376 |
| IPP | pET29b-IPP1-His | BL21(DE3) | 118978 |
| T7 RNAP** | pQE30-His-T7RNAP | M15 | 124138 |

** In the single-strain one-pot, T7 RNAP is purified separately. To prevent proteolysis from M15, we recommend expressing T7 RNAP in a BL21 (non-DE3) strain deficient in Lon and OmpT proteases with a $LacI^q$ plasmid (pREP4 and others).

#### *Reporter plasmids*

**Table SI-I 3.** Addgene accession number for reporter plasmids used in this work

| Plasmid | *E. coli* Strain | Addgene Plasmid # |
|---|---|---|
| $P_{T7}$-MGA-UTR1-deGFP-$T_{T7}$ | NEB Turbo | * pending |
| $P_{T7}$-UTR1-deGFP-$T_{T7}$ | NEB Turbo | * pending |

#### *Sequences of calibration standards*

*Malachite Green RNA aptamer sequence*

```
rArCrUrGrGrArUrCrCrCrGrArCrUrGrGrCrGrArGrArGrCrCrArGrGrUrArArCrGrArArUrGrGrAr
UrCrCrArArU
```

*His6-deGFP sequence*

```
ATGCATCACCATCACCATCACGGATCCATGGAGCTTTTCACTGGCGTTGTTCCCATCCTGGTCGAGCTGGACGGCGA
CGTAAACGGCCACAAGTTCAGCGTGTCCGGCGAGGGCGAGGGCGATGCCACCTACGGCAAGCTGACCCTGAAGTTCA
TCTGCACCACCGGCAAGCTGCCCGTGCCCTGGCCCACCCTCGTGACCACCCTGACCTACGGCGTGCAGTGCTTCAGC
CGCTACCCCGACCACATGAAGCAGCACGACTTCTTCAAGTCCGCCATGCCCGAAGGCTACGTCCAGGAGCGCACCAT
CTTCTTCAAGGACGACGGCAACTACAAGACCCGCGCCGAGGTGAAGTTCGAGGGCGACACCCTGGTGAACCGCATCG
AGCTGAAGGGCATCGACTTCAAGGAGGACGGCAACATCCTGGGGCACAAGCTGGAGTACAACTACAACAGCCACAAC
GTCTATATCATGGCCGACAAGCAGAAGAACGGCATCAAGGTGAACTTCAAGATCCGCCACAACATCGAGGACGGCAG
CGTGCAGCTCGCCGACCACTACCAGCAGAACACCCCCATCGGCGACGGCCCCGTGCTGCTGCCCGACAACCACTACC
TGAGCACCCAGTCCGCCCTGAGCAAAGACCCCAACGAGAAGCGCGATCACATGGTCCTGCTGGAGTTCGTGACCGCC
GCCGGGATCTAA
```

### *Sequence of pET21a-His-AlaRS*

#### $P_{T7}$ promoter - His6-AlaRS - T7 terminator

taatacgactcactataggggaattgtgagcggataacaattcccctctagaaataattttgtttaactttaagaaggagatatacatATGAGAGGA
TCGCATCACCATCACCATCACGGATCCGCATGCGATGACGATGACAAAAGCAACAGCACCGCTGAGATCCGTCAGGCGTTTCTCGACTTTTTCCATA
GTAAGGGACATCAGGTAGTTGCCAGCAGCTCCCTGGTACCCCATAACGACCCAACTTTGTTGTTTACCAACGCCGGGATGAACCAGTTCAAGGATGT
GTTCCTTGGGCTCGACAAGCGTAATTATTCCCGCGCTACCACTTCCCAACGCTGCGTGCGTGCGGGTGGTAAACACAACGACCTGGAAAACGTCGGT
TACACCGCGCGTCACCATACCTTCTTCGAAATGCTGGGCAACTTCAGCTTCGGCGACTATTTCAAACACGATGCCATTCAGTTTGCATGGGAACTGC
TGACCAGCGAAAAATGGTTTGCCCTGCCGAAAGAGCGTCTGTGGGTTACCGTCTATGAAAGCGACGACGAAGCCTACGAAATCTGGGAAAAAGAAGT
AGGGATCCCGCGCGAACGTATTATTCGCATCGGCGATAACAAAGGTGCGCCATACGCATCTGACAACTTCTGGCAGATGGGTGACACTGGTCCGTGC
GGCCCGTGCACCGAAATCTTCTACGATCACGGCGACCACATTTGGGGGGGCCCTCCGGGAAGCCCGGAAGAAGACGGCGACCGCTACATTGAGATCT
GGAACATCGTCTTCATGCAGTTCAACCGCCAGGCCGATGGCACGATGGAACCGCTGCCGAAGCCGTCTGTAGATACCGGTATGGGTCTGGAGCGTAT
TGCTGCGGTGCTGCAACACGTTAACTCTAACTATGACATCGACCTGTTCCGCACGCTGATCCAGGCGGTAGCGAAAGTCACTGGCGCAACCGATCTG
AGCAATAAATCGCTGCGCGTAATCGCTGACCACATTCGTTCTTGTGCGTTCCTAATCGCGGATGGCGTAATGCCGTCCAATGAAAACCGTGGTTATG
TACTGCGTCGTATCATTCGTCGCGCAGTGCGTCACGGTAATATGCTCGGCGCGAAAGAAACCTTCTTCTACAAACTGGTTGGTCCGCTGATCGACGT
TATGGGCTCTGCGGGTGAAGACCTGAAACGCCAGCAGGCGCAGGTTGAGCAGGTGCTGAAGACTGAAGAAGAGCAGTTTGCTCGTACTCTGGAGCGC
GGTCTGGCGTTGCTGGATGAAGAGCTGGCAAAACTTTCTGGTGATACGCTGGATGGTGAAACTGCTTTCCGTCTGTACGACACCTATGGCTTCCCGG
TTGACCTGACGGCTGATGTTTGTCGTGAGCGCAACATCAAAGTTGACGAAGCTGGTTTTGAAGCTGCAATGGAAGAGCAGCGTCGTCGCGCGCGCGA
AGCCAGCGGCTTTGGTGCCGATTACAACGCAATGATCCGTGTTGACAGTGCATCTGAATTTAAAGGCTATGACCATCTGGAACTGAACGGCAAAGTG
ACTGCGCTGTTTGTTGATGGTAAAGCGGTTGATGCCATCAATGCAGGCCAGGAAGCTGTGGTCGTGCTGGATCAAACGCCATTCTATGCGGAATCCG
GCGGTCAGGTTGGCGATAAAGGCGAACTGAAAGGCGCTAACTTCTCCTTTGCGGTGGAAGATACGCAGAAATACGGCCAGGCGATTGGTCACATCGG
TAAACTTGCTGCGGGTTCTCTGAAAGTGGGCGACGCGGTGCAGGCTGATGTTGATGAGGCTCGTCGCGCCCGTATTCGTCTGAATCACTCCGCAACG
CACCTGATGCACGCTGCGCTGCGCCAGGTTCTGGGTACTCATGTATCGCAGAAAGGTTCACTGGTTAACGACAAGGTGCTGCGCTTCGACTTCTCAC
ACAACGAAGCGATGAAACCAGAAGAGATTCGTGCGGTCGAAGACCTGGTGAACACACAGATTCGTCGCAATTTGCCGATCGAAACCAACATCATGGA
TCTCGAAGCGGCGAAAGCGAAAGGTGCGATGGCGCTGTTCGGCGAGAAGTATGATGAGCGCGTACGCGTGCTGAGCATGGGCGATTTCTCTACCGAG
TTGTGTGGCGGTACTCACGCCAGCCGCACTGGTGATATTGGTCTGTTCCGCATCATCTCTGAATCGGGTACTGCTGCAGGCGTTCGTCGTATCGAAG
CGGTAACCGGAGAAGGTGCTATCGCCACCGTTCATGCAGACAGCGATCGCTTAAGCGAAGTCGCGCATCTGCTGAAAGGCGATAGCAATAATCTGGC
TGATAAAGTGCGCTCAGTACTGGAACGTACGCGTCAGCTGGAAAAAGAGTTACAACAGCTTAAAGAACAAGCTGCCGCACAGGAGAGCGCAAATCTT
TCCAGTAAGGCAATTGATGTTAATGGTGTTAAGCTGTTGGTTAGCGAGCTTAGCGGTGTTGAGCCGAAAATGTTGCGTACCATGGTTGACGATTTAA
AAAATCAGCTGGGGTCGACAATTATCGTGCTGGCAACGGTAGTCGAAGGTAAGGTTTCTCTGATTGCAGGCGTATCTAAGGACGTCACAGATCGTGT
GAAAGCAGGGGAACTGATTGGTATGGTCGCTCAGCAGGTGGGCGGCAAGGGTGGTGGACGTCCTGACATGGCGCAAGCCGGTGGTACGGATGCTGCG
GCCTTACCTGCAGCGTTAGCCAGTGTGAAAGGCTGGGTCAGCGCGAAATTGCAATAATGAgatccggctgctaacaaagcccgaaaggaagctgagt
tggctgctgccaccgctgagcaataactagcataaccccttggggcctctaaacgggtcttgaggggttttttgctgaaaggaggaactatatccgg
attggcgaatgggacgcgccctgtagcggcgcattaagcgcggcgggtgtggtggttacgcgcagcgtgaccgctacacttgccagcgccctagcgc
ccgctcctttcgctttcttcccttcctttctcgccacgttcgccggctttccccgtcaagctctaaatcgggggctccctttagggttccgatttag
tgctttacggcacctcgaccccaaaaaacttgattagggtgatggttcacgtagtgggccatcgccctgatagacggtttttcgccctttgacgttg
gagtccacgttctttaatagtggactcttgttccaaactggaacaacactcaaccctatctcggtctattcttttgatttataagggattttgccga
tttcggcctattggttaaaaaatgagctgatttaacaaaaatttaacgcgaattttaacaaaatattaacgcttacaatttaggtggcacttttcgg
ggaaatgtgcgcggaacccctatttgtttatttttctaaatacattcaaatatgtatccgctcatgagacaataaccctgataaatgcttcaataat
attgaaaaaggaagagtatgagtattcaacatttccgtgtcgcccttattcccttttttgcggcattttgccttcctgtttttgctcacccagaaac
gctggtgaaagtaaaagatgctgaagatcagttgggtgcacgagtgggttacatcgaactggatctcaacagcggtaagatccttgagagttttcgc
cccgaagaacgttttccaatgatgagcacttttaaagttctgctatgtggcgcggtattatcccgtattgacgccgggcaagagcaactcggtcgcc
gcatacactattctcagaatgacttggttgagtactcaccagtcacagaaaagcatcttacggatggcatgacagtaagagaattatgcagtgctgc
cataaccatgagtgataacactgcggccaacttacttctgacaacgatcggaggaccgaaggagctaaccgcttttttgcacaacatgggggatcat
gtaactcgccttgatcgttgggaaccggagctgaatgaagccataccaaacgacgagcgtgacaccacgatgcctgcagcaatggcaacaacgttgc
gcaaactattaactggcgaactacttactctagcttcccggcaacaattaatagactggatggaggcggataaagttgcaggaccacttctgcgctc
ggcccttccggctggctggtttattgctgataaatctggagccggtgagcgtgggtctcgcggtatcattgcagcactggggccagatggtaagccc
tcccgtatcgtagttatctacacgacggggagtcaggcaactatggatgaacgaaatagacagatcgctgagataggtgcctcactgattaagcatt
ggtaactgtcagaccaagtttactcatatatactttagattgatttaaaacttcatttttaatttaaaaggatctaggtgaagatcctttttgataa
tctcatgaccaaaatcccttaacgtgagttttcgttccactgagcgtcagaccccgtagaaaagatcaaaggatcttcttgagatcctttttttctg
cgcgtaatctgctgcttgcaaacaaaaaaaccaccgctaccagcggtggtttgtttgccggatcaagagctaccaactctttttccgaaggtaactg
gcttcagcagagcgcagataccaaatactgtccttctagtgtagccgtagttaggccaccacttcaagaactctgtagcaccgcctacatacctcgc
tctgctaatcctgttaccagtggctgctgccagtggcgataagtcgtgtcttaccgggttggactcaagacgatagttaccggataaggcgcagcgg
tcgggctgaacggggggttcgtgcacacagcccagcttggagcgaacgacctacaccgaactgagatacctacagcgtgagctatgagaaagcgcca
cgcttcccgaagggagaaaggcggacaggtatccggtaagcggcagggtcggaacaggagagcgcacgagggagcttccagggggaaacgcctggta
tctttatagtcctgtcgggtttcgccacctctgacttgagcgtcgatttttgtgatgctcgtcaggggggcggagcctatggaaaaacgccagcaac
gcggcctttttacggttcctggccttttgctggccttttgctcacatgttctttcctgcgttatcccctgattctgtggataaccgtattaccgcct
ttgagtgagctgataccgctcgccgcagccgaacgaccgagcgcagcgagtcagtgagcgaggaagcggaagagcgcctgatgcggtattttctcct
tacgcatctgtgcggtatttcacaccgcaatggtgcactctcagtacaatctgctctgatgccgcatagttaagccagtatacactccgctatcgct
acgtgactgggtcatggctgcgccccgacacccgccaacacccgctgacgcgccctgacgggcttgtctgctcccggcatccgcttacagacaagct

gtgaccgtctccgggagctgcatgtgtcagaggttttcaccgtcatcaccgaaacgcgcgaggcagctgcggtaaagctcatcagcgtggtcgtgaa
gcgattcacagatgtctgcctgttcatccgcgtccagctcgttgagtttctccagaagcgttaatgtctggcttctgataaagcgggccatgttaag
ggcggtttttcctgtttggtcactgatgcctccgtgtaagggggatttctgttcatgggggtaatgataccgatgaaacgagagaggatgctcacg
atacgggttactgatgatgaacatgcccggttactggaacgttgtgagggtaaacaactggcggtatggatgcggcgggaccagagaaaaatcactc
agggtcaatgccagcgcttcgttaatacagatgtaggtgttccacagggtagccagcagcatcctgcgatgcagatccggaacataatggtgcaggg
cgctgacttccgcgtttccagactttacgaaacacggaaaccgaagaccattcatgttgttgctcaggtcgcagacgttttgcagcagcagtcgctt
cacgttcgctcgcgtatcggtgattcattctgctaaccagtaaggcaaccccgccagcctagccgggtcctcaacgacaggagcacgatcatgcgca
cccgtggggccgccatgccggcgataatggcctgcttctcgccgaaacgtttggtggcgggaccagtgacgaaggcttgagcgagggcgtgcaagat
tccgaataccgcaagcgacaggccgatcatcgtcgcgctccagcgaaagcggtcctcgccgaaaatgacccagagcgctgccggcacctgtcctacg
agttgcatgataaagaagacagtcataagtgcggcgacgatagtcatgccccgcgcccaccggaaggagctgactgggttgaaggctctcaagggca
tcggtcgagatcccggtgcctaatgagtgagctaacttacattaattgcgttgcgctcactgcccgctttccagtcgggaaacctgtcgtgccagct
gcattaatgaatcggccaacgcgcggggagaggcggtttgcgtattgggcgccagggtggtttttcttttcaccagtgagacgggcaacagctgatt
gcccttcaccgcctggccctgagagagttgcagcaagcggtccacgctggtttgccccagcaggcgaaaatcctgtttgatggtggttaacggcggg
atataacatgagctgtcttcggtatcgtcgtatcccactaccgagatatccgcaccaacgcgcagcccggactcggtaatggcgcgcattgcgccca
gcgccatctgatcgttggcaaccagcatcgcagtgggaacgatgccctcattcagcatttgcatggtttgttgaaaaccggacatggcactccagtc
gccttcccgttccgctatcggctgaatttgattgcgagtgagatatttatgccagccagccagacgcagacgcgccgagacagaacttaatgggccc
gctaacagcgcgatttgctggtgacccaatgcgaccagatgctccacgcccagtcgcgtaccgtcttcatgggagaaaataatactgttgatgggtg
tctggtcagagacatcaagaaataacgccggaacattagtgcaggcagcttccacagcaatggcatcctggtcatccagcggatagttaatgatcag
cccactgacgcgttgcgcgagaagattgtgcaccgccgctttacaggcttcgacgccgcttcgttctaccatcgacaccaccacgctggcacccagt
tgatcggcgcgagatttaatcgccgcgacaatttgcgacggcgcgtgcagggccagactggaggtggcaacgccaatcagcaacgactgtttgcccg
ccagttgttgtgccacgcggttgggaatgtaattcagctccgccatcgccgcttccactttttcccgcgttttcgcagaaacgtggctggcctggtt
caccacgcgggaaacggtctgataagagacaccggcatactctgcgacatcgtataacgttactggtttcacattcaccaccctgaattgactctct
tccgggcgctatcatgccataccgcgaaaggttttgcgccattcgatggtgtccgggatctcgacgctctcccttatgcgactcctgcattaggaag
cagcccagtagtaggttgaggccgttgagcaccgccgccgcaaggaatggtgcatgcaaggagatggcgcccaacagtcccccggccacggggcctg
ccaccatacccacgccgaaacaagcgctcatgagcccgaagtggcgagcccgatcttccccatcggtgatgtcggcgatataggcgccagcaaccgc
acctgtggcgccggtgatgccggccacgatgcgtccggcgtagaggatcgagatctcgatcccgcgaaat

### *Sequence of pET21a-His-AsnRS*

#### $P_{T7}$ promoter - His6-AsnRS - T7 terminator

taatacgactcactataggggaattgtgagcggataacaattcccctctagaaataattttgtttaactttaagaaggagatatacatATGAGAGGA
TCGCATCACCATCACCATCACGGATCCATGAGCGTTGTGCCTGTAGCCGACGTACTCCAGGGCCGTGTAGCCGTTGACAGCGAAGTCACCGTGCGCG
GATGGGTACGTACCCGCCGAGATTCAAAAGCTGGCATCTCCTTCCTCGCCGTTTATGACGGTTCCTGCTTTGATCCTGTACAGGCTGTCATCAATAA
TTCTCTGCCCAATTACAATGAAGACGTCCTGCGTCTGACCACCGGCTGCTCGGTCATTGTGACGGGTAAAGTCGTGGCGTCGCCGGGCCAGGGGCAA
CAATTTGAAATTCAGGCCAGCAAGGTTGAAGTTGCTGGTTGGGTTGAAGATCCAGACACTTACCCGATGGCGGCAAAACGCCACAGCATTGAGTATC
TGCGTGAAGTCGCTCACCTGCGTCCGCGCACAAACCTGATTGGTGCCGTCGCGCGCGTTCGCCATACGCTGGCGCAGGCGCTGCATCGCTTCTTTAA
CGAGCAGGGATTCTTCTGGGTTTCAACGCCACTGATTACCGCATCTGATACCGAAGGTGCAGGCGAAATGTTCCGCGTTTCTACGCTGGATCTGGAA
AACCTGCCGCGTAACGATCAGGGCAAAGTGGATTTCGACAAAGACTTCTTTGGTAAAGAGTCTTTCCTGACCGTATCTGGCCAGTTGAACGGCGAAA
CCTACGCTTGCGCATTGTCCAAAATTTATACCTTCGGCCCGACTTTCCGTGCTGAAAACTCCAACACCAGCCGTCACCTGGCGGAATTCTGGATGCT
GGAGCCGGAAGTGGCGTTTGCTAACCTGAACGATATTGCGGGTCTGGCTGAAGCCATGCTGAAATATGTCTTCAAAGCGGTTCTCGAAGAACGCGCT
GACGACATGAAATTCTTCGCTGAACGCGTAGATAAAGATGCCGTTTCACGTCTGGAACGCTTCATTGAAGCCGATTTTGCGCAGGTGGATTATACCG
ACGCAGTGACCATTCTCGAAAACTGCGGCAGGAAGTTTGAAAACCCGGTTTACTGGGGAGTCGATCTCTCTTCTGAGCATGAGCGTTATCTGGCGGA
AGAACACTTTAAAGCACCGGTAGTGGTTAAAAACTATCCGAAAGATATTAAAGCGTTCTATATGCGCCTTAACGAAGACGGTAAAACCGTTGCGGCT
ATGGACGTTCTGGCTCCGGGCATCGGTGAGATCATTGGTGGCTCCCAGCGTGAAGAACGTCTGGACGTGCTGGACGAGCGTATGCTGGAAATGGGCC
TGAATAAAGAAGATTACTGGTGGTATCGCGATCTGCGTCGCTACGGTACTGTTCCGCATTCAGGTTTCGGTCTTGGTTTTGAACGTCTGATTGCTTA
CGTAACTGGCGTGCAAAACGTACGTGATGTGATTCCGTTCCCACGTACTCCGCGTAACGCCAGCTTCTAATGAgatccggctgctaacaaagcccga
aaggaagctgagttggctgctgccaccgctgagcaataactagcataaccccttggggcctctaaacgggtcttgaggggttttttgctgaaaggag
gaactatatccggattggcgaatgggacgcgccctgtagcggcgcattaagcgcggcgggtgtggtggttacgcgcagcgtgaccgctacacttgcc
agcgccctagcgcccgctcctttcgctttcttcccttcctttctcgccacgttcgccggctttccccgtcaagctctaaatcgggggctccctttag
ggttccgatttagtgctttacggcacctcgaccccaaaaaacttgattagggtgatggttcacgtagtgggccatcgccctgatagacggtttttcg
ccctttgacgttggagtccacgttctttaatagtggactcttgttccaaactggaacaacactcaaccctatctcggtctattcttttgatttataa
gggattttgccgatttcggcctattggttaaaaaatgagctgatttaacaaaaatttaacgcgaattttaacaaaatattaacgcttacaatttagg
tggcacttttcggggaaatgtgcgcggaacccctatttgtttatttttctaaatacattcaaatatgtatccgctcatgagacaataaccctgataa
atgcttcaataatattgaaaaaggaagagtatgagtattcaacatttccgtgtcgcccttattcccttttttgcggcattttgccttcctgtttttg
ctcacccagaaacgctggtgaaagtaaaagatgctgaagatcagttgggtgcacgagtgggttacatcgaactggatctcaacagcggtaagatcct
tgagagttttcgccccgaagaacgttttccaatgatgagcacttttaaagttctgctatgtggcgcggtattatcccgtattgacgccgggcaagag
caactcggtcgccgcatacactattctcagaatgacttggttgagtactcaccagtcacagaaaagcatcttacggatggcatgacagtaagagaat
tatgcagtgctgccataaccatgagtgataacactgcggccaacttacttctgacaacgatcggaggaccgaaggagctaaccgcttttttgcacaa
catgggggatcatgtaactcgccttgatcgttgggaaccggagctgaatgaagccataccaaacgacgagcgtgacaccacgatgcctgcagcaatg
gcaacaacgttgcgcaaactattaactggcgaactacttactctagcttcccggcaacaattaatagactggatggaggcggataaagttgcaggac
cacttctgcgctcggcccttccggctggctggtttattgctgataaatctggagccggtgagcgtgggtctcgcggtatcattgcagcactggggcc
agatggtaagccctcccgtatcgtagttatctacacgacggggagtcaggcaactatggatgaacgaaatagacagatcgctgagataggtgcctca
ctgattaagcattggtaactgtcagaccaagtttactcatatatactttagattgatttaaaacttcatttttaatttaaaaggatctaggtgaaga
tcctttttgataatctcatgaccaaaatcccttaacgtgagttttcgttccactgagcgtcagaccccgtagaaaagatcaaaggatcttcttgaga
tcctttttttctgcgcgtaatctgctgcttgcaaacaaaaaaaccaccgctaccagcggtggtttgtttgccggatcaagagctaccaactcttttt
ccgaaggtaactggcttcagcagagcgcagataccaaatactgtccttctagtgtagccgtagttaggccaccacttcaagaactctgtagcaccgc
ctacatacctcgctctgctaatcctgttaccagtggctgctgccagtggcgataagtcgtgtcttaccgggttggactcaagacgatagttaccgga
taaggcgcagcggtcgggctgaacggggggttcgtgcacacagcccagcttggagcgaacgacctacaccgaactgagatacctacagcgtgagcta
tgagaaagcgccacgcttcccgaagggagaaaggcggacaggtatccggtaagcggcagggtcggaacaggagagcgcacgagggagcttccagggg
gaaacgcctggtatctttatagtcctgtcgggtttcgccacctctgacttgagcgtcgatttttgtgatgctcgtcaggggggcggagcctatggaa
aaacgccagcaacgcggcctttttacggttcctggccttttgctggccttttgctcacatgttctttcctgcgttatcccctgattctgtggataac
cgtattaccgcctttgagtgagctgataccgctcgccgcagccgaacgaccgagcgcagcgagtcagtgagcgaggaagcggaagagcgcctgatgc
ggtattttctccttacgcatctgtgcggtatttcacaccgcaatggtgcactctcagtacaatctgctctgatgccgcatagttaagccagtataca
ctccgctatcgctacgtgactgggtcatggctgcgccccgacacccgccaacacccgctgacgcgccctgacgggcttgtctgctcccggcatccgc
ttacagacaagctgtgaccgtctccgggagctgcatgtgtcagaggttttcaccgtcatcaccgaaacgcgcgaggcagctgcggtaaagctcatca
gcgtggtcgtgaagcgattcacagatgtctgcctgttcatccgcgtccagctcgttgagtttctccagaagcgttaatgtctggcttctgataaagc
gggccatgttaagggcggttttttcctgtttggtcactgatgcctccgtgtaagggggatttctgttcatgggggtaatgataccgatgaaacgaga
gaggatgctcacgatacgggttactgatgatgaacatgcccggttactggaacgttgtgagggtaaacaactggcggtatggatgcggcgggaccag
agaaaaatcactcagggtcaatgccagcgcttcgttaatacagatgtaggtgttccacagggtagccagcagcatcctgcgatgcagatccggaaca
taatggtgcagggcgctgacttccgcgtttccagactttacgaaacacggaaaccgaagaccattcatgttgttgctcaggtcgcagacgttttgca
gcagcagtcgcttcacgttcgctcgcgtatcggtgattcattctgctaaccagtaaggcaaccccgccagcctagccgggtcctcaacgacaggagc
acgatcatgcgcacccgtgggccgccatgccggcgataatggcctgcttctcgccgaaacgtttggtggcgggaccagtgacgaaggcttgagcga
gggcgtgcaagattccgaataccgcaagcgacaggccgatcatcgtcgcgctccagcgaaagcggtcctcgccgaaaatgacccagagcgctgccgg
cacctgtcctacgagttgcatgataaagaagacagtcataagtgcggcgacgatagtcatgccccgcgcccaccggaaggagctgactgggttgaag
gctctcaagggcatcggtcgagatcccggtgcctaatgagtgagctaacttacattaattgcgttgcgctcactgcccgctttccagtcgggaaacc
tgtcgtgccagctgcattaatgaatcggccaacgcgcggggagaggcggtttgcgtattgggcgccagggtggtttttcttttcaccagtgagacgg
gcaacagctgattgcccttcaccgcctggccctgagagagttgcagcaagcggtccacgctggtttgccccagcaggcgaaaatcctgtttgatggt

ggttaacggcgggatataacatgagctgtcttcggtatcgtcgtatcccactaccgagatatccgcaccaacgcgcagcccggactcggtaatggcg
cgcattgcgcccagcgccatctgatcgttggcaaccagcatcgcagtgggaacgatgccctcattcagcatttgcatggtttgttgaaaaccggaca
tggcactccagtcgccttcccgttccgctatcggctgaatttgattgcgagtgagatatttatgccagccagccagacgcagacgcgccgagacaga
acttaatgggcccgctaacagcgcgatttgctggtgacccaatgcgaccagatgctccacgcccagtcgcgtaccgtcttcatgggagaaaataata
ctgttgatgggtgtctggtcagagacatcaagaaataacgccggaacattagtgcaggcagcttccacagcaatggcatcctggtcatccagcggat
agttaatgatcagcccactgacgcgttgcgcgagaagattgtgcaccgccgctttacaggcttcgacgccgcttcgttctaccatcgacaccaccac
gctggcacccagttgatcggcgcgagatttaatcgccgcgacaatttgcgacggcgcgtgcagggccagactggaggtggcaacgccaatcagcaac
gactgtttgcccgccagttgttgtgccacgcggttgggaatgtaattcagctccgccatcgccgcttccactttttcccgcgttttcgcagaaacgt
ggctggcctggttcaccacgcgggaaacggtctgataagagacaccggcatactctgcgacatcgtataacgttactggtttcacattcaccaccct
gaattgactctcttccgggcgctatcatgccataccgcgaaaggttttgcgccattcgatggtgtccgggatctcgacgctctcccttatgcgactc
ctgcattaggaagcagcccagtagtaggttgaggccgttgagcaccgccgccgcaaggaatggtgcatgcaaggagatggcgcccaacagtcccccg
gccacggggcctgccaccatacccacgccgaaacaagcgctcatgagcccgaagtggcgagcccgatcttccccatcggtgatgtcggcgatatagg
cgccagcaaccgcacctgtggcgccggtgatgccggccacgatgcgtccggcgtagaggatcgagatctcgatcccgcgaaat

### *Sequence of pET21a-His-PheRS*

#### $P_{T7}$ promoter - His6-PheRSa-PheRSb - T7 terminator

```
taatacgactcactataggggaattgtgagcggataacaattcccctctagaaataattttgtttaactttaagaaggagatatacatATGAGAGGA
TCGCATCACCATCACCATCACGGATCCGCATGCGATGACGATGACAAATCACATCTCGCAGAACTGGTTGCCAGTGCGAAGGCGGCCATTAGCCAGG
CGTCAGATGTTGCCGCGTTAGATAATGTGCGCGTCGAATATTTGGGTAAAAAGGGCACTTAACCCTTCAGATGACGACCCTGCGTGAGCTGCCGCC
AGAAGAGCGTCCGGCAGCTGGTGCGGTTATCAACGAAGCGAAAGAGCAGGTTCAGCAGGCGCTGAATGCGCGTAAAGCGGAACTGGAAAGCGCTGCA
CTGAATGCGCGTCTGGCGGCGGAAACGATTGATGTCTCTCTGCCAGGTCGTCGCATTGAAAACGGCGGTCTGCATCCGGTTACCCGTACCATCGACC
GTATCGAAAGTTTCTTCGGTGAGCTTGGCTTTACCGTGGCAACCGGGCCGGAAATCGAAGACGATTATCATAACTTCGATGCTCTGAACATTCCTGG
TCACCACCCGGCGCGCGCTGACCACGACACTTTCTGGTTTGACACTACCCGCCTGCTGCGTACCCAGACCTCTGGCGTACAGATCCGCACCATGAAA
GCCCAGCAGCCACCGATTCGTATCATCGCGCCTGGCCGTGTTTATCGTAACGACTACGACCAGACTCACACGCCGATGTTCCATCAGATGGAAGGTC
TGATTGTTGATACCAACATCAGCTTTACCAACCTGAAAGGCACGCTGCACGACTTCCTGCGTAACTTCTTTGAGGAAGATTTGCAGATTCGCTTCCG
TCCTTCCTACTTCCCGTTTACCGAACCTTCTGCAGAAGTGGACGTCATGGGTAAAAACGGTAAATGGCTGGAAGTGCTGGGCTGCGGGATGGTGCAT
CCGAACGTGTTGCGTAACGTTGGCATCGACCCGGAAGTTTACTCTGGTTTCGCCTTCGGGATGGGGATGGAGCGTCTGACTATGTTGCGTTACGGCG
TCACCGACCTGCGTTCATTCTTCGAAAACGATCTGCGTTTCCTCAAACAGTTTAAATAAGGCAGGAATAGATTATGAAATTCGGTGAACTGTGGTTA
CGCGAATGGGTGAACCCGGCGATTGATAGCGATGCGCTGGCAAATCAAATCACTATGGCGGGCCTGGAAGTTGACGGTGTAGAACCGGTTGCCGGCA
GCTTCCACGGCGTGGTCGTTGGTGAAGTGGTTGAGTGTGCGCAGCATCCGAACGCTGACAAACTGCGTGTGACAAAAGTGAATGTCGGCGGCGATCG
CCTGCTGGACATCGTCTGCGGTGCGCCAAACTGCCGTCAGGGCCTGCGTGTAGCGGTAGCGACCATTGGTGCTGTTCTGCCGGGTGATTTCAAAATT
AAAGCGGCGAAACTGCGTGGCGAACCGTCTGAAGGGATGCTGTGCTCCTTCTCTGAACTGGGCATTTCTGACGATCACAGCGGCATTATCGAACTGC
CTGCGGATGCGCCGATTGGCACCGATATCCGTGAATACCTGAAACTTGATGACAACACCATCGAAATCAGCGTGACGCCAAACCGTGCCGACTGCTT
AGGCATCATTGGTGTTGCGCGTGACGTTGCCGTGCTGAACCAGCTGCCGCTGGTTCAACCGGAAATCGTTCCGGTTGGTGCGACCATCGACGACACG
CTGCCGATTACAGTCGAAGCGCCGGAAGCCTGCCCGCGTTATCTTGGCCGTGTGGTAAAAGGCATTAACGTTAAAGCGCCAACTCCGCTGTGGATGA
AAGAAAAACTGCGTCGTTGCGGGATCCGTTCTATCGATGCAGTTGTTGACGTCACCAACTATGTGCTGCTCGAACTGGGCCAGCCGATGCACGCTTT
CGATAAAGATCGCATTGAAGGCGGCATTGTGGTGCGGATGGCGAAAGAGGGCGAAACGCTGGTGCTGCTCGACGGTACTGAAGCGAAGCTGAATGCT
GACACTCTGGTCATCGCCGACCACAACAAGGCGCTGGCGATGGGCGGCATCTTCGGTGGCGAACACTCTGGCGTGAATGACGAAACACAAAACGTGC
TGCTGGAATGCGCGTTCTTTAGCCCGCTGTCTATCACCGGTCGTGCTCGTCGTCATGGCCTGCATACCGATGCGTCTCACCGTTATGAGCGTGGCGT
TGATCCGGCACTGCAGCACAAAGCGATGGAACGTGCGACCCGTCTGCTGATCGACATCTGCGGTGGTGAGGCTGGCCCGGTAATTGATATCACCAAC
GAAGCAACGCTGCCGAAGCGTGCAACCATCACTCTACGTCGTAGCAAACTGGATCGCCTGATCGGCCATCATATTGCGGATGAGCAGGTAACTGACA
TTCTGCGTCGTCTCGGCTGCGAAGTGACCGAAGGCAAAGACGAATGGCAGGCAGTTGCGCCGAGCTGGCGTTTCGATATGGAGATTGAAGAAGATCT
GGTTGAAGAAGTCGCGCGTGTTTACGGCTACAACAACATCCCGGATGAGCCGGTACAGGCAAGCCTGATTATGGGTACTCACCGTGAAGCTGACCTG
TCGCTCAAGCGCGTGAAAACGCTGCTCAACGACAAAGGCTATCAGGAAGTGATCACCTACAGCTTCGTTGATCCGAAAGTGCAGCAGATGATCCATC
CAGGCGTTGAAGCCTTACTGCTGCCAAGCCCGATCTCTGTTGAAATGTCAGCAATGCGTCTTTCTCTGTGGACTGGCCTGCTGGCAACCGTGGTGTA
CAACCAGAACCGTCAGCAGAACCGTGTGCGCATTTTCGAAAGCGGTCTGCGTTTCGTACCAGATACTCAGGCACCGTTGGGCATTCGTCAGGATCTG
ATGTTAGCCGGTGTGATTTGCGGTAACCGTTACGAAGAGCACTGGAACCTGGCAAAAGAGACCGTTGATTTCTATGATTTGAAAGGCGATCTTGAAT
CCGTTCTCGACCTGACCGGTAAACTGAATGAGGTTGAGTTCCGTGCAGAAGCGAATCCGGCACTGCATCCGGGGCAATCCGCAGCGATTTATCTGAA
AGGTGAACGTATTGGTTTTGTTGGGGTTGTTCATCCTGAACTGGAACGTAAACTGGATCTTAACGGTCGCACTCTGGTGTTCGAACTGGAGTGGAAC
AAGCTCGCAGACCGCGTGGTGCCTCAGGCGCGCGAGATTTCTCGCTTCCCGGCGAACCGTCGTGACATCGCGGTGGTGGTCGCAGAAAACGTTCCCG
CAGCGGATATTTTATCCGAATGTAAGAAAGTTGGCGTAAATCAGGTAGTTGGCGTAAACTTATTTGACGTGTACCGCGGTAAGGGTGTTGCGGAGGG
GTATAAGAGCCTCGCCATAAGCCTGATCCTGCAAGATACCAGCCGTACACTCGAAGAAGAGGAGATTGCCGCTACCGTCGCCAAATGTGTAGAGGCA
TTAAAAGAGCGATTCCAGGCATCATTGAGGGATTGATGAGATCCGGCTGCTAACTGAgatccggctgctaacaaagcccgaaaggaagctgagttgg
ctgctgccaccgctgagcaataactagcataaccccttggggcctctaaacgggtcttgaggggttttttgctgaaaggaggaactatatccggatt
ggcgaatgggacgcgccctgtagcggcgcattaagcgcggcgggtgtggtggttacgcgcagcgtgaccgctacacttgccagcgccctagcgcccg
ctcctttcgctttcttcccttcctttctcgccacgttcgccggctttccccgtcaagctctaaatcgggggctccctttagggttccgatttagtgc
tttacggcacctcgaccccaaaaaacttgattagggtgatggttcacgtagtgggccatcgccctgatagacggtttttcgccctttgacgttggag
tccacgttctttaatagtggactcttgttccaaactggaacaacactcaaccctatctcggtctattcttttgatttataagggattttgccgattt
cggcctattggttaaaaaatgagctgatttaacaaaaatttaacgcgaattttaacaaaatattaacgcttacaatttaggtggcacttttcgggga
aatgtgcgcggaacccctatttgtttatttttctaaatacattcaaatatgtatccgctcatgagacaataaccctgataaatgcttcaataatatt
gaaaaaggaagagtatgagtattcaacatttccgtgtcgcccttattcccttttttgcggcattttgccttcctgtttttgctcacccagaaacgct
ggtgaaagtaaaagatgctgaagatcagttgggtgcacgagtgggttacatcgaactggatctcaacagcggtaagatccttgagagttttcgcccc
gaagaacgttttccaatgatgagcacttttaaagttctgctatgtggcgcggtattatcccgtattgacgccgggcaagagcaactcggtcgccgca
tacactattctcagaatgacttggttgagtactcaccagtcacagaaaagcatcttacggatggcatgacagtaagagaattatgcagtgctgccat
aaccatgagtgataacactgcggccaacttacttctgacaacgatcggaggaccgaaggagctaaccgcttttttgcacaacatgggggatcatgta
actcgccttgatcgttgggaaccggagctgaatgaagccataccaaacgacgagcgtgacaccacgatgcctgcagcaatggcaacaacgttgcgca
aactattaactggcgaactacttactctagcttcccggcaacaattaatagactggatggaggcggataaagttgcaggaccacttctgcgctcggc
ccttccggctggctggtttattgctgataaatctggagccggtgagcgtgggtctcgcggtatcattgcagcactggggccagatggtaagccctcc
cgtatcgtagttatctacacgacggggagtcaggcaactatggatgaacgaaatagacagatcgctgagataggtgcctcactgattaagcattggt
aactgtcagaccaagtttactcatatatactttagattgatttaaaacttcatttttaatttaaaaggatctaggtgaagatcctttttgataatct
catgaccaaaatcccttaacgtgagttttcgttccactgagcgtcagaccccgtagaaaagatcaaaggatcttcttgagatcctttttttctgcgc
gtaatctgctgcttgcaaacaaaaaaaccaccgctaccagcggtggtttgtttgccggatcaagagctaccaactctttttccgaaggtaactggct
tcagcagagcgcagataccaaatactgtccttctagtgtagccgtagttaggccaccacttcaagaactctgtagcaccgcctacatacctcgctct
```

gctaatcctgttaccagtggctgctgccagtggcgataagtcgtgtcttaccgggttggactcaagacgatagttaccggataaggcgcagcggtcg
ggctgaacggggggttcgtgcacacagcccagcttggagcgaacgacctacaccgaactgagatacctacagcgtgagctatgagaaagcgccacgc
ttcccgaagggagaaaggcggacaggtatccggtaagcggcagggtcggaacaggagagcgcacgagggagcttccagggggaaacgcctggtatct
ttatagtcctgtcgggtttcgccacctctgacttgagcgtcgatttttgtgatgctcgtcaggggggcggagcctatggaaaaacgccagcaacgcg
gcctttttacggttcctggccttttgctggccttttgctcacatgttctttcctgcgttatcccctgattctgtggataaccgtattaccgcctttg
agtgagctgataccgctcgccgcagccgaacgaccgagcgcagcgagtcagtgagcgaggaagcggaagagcgcctgatgcggtattttctccttac
gcatctgtgcggtatttcacaccgcaatggtgcactctcagtacaatctgctctgatgccgcatagttaagccagtatacactccgctatcgctacg
tgactgggtcatggctgcgccccgacacccgccaacacccgctgacgcgccctgacgggcttgtctgctcccggcatccgcttacagacaagctgtg
accgtctccgggagctgcatgtgtcagaggttttcaccgtcatcaccgaaacgcgcgaggcagctgcggtaaagctcatcagcgtggtcgtgaagcg
attcacagatgtctgcctgttcatccgcgtccagctcgttgagtttctccagaagcgttaatgtctggcttctgataaagcgggccatgttaagggc
ggttttttcctgtttggtcactgatgcctccgtgtaagggggatttctgttcatgggggtaatgataccgatgaaacgagagaggatgctcacgata
cgggttactgatgatgaacatgcccggttactggaacgttgtgagggtaaacaactggcggtatggatgcggcgggaccagagaaaaatcactcagg
gtcaatgccagcgcttcgttaatacagatgtaggtgttccacagggtagccagcagcatcctgcgatgcagatccggaacataatggtgcagggcgc
tgacttccgcgtttccagactttacgaaacacggaaaccgaagaccattcatgttgttgctcaggtcgcagacgttttgcagcagcagtcgcttcac
gttcgctcgcgtatcggtgattcattctgctaaccagtaaggcaaccccgccagcctagccgggtcctcaacgacaggagcacgatcatgcgcaccc
gtggggccgccatgccggcgataatggcctgcttctcgccgaaacgtttggtggcgggaccagtgacgaaggcttgagcgagggcgtgcaagattcc
gaataccgcaagcgacaggccgatcatcgtcgcgctccagcgaaagcggtcctcgccgaaaatgacccagagcgctgccggcacctgtcctacgagt
tgcatgataaagaagacagtcataagtgcggcgacgatagtcatgccccgcgcccaccggaaggagctgactgggttgaaggctctcaagggcatcg
gtcgagatcccggtgcctaatgagtgagctaacttacattaattgcgttgcgctcactgcccgctttccagtcgggaaacctgtcgtgccagctgca
ttaatgaatcggccaacgcgcggggagaggcggtttgcgtattgggcgccagggtggtttttcttttcaccagtgagacgggcaacagctgattgcc
cttcaccgcctggccctgagagagttgcagcaagcggtccacgctggtttgccccagcaggcgaaaatcctgtttgatggtggttaacggcgggata
taacatgagctgtcttcggtatcgtcgtatcccactaccgagatatccgcaccaacgcgcagcccggactcggtaatggcgcgcattgcgcccagcg
ccatctgatcgttggcaaccagcatcgcagtgggaacgatgccctcattcagcatttgcatggtttgttgaaaaccggacatggcactccagtcgcc
ttcccgttccgctatcggctgaatttgattgcgagtgagatatttatgccagccagccagacgcagacgcgccgagacagaacttaatgggcccgct
aacagcgcgatttgctggtgacccaatgcgaccagatgctccacgcccagtcgcgtaccgtcttcatgggagaaaataatactgttgatgggtgtct
ggtcagagacatcaagaaataacgccggaacattagtgcaggcagcttccacagcaatggcatcctggtcatccagcggatagttaatgatcagccc
actgacgcgttgcgcgagaagattgtgcaccgccgctttacaggcttcgacgccgcttcgttctaccatcgacaccaccacgctggcacccagttga
tcggcgcgagatttaatcgccgcgacaatttgcgacggcgcgtgcagggccagactggaggtggcaacgccaatcagcaacgactgtttgcccgcca
gttgttgtgccacgcggttgggaatgtaattcagctccgccatcgccgcttccactttttcccgcgttttcgcagaaacgtggctggcctggttcac
cacgcgggaaacggtctgataagagacaccggcatactctgcgacatcgtataacgttactggtttcacattcaccaccctgaattgactctcttcc
gggcgctatcatgccataccgcgaaaggttttgcgccattcgatggtgtccgggatctcgacgctctcccttatgcgactcctgcattaggaagcag
cccagtagtaggttgaggccgttgagcaccgccgccgcaaggaatggtgcatgcaaggagatggcgcccaacagtcccccggccacggggcctgcca
ccatacccacgccgaaacaagcgctcatgagcccgaagtggcgagcccgatcttccccatcggtgatgtcggcgatataggcgccagcaaccgcacc
tgtggcgccggtgatgccggccacgatgcgtccggcgtagaggatcgagatctcgatcccgcgaaat

### ***Sequence of pET21a-His-ThrRS***

#### $P_{T7}$ promoter - His6-ThrRS - T7 terminator

taatacgactcactataggggaattgtgagcggataacaattcccctctagaaataattttgtttaactttaagaaggagatatacatATGAGAGGA
TCGCATCACCATCACCATCACGGATCCGATGACGATGACAAACCTGTTATAACTCTTCCTGATGGCAGCCAACGCCATTACGATCACGCTGTAAGCC
CCATGGATGTTGCGCTGGACATTGGTCCAGGTCTGGCGAAAGCCTGTATCGCAGGGCGCGTTAATGGCGAACTGGTTGATGCTTGCGATCTGATTGA
AAACGACGCACAACTGTCGATCATTACCGCCAAAGACGAAGAAGGTCTGGAGATCATTCGTCACTCCTGTGCGCACCTGTTAGGGCACGCGATTAAA
CAACTTTGGCCGCATACCAAAATGGCAATCGGCCCGGTTATTGACAACGGTTTTTATTACGACGTTGATCTTGACCGCACGTTAACCCAGGAAGATG
TCGAAGCACTCGAGAAGCGGATGCATGAGCTTGCTGAGAAAAACTACGACGTCATTAAGAAGAAAGTCAGCTGGCACGAAGCGCGTGAAACTTTCGC
CAACCGTGGGGAGAGCTACAAAGTCTCCATTCTTGACGAAAACATCGCCCATGATGACAAGCCAGGTCTGTACTTCCATGAAGAATATGTCGATATG
TGCCGCGGTCCGCACGTACCGAACATGCGTTTCTGCCATCATTTCAAACTAATGAAAACGGCAGGGGCTTACTGGCGTGGCGACAGCAACAACAAAA
TGTTGCAACGTATTTACGGTACGGCGTGGGCAGACAAAAAAGCACTTAACGCTTACCTGCAGCGCCTGGAAGAAGCCGCGAAACGCGACCACCGTAA
AATCGGTAAACAGCTCGACCTGTACCATATGCAGGAAGAAGCGCCGGGTATGGTATTCTGGCACAACGACGGCTGGACCATCTTCCGTGAACTGGAA
GTGTTTGTTCGTTCTAAACTGAAAGAGTACCAGTATCAGGAAGTTAAAGGTCCGTTCATGATGGACCGTGTCCTGTGGGAAAAAACCGGTCACTGGG
ACAACTACAAAGATGCAATGTTCACCACATCTTCTGAGAACCGTGAATACTGCATTAAGCCGATGAACTGCCCGGGTCACGTACAAATTTTCAACCA
GGGGCTGAAGTCTTATCGCGATCTGCCGCTGCGTATGGCCGAGTTTGGTAGCTGCCACCGTAACGAGCCGTCAGGTTCGCTGCATGGCCTGATGCGC
GTGCGTGGATTTACCCAGGATGACGCGCATATCTTCTGTACTGAAGAACAAATTCGCGATGAAGTTAACGGATGTATCCGTTTAGTCTATGATATGT
ACAGCACTTTTGGCTTCGAGAAGATCGTCGTCAAACTCTCCACTCGTCCTGAAAAACGTATTGGCAGCGACGAAATGTGGGATCGTGCTGAGGCGGA
CCTGGCGGTTGCGCTGGAAGAAAACAACATCCCGTTTGAATATCAACTGGGTGAAGGCGCTTTCTACGGTCCGAAAATTGAATTTACCCTGTATGAC
TGCCTCGATCGTGCATGGCAGTGCGGTACAGTACAGCTGGACTTCTCTTTGCCGTCTCGTCTGAGCGCTTCTTATGTAGGCGAAGACAATGAACGTA
AAGTACCGGTAATGATTCACCGCGCAATTCTGGGGTCGATGGAACGTTTCATCGGTATCCTGACCGAAGAGTTCGCTGGTTTCTTCCCGACCTGGCT
TGCGCCGGTTCAGGTTGTTATCATGAATATTACCGATTCACAGTCTGAATACGTTAACGAATTGACGCAAAAACTATCAAATGCGGGCATTCGTGTT
AAAGCAGACTTGAGAAATGAGAAGATTGGCTTTAAAATCCGCGAGCACACTTTGCGTCGCGTCCCATATATGCTGGTCTGTGGTGATAAAGAGGTGG
AATCAGGCAAAGTTGCCGTTCGCACCCGCCGTGGTAAAGACCTGGGAAGCATGGACGTAAATGAAGTGATCGAGAAGCTGCAACAAGAGATTCGCAG
CCGCAGTCTTAAACAATTGGAGGAATAATGAgatccggctgctaacaaagcccgaaaggaagctgagttggctgctgccaccgctgagcaataacta
gcataaccccttggggcctctaaacgggtcttgaggggttttttgctgaaaggaggaactatatccggattggcgaatgggacgcgccctgtagcgg
cgcattaagcgcggcgggtgtggtggttacgcgcagcgtgaccgctacacttgccagcgccctagcgcccgctcctttcgctttcttcccttccttt
ctcgccacgttcgccggctttccccgtcaagctctaaatcgggggctccctttagggttccgatttagtgctttacggcacctcgaccccaaaaaac
ttgattagggtgatggttcacgtagtgggccatcgccctgatagacggtttttcgccctttgacgttggagtccacgttctttaatagtggactctt
gttccaaactggaacaacactcaaccctatctcggtctattcttttgatttataagggattttgccgatttcggcctattggttaaaaaatgagctg
atttaacaaaaatttaacgcgaattttaacaaaatattaacgcttacaatttaggtggcacttttcggggaaatgtgcgcggaacccctatttgttt
atttttctaaatacattcaaatatgtatccgctcatgagacaataaccctgataaatgcttcaataatattgaaaaaggaagagtatgagtattcaa
catttccgtgtcgcccttattcccttttttgcggcattttgccttcctgtttttgctcacccagaaacgctggtgaaagtaaaagatgctgaagatc
agttgggtgcacgagtgggttacatcgaactggatctcaacagcggtaagatccttgagagttttcgccccgaagaacgttttccaatgatgagcac
ttttaaagttctgctatgtggcgcggtattatcccgtattgacgccgggcaagagcaactcggtcgccgcatacactattctcagaatgacttggtt
gagtactcaccagtcacagaaaagcatcttacggatggcatgacagtaagagaattatgcagtgctgccataaccatgagtgataacactgcggcca
acttacttctgacaacgatcggaggaccgaaggagctaaccgcttttttgcacaacatgggggatcatgtaactcgccttgatcgttgggaaccgga
gctgaatgaagccataccaaacgacgagcgtgacaccacgatgcctgcagcaatggcaacaacgttgcgcaaactattaactggcgaactacttact
ctagcttcccggcaacaattaatagactggatggaggcggataaagttgcaggaccacttctgcgctcggcccttccggctggctggtttattgctg
ataaatctggagccggtgagcgtgggtctcgcggtatcattgcagcactggggccagatggtaagccctcccgtatcgtagttatctacacgacggg
gagtcaggcaactatggatgaacgaaatagacagatcgctgagataggtgcctcactgattaagcattggtaactgtcagaccaagtttactcatat
atactttagattgatttaaaacttcatttttaatttaaaaggatctaggtgaagatcctttttgataatctcatgaccaaaatcccttaacgtgagt
tttcgttccactgagcgtcagaccccgtagaaaagatcaaaggatcttcttgagatcctttttttctgcgcgtaatctgctgcttgcaaacaaaaaa
accaccgctaccagcggtggtttgtttgccggatcaagagctaccaactctttttccgaaggtaactggcttcagcagagcgcagataccaaatact
gtccttctagtgtagccgtagttaggccaccacttcaagaactctgtagcaccgcctacatacctcgctctgctaatcctgttaccagtggctgctg
ccagtggcgataagtcgtgtcttaccgggttggactcaagacgatagttaccggataaggcgcagcggtcgggctgaacggggggttcgtgcacaca
gcccagcttggagcgaacgacctacaccgaactgagatacctacagcgtgagctatgagaaagcgccacgcttcccgaagggagaaaggcggacagg
tatccggtaagcggcagggtcggaacaggagagcgcacgagggagcttccagggggaaacgcctggtatctttatagtcctgtcgggtttcgccacc
tctgacttgagcgtcgatttttgtgatgctcgtcaggggggcggagcctatggaaaaacgccagcaacgcggcctttttacggttcctggccttttg
ctggccttttgctcacatgttctttcctgcgttatcccctgattctgtggataaccgtattaccgcctttgagtgagctgataccgctcgccgcagc
cgaacgaccgagcgcagcgagtcagtgagcgaggaagcggaagagcgcctgatgcggtattttctccttacgcatctgtgcggtatttcacaccgca
atggtgcactctcagtacaatctgctctgatgccgcatagttaagccagtatacactccgctatcgctacgtgactgggtcatggctgcgccccgac
acccgccaacacccgctgacgcgccctgacgggcttgtctgctcccggcatccgcttacagacaagctgtgaccgtctccgggagctgcatgtgtca
gaggttttcaccgtcatcaccgaaacgcgcgaggcagctgcggtaaagctcatcagcgtggtcgtgaagcgattcacagatgtctgcctgttcatcc
gcgtccagctcgttgagtttctccagaagcgttaatgtctggcttctgataaagcgggccatgttaagggcggttttttcctgtttggtcactgatg
cctccgtgtaagggggatttctgttcatgggggtaatgataccgatgaaacgagagaggatgctcacgatacgggttactgatgatgaacatgcccg
gttactggaacgttgtgagggtaaacaactggcggtatggatgcggcgggaccagagaaaaatcactcagggtcaatgccagcgcttcgttaataca
gatgtaggtgttccacagggtagccagcagcatcctgcgatgcagatccggaacataatggtgcagggcgctgacttccgcgtttccagactttacg
aaacacggaaaccgaagaccattcatgttgttgctcaggtcgcagacgttttgcagcagcagtcgcttcacgttcgctcgcgtatcggtgattcatt
ctgctaaccagtaaggcaaccccgccagcctagccgggtcctcaacgacaggagcacgatcatgcgcacccgtggggccgccatgccggcgataatg

gcctgcttctcgccgaaacgtttggtggcgggaccagtgacgaaggcttgagcgagggcgtgcaagattccgaataccgcaagcgacaggccgatca
tcgtcgcgctccagcgaaagcggtcctcgccgaaaatgacccagagcgctgccggcacctgtcctacgagttgcatgataaagaagacagtcataag
tgcggcgacgatagtcatgccccgcgcccaccggaaggagctgactgggttgaaggctctcaagggcatcggtcgagatcccggtgcctaatgagtg
agctaacttacattaattgcgttgcgctcactgcccgctttccagtcgggaaacctgtcgtgccagctgcattaatgaatcggccaacgcgcgggga
gaggcggtttgcgtattgggcgccagggtggtttttcttttcaccagtgagacgggcaacagctgattgcccttcaccgcctggccctgagagagtt
gcagcaagcggtccacgctggtttgccccagcaggcgaaaatcctgtttgatggtggttaacggcgggatataacatgagctgtcttcggtatcgtc
gtatcccactaccgagatatccgcaccaacgcgcagcccggactcggtaatggcgcgcattgcgcccagcgccatctgatcgttggcaaccagcatc
gcagtgggaacgatgccctcattcagcatttgcatggtttgttgaaaaccggacatggcactccagtcgccttcccgttccgctatcggctgaattt
gattgcgagtgagatatttatgccagccagccagacgcagacgcgccgagacagaacttaatgggcccgctaacagcgcgatttgctggtgacccaa
tgcgaccagatgctccacgcccagtcgcgtaccgtcttcatgggagaaaataatactgttgatgggtgtctggtcagagacatcaagaaataacgcc
ggaacattagtgcaggcagcttccacagcaatggcatcctggtcatccagcggatagttaatgatcagcccactgacgcgttgcgcgagaagattgt
gcaccgccgctttacaggcttcgacgccgcttcgttctaccatcgacaccaccacgctggcacccagttgatcggcgcgagatttaatcgccgcgac
aatttgcgacggcgcgtgcagggccagactggaggtggcaacgccaatcagcaacgactgtttgcccgccagttgttgtgccacgcggttgggaatg
taattcagctccgccatcgccgcttccactttttcccgcgttttcgcagaaacgtggctggcctggttcaccacgcgggaaacggtctgataagaga
caccggcatactctgcgacatcgtataacgttactggtttcacattcaccaccctgaattgactctcttccgggcgctatcatgccataccgcgaaa
ggttttgcgccattcgatggtgtccgggatctcgacgctctcccttatgcgactcctgcattaggaagcagcccagtagtaggttgaggccgttgag
caccgccgccgcaaggaatggtgcatgcaaggagatggcgcccaacagtcccccggccacggggcctgccaccatacccacgccgaaacaagcgctc
atgagcccgaagtggcgagcccgatcttccccatcggtgatgtcggcgatataggcgccagcaaccgcacctgtggcgccggtgatgccggccacga
tgcgtccggcgtagaggatcgagatctcgatcccgcgaaat

### *Sequence of pET21a-His-IF1*

#### $P_{T7}$ promoter - His6-IF1 - T7 terminator

taatacgactcactataggggaattgtgagcggataacaattccccctctagaaataattttgtttaactttaagaaggagatatacatATGAGAGGA
TCGCATCACCATCACCATCACGGATCCATGGCCAAAGAAGACAATATTGAAATGCAAGGTACCGTTCTTGAAACGTTGCCTAATACCATGTTCCGCG
TAGAGTTAGAAAACGGTCACGTGGTTACTGCACACATCTCCGGTAAAATGCGCAAAAACTACATCCGCATCCTGACGGGCGACAAAGTGACTGTTGA
ACTGACCCCGTACGACCTGAGCAAAGGCCGCATTGTCTTCCGTAGTCGCTGAgatccggctgctaacaaagcccgaaaggaagctgagttggctgct
gccaccgctgagcaataactagcataaccccttggggcctctaaacgggtcttgaggggttttttgctgaaaggaggaactatatccggattggcga
atgggacgcgccctgtagcggcgcattaagcgcggcgggtgtggtggttacgcgcagcgtgaccgctacacttgccagcgccctagcgcccgctcct
ttcgctttcttcccttcctttctcgccacgttcgccggctttccccgtcaagctctaaatcgggggctccctttagggttccgatttagtgctttac
ggcacctcgaccccaaaaaacttgattagggtgatggttcacgtagtgggccatcgccctgatagacggtttttcgccctttgacgttggagtccac
gttctttaatagtggactcttgttccaaactggaacaacactcaaccctatctcggtctattcttttgatttataagggattttgccgatttcggcc
tattggttaaaaaatgagctgatttaacaaaaatttaacgcgaattttaacaaaatattaacgcttacaatttaggtggcacttttcggggaaatgt
gcgcggaacccctatttgtttatttttctaaatacattcaaatatgtatccgctcatgagacaataaccctgataaatgcttcaataatattgaaaa
aggaagagtatgagtattcaacatttccgtgtcgcccttattcccttttttgcggcattttgccttcctgtttttgctcacccagaaacgctggtga
aagtaaaagatgctgaagatcagttgggtgcacgagtgggttacatcgaactggatctcaacagcggtaagatccttgagagttttcgccccgaaga
acgttttccaatgatgagcacttttaaagttctgctatgtggcgcggtattatcccgtattgacgccgggcaagagcaactcggtcgccgcatacac
tattctcagaatgacttggttgagtactcaccagtcacagaaaagcatcttacggatggcatgacagtaagagaattatgcagtgctgccataacca
tgagtgataacactgcggccaacttacttctgacaacgatcggaggaccgaaggagctaaccgcttttttgcacaacatgggggatcatgtaactcg
ccttgatcgttgggaaccggagctgaatgaagccataccaaacgacgagcgtgacaccacgatgcctgcagcaatggcaacaacgttgcgcaaacta
ttaactggcgaactacttactctagcttcccggcaacaattaatagactggatggaggcggataaagttgcaggaccacttctgcgctcggcccttc
cggctggctggtttattgctgataaatctggagccggtgagcgtgggtctcgcggtatcattgcagcactggggccagatggtaagccctcccgtat
cgtagttatctacacgacggggagtcaggcaactatggatgaacgaaatagacagatcgctgagataggtgcctcactgattaagcattggtaactg
tcagaccaagtttactcatatatactttagattgatttaaaacttcatttttaatttaaaaggatctaggtgaagatcctttttgataatctcatga
ccaaaatcccttaacgtgagttttcgttccactgagcgtcagaccccgtagaaaagatcaaaggatcttcttgagatcctttttttctgcgcgtaat
ctgctgcttgcaaacaaaaaaaccaccgctaccagcggtggtttgtttgccggatcaagagctaccaactctttttccgaaggtaactggcttcagc
agagcgcagataccaaatactgtccttctagtgtagccgtagttaggccaccacttcaagaactctgtagcaccgcctacatacctcgctctgctaa
tcctgttaccagtggctgctgccagtggcgataagtcgtgtcttaccgggttggactcaagacgatagttaccggataaggcgcagcggtcgggctg
aacggggggttcgtgcacacagcccagcttggagcgaacgacctacaccgaactgagatacctacagcgtgagctatgagaaagcgccacgcttccc
gaagggagaaaggcggacaggtatccggtaagcggcagggtcggaacaggagagcgcacgagggagcttccagggggaaacgcctggtatctttata
gtcctgtcgggtttcgccacctctgacttgagcgtcgatttttgtgatgctcgtcaggggggcggagcctatggaaaaacgccagcaacgcggcctt
tttacggttcctggccttttgctggccttttgctcacatgttctttcctgcgttatcccctgattctgtggataaccgtattaccgcctttgagtga
gctgataccgctcgccgcagccgaacgaccgagcgcagcgagtcagtgagcgaggaagcggaagagcgcctgatgcggtattttctccttacgcatc
tgtgcggtatttcacaccgcaatggtgcactctcagtacaatctgctctgatgccgcatagttaagccagtatacactccgctatcgctacgtgact
gggtcatggctgcgccccgacacccgccaacacccgctgacgcgccctgacgggcttgtctgctcccggcatccgcttacagacaagctgtgaccgt
ctccgggagctgcatgtgtcagaggttttcaccgtcatcaccgaaacgcgcgaggcagctgcggtaaagctcatcagcgtggtcgtgaagcgattca
cagatgtctgcctgttcatccgcgtccagctcgttgagtttctccagaagcgttaatgtctggcttctgataaagcgggccatgttaagggcggttt
tttcctgtttggtcactgatgcctccgtgtaagggggatttctgttcatgggggtaatgataccgatgaaacgagagaggatgctcacgatacgggt
tactgatgatgaacatgcccggttactggaacgttgtgagggtaaacaactggcggtatggatgcggcgggaccagagaaaaatcactcagggtcaa
tgccagcgcttcgttaatacagatgtaggtgttccacagggtagccagcagcatcctgcgatgcagatccggaacataatggtgcagggcgctgact
tccgcgtttccagactttacgaaacacggaaaccgaagaccattcatgttgttgctcaggtcgcagacgttttgcagcagcagtcgcttcacgttcg
ctcgcgtatcggtgattcattctgctaaccagtaaggcaaccccgccagcctagccgggtcctcaacgacaggagcacgatcatgcgcacccgtggg
gccgccatgccggcgataatggcctgcttctcgccgaaacgtttggtggcgggaccagtgacgaaggcttgagcgagggcgtgcaagattccgaata
ccgcaagcgacaggccgatcatcgtcgcgctccagcgaaagcggtcctcgccgaaaatgacccagagcgctgccggcacctgtcctacgagttgcat
gataaagaagacagtcataagtgcggcgacgatagtcatgccccgcgcccaccggaaggagctgactgggttgaaggctctcaagggcatcggtcga
gatcccggtgcctaatgagtgagctaacttacattaattgcgttgcgctcactgcccgctttccagtcgggaaacctgtcgtgccagctgcattaat
gaatcggccaacgcgcggggagaggcggtttgcgtattgggcgccagggtggtttttcttttcaccagtgagacgggcaacagctgattgcccttca
ccgcctggccctgagagagttgcagcaagcggtccacgctggtttgccccagcaggcgaaaatcctgtttgatggtggttaacggcgggatataaca
tgagctgtcttcggtatcgtcgtatcccactaccgagatatccgcaccaacgcgcagcccggactcggtaatggcgcgcattgcgcccagcgccatc
tgatcgttggcaaccagcatcgcagtgggaacgatgccctcattcagcatttgcatggtttgttgaaaaccggacatggcactccagtcgccttccc
gttccgctatcggctgaatttgattgcgagtgagatatttatgccagccagccagacgcagacgcgccgagacagaacttaatgggcccgctaacag
cgcgatttgctggtgacccaatgcgaccagatgctccacgcccagtcgcgtaccgtcttcatgggagaaaataatactgttgatgggtgtctggtca
gagacatcaagaaataacgccggaacattagtgcaggcagcttccacagcaatggcatcctggtcatccagcggatagttaatgatcagcccactga
cgcgttgcgcgagaagattgtgcaccgccgctttacaggcttcgacgccgcttcgttctaccatcgacaccaccacgctggcacccagttgatcggc
gcgagatttaatcgccgcgacaatttgcgacggcgcgtgcagggccagactggaggtggcaacgccaatcagcaacgactgtttgcccgccagttgt
tgtgccacgcggttgggaatgtaattcagctccgccatcgccgcttccactttttcccgcgttttcgcagaaacgtggctggcctggttcaccacgc
gggaaacggtctgataagagacaccggcatactctgcgacatcgtataacgttactggtttcacattcaccaccctgaattgactctcttccgggcg
ctatcatgccataccgcgaaaggttttgcgccattcgatggtgtccgggatctcgacgctctcccttatgcgactcctgcattaggaagcagcccag
tagtaggttgaggccgttgagcaccgccgccgcaaggaatggtgcatgcaaggagatggcgcccaacagtcccccggccacggggcctgccaccata

cccacgccgaaacaagcgctcatgagcccgaagtggcgagcccgatcttccccatcggtgatgtcggcgatataggcgccagcaaccgcacctgtgg
cgccggtgatgccggccacgatgcgtccggcgtagaggatcgagatctcgatcccgcgaaat

### *Sequence of pET21a-His-IF2*

$P_{T7}$ promoter - His6-IF2 - T7 terminator

taatacgactcactataggggaattgtgagcggataacaattccсctctagaaataattttgtttaactttaagaaggagatatacatATGAGAGGA
TCGCATCACCATCACCATCACGGATCCATGACAGATGTAACGATTAAAACGCTGGCCGCAGAGCGACAGACCTCCGTGGAACGCCTGGTACAGCAAT
TTGCTGATGCAGGTATCCGGAAGTCTGCTGACGACTCTGTGTCTGCACAAGAGAAACAGACTTTGATTGACCACCTGAATCAGAAAAATTCAGGCCC
GGACAAATTGACGCTGCAACGTAAAACACGCAGCACCCTTAACATTCCTGGTACCGGTGGAAAAAGCAAATCGGTACAAATCGAAGTCCGCAAGAAA
CGCACCTTTGTGAAACGCGATCCGCAAGAGGCTGAACGCCTTGCAGCGGAAGAGCAAGCGCAGCGTGAAGCGGAAGAGCAAGCCCGTCGTGAGGCAG
AAGAATCGGCTAAACGCGAGGCGCAACAAAAAGCTGAACGTGAGGCCGCAGAACAAGCTAAGCGTGAAGCTGCTGAACAAGCGAAACGTGAAGCTGC
GGAAAAAGACAAAGTGAGCAATCAACAAGACGATATGACTAAAAACGCCCAGGCTGAAAAAGCCCGCCGTGAGCAGGAAGCTGCAGAGCTCAAGCGT
AAAGCTGAAGAAGAAGCGCGTCGTAAACTCGAAGAAGAAGCACGTCGCGTTGCTGAAGAAGCACGTCGTATGGCGGAAGAAAACAAATGGACTGATA
ACGCGGAACCGACTGAAGATTCCAGCGATTATCACGTCACTACTTCTCAACATGCTCGCCAGGCAGAAGACGAAAGCGATCGTGAAGTCGAAGGCGG
CCGTGGCCGTGGTCGTAACGCGAAAGCAGCGCGTCCGAAGAAAGGCAACAAACACGCTGAATCAAAAGCTGATCGTGAAGAAGCACGCGCAGCAGTA
CGTGGCGGTAAAGGCGGAAAACGTAAAGGTTCTTCGCTGCAGCAAGGCTTCCAGAAGCCTGCTCAGGCCGTTAACCGTGACGTTGTGATCGGCGAAA
CTATCACCGTTGGCGAACTGGCGAACAAGATGGCGGTTAAAGGCTCTCAGGTCATCAAAGCGATGATGAAACTGGGCGCAATGGCAACCATCAACCA
GGTTATCGATCAGGAAACCGCACAGCTGGTTGCTGAAGAGATGGGCCATAAAGTTATCCTGCGTCGTGAAAACGAGCTGGAAGAGGCGGTAATGAGC
GACCGTGACACGGGTGCTGCGGCTGAACCGCGCGCGCCGGTTGTGACCATCATGGGTCACGTTGACCACGGTAAAACCTCTCTGCTGGACTACATTC
GTTCAACGAAAGTGGCCTCTGGCGAAGCGGGCGGCATTACCCAGCACATTGGTGCATACCACGTTGAAACTGAAAACGGCATGATCACCTTCCTGGA
CACCCCGGGGCACGCCGCGTTTACTTCAATGCGTGCTCGTGGTGCGCAGGCAACGGACATCGTAGTCCTGGTTGTTGCTGCCGACGACGGTGTGATG
CCGCAGACCATCGAAGCAATCCAGCACGCGAAAGCGGCGCAGGTACCGGTGGTGGTTGCAGTGAACAAGATCGATAAACCAGAAGCTGATCCGGATC
GCGTTAAGAACGAACTCTCCCAGTACGGCATCCTGCCGGAAGAGTGGGGCGGTGAAAGCCAGTTCGTACACGTATCTGCGAAAGCGGGTACCGGTAT
CGATGAACTGCTGGACGCTATCCTGCTGCAGGCGGAAGTTCTGGAGCTGAAAGCGGTACGTAAAGGTATGGCGAGCGGTGCGGTTATCGAATCCTTC
CTCGATAAAGGTCGTGGTCCGGTTGCTACCGTTCTGGTACGTGAAGGTACTCTGCACAAGGGCGATATCGTTCTGTGTGGCTTCGAATACGGTCGTG
TTCGTGCGATGCGTAACGAACTGGGTCAGGAAGTGCTGGAAGCGGGTCCGTCCATTCCGGTGGAAATCCTCGGCCTGTCCGGCGTACCGGCTGCGGG
TGATGAAGTTACCGTTGTACGTGACGAGAAGAAAGCGCGTGAAGTTGCACTCTATCGTCAGGGTAAATTCCGCGAAGTTAAACTGGCGCGTCAGCAG
AAATCTAAACTCGAGAACATGTTCGCCAACATGACCGAAGGCGAAGTTCACGAAGTGAATATCGTCCTGAAGGCAGACGTACAGGGTTCTGTCGAAG
CGATCTCCGACTCCTTGCTGAAACTGTCTACTGACGAAGTTAAAGTGAAGATCATCGGTTCTGGCGTAGGTGGTATCACCGAAACCGACGCCACCCT
GGCTGCGGCGTCCAACGCCATCCTGGTTGGCTTTAACGTACGTGCTGATGCCTCTGCACGTAAAGTGATTGAAGCGGAAAGCCTGGATCTGCGTTAC
TACTCCGTCATCTATAACCTGATTGACGAAGTGAAAGCGGCGATGAGCGGTATGCTGTCTCCGGAACTGAAACAGCAGATTATCGGTCTGGCGGAAG
TTCGTGACGTGTTCAAATCGCCGAAATTTGGTGCCATCGCAGGCTGTATGGTTACCGAAGGTGTGGTTAAACGTCACAACCCGATCCGCGTTCTGCG
TGACAACGTGGTTATCTACGAAGGCGAGCTGGAGTCCCTGCGCCGCTTCAAAGATGACGTTAACGAAGTCCGTAACGGTATGGAATGTGGTATCGGC
GTTAAGAACTACAACGACGTCCGCACTGGCGATGTGATCGAAGTATTCGAAATCATCGAGATCCAACGTACCATTGCTTAATGAgatccggctgcta
acaaagcccgaaaggaagctgagttggctgctgccaccgctgagcaataactagcataaccccttggggcctctaaacgggtcttgaggggttttt
gctgaaaggaggaactatatccggattggcgaatgggacgcgccctgtagcggcgcattaagcgcggcgggtgtggtggttacgcgcagcgtgaccg
ctacacttgccagcgccctagcgcccgctcctttcgctttcttcccttcctttctcgccacgttcgccggctttccccgtcaagctctaaatcgggg
gctccctttagggttccgatttagtgctttacggcacctcgaccccaaaaaacttgattagggtgatggttcacgtagtgggccatcgccctgatag
acggtttttcgccctttgacgttggagtccacgttctttaatagtggactcttgttccaaactggaacaacactcaaccctatctcggtctattctt
ttgatttataagggattttgccgatttcggcctattggttaaaaaatgagctgatttaacaaaaatttaacgcgaattttaacaaaatattaacgct
tacaatttaggtggcacttttcggggaaatgtgcgcggaacccctatttgtttatttttctaaatacattcaaatatgtatccgctcatgagacaat
aaccctgataaatgcttcaataatattgaaaaaggaagagtatgagtattcaacatttccgtgtcgcccttattcccttttttgcggcattttgcct
tcctgtttttgctcacccagaaacgctggtgaaagtaaaagatgctgaagatcagttgggtgcacgagtgggttacatcgaactggatctcaacagc
ggtaagatccttgagagttttcgccccgaagaacgttttccaatgatgagcacttttaaagttctgctatgtggcgcggtattatcccgtattgacg
ccgggcaagagcaactcggtcgccgcatacactattctcagaatgacttggttgagtactcaccagtcacagaaaagcatcttacggatggcatgac
agtaagagaattatgcagtgctgccataaccatgagtgataacactgcggccaacttacttctgacaacgatcggaggaccgaaggagctaaccgct
tttttgcacaacatgggggatcatgtaactcgccttgatcgttgggaaccggagctgaatgaagccataccaaacgacgagcgtgacaccacgatgc
ctgcagcaatggcaacaacgttgcgcaaactattaactggcgaactacttactctagcttcccggcaacaattaatagactggatggaggcggataa
agttgcaggaccacttctgcgctcggcccttccggctggctggtttattgctgataaatctggagccggtgagcgtgggtctcgcggtatcattgca
gcactggggccagatggtaagccctcccgtatcgtagttatctacacgacggggagtcaggcaactatggatgaacgaaatagacagatcgctgaga
taggtgcctcactgattaagcattggtaactgtcagaccaagtttactcatatatactttagattgatttaaaacttcatttttaatttaaaaggat
ctaggtgaagatcctttttgataatctcatgaccaaaatcccttaacgtgagttttcgttccactgagcgtcagaccccgtagaaaagatcaaagga
tcttcttgagatcctttttttctgcgcgtaatctgctgcttgcaaacaaaaaaaccaccgctaccagcggtggtttgtttgccggatcaagagctac
caactctttttccgaaggtaactggcttcagcagagcgcagataccaaatactgtccttctagtgtagccgtagttaggccaccacttcaagaactc
tgtagcaccgcctacatacctcgctctgctaatcctgttaccagtggctgctgccagtggcgataagtcgtgtcttaccgggttggactcaagacga
tagttaccggataaggcgcagcggtcgggctgaacggggggttcgtgcacacagcccagcttggagcgaacgacctacaccgaactgagatacctac
agcgtgagctatgagaaagcgccacgcttcccgaagggagaaaggcggacaggtatccggtaagcggcagggtcggaacaggagagcgcacgaggga
gcttccagggggaaacgcctggtatctttatagtcctgtcgggtttcgccacctctgacttgagcgtcgatttttgtgatgctcgtcaggggggcgg
agcctatggaaaaacgccagcaacgcggcctttttacggttcctggccttttgctggccttttgctcacatgttctttcctgcgttatcccctgatt
ctgtggataaccgtattaccgcctttgagtgagctgataccgctcgccgcagccgaacgaccgagcgcagcgagtcagtgagcgaggaagcggaaga
gcgcctgatgcggtattttctccttacgcatctgtgcggtatttcacaccgcaatggtgcactctcagtacaatctgctctgatgccgcatagttaa
gccagtatacactccgctatcgctacgtgactgggtcatggctgcgccccgacacccgccaacacccgctgacgcgccctgacgggcttgtctgctc

ccggcatccgcttacagacaagctgtgaccgtctccgggagctgcatgtgtcagaggttttcaccgtcatcaccgaaacgcgcgaggcagctgcggt
aaagctcatcagcgtggtcgtgaagcgattcacagatgtctgcctgttcatccgcgtccagctcgttgagtttctccagaagcgttaatgtctggct
tctgataaagcgggccatgttaagggcggtttttttcctgtttggtcactgatgcctccgtgtaagggggatttctgttcatgggggtaatgataccg
atgaaacgagagaggatgctcacgatacgggttactgatgatgaacatgcccggttactggaacgttgtgagggtaaacaactggcggtatggatgc
ggcgggaccagagaaaaatcactcagggtcaatgccagcgcttcgttaatacagatgtaggtgttccacagggtagccagcagcatcctgcgatgca
gatccggaacataatggtgcagggcgctgacttccgcgtttccagactttacgaaacacggaaaccgaagaccattcatgttgttgctcaggtcgca
gacgttttgcagcagcagtcgcttcacgttcgctcgcgtatcggtgattcattctgctaaccagtaaggcaaccccgccagcctagccgggtcctca
acgacaggagcacgatcatgcgcacccgtggggccgccatgccggcgataatggcctgcttctcgccgaaacgtttggtggcgggaccagtgacgaa
ggcttgagcgagggcgtgcaagattccgaataccgcaagcgacaggccgatcatcgtcgcgctccagcgaaagcggtcctcgccgaaaatgacccag
agcgctgccggcacctgtcctacgagttgcatgataaagaagacagtcataagtgcggcgacgatagtcatgccccgcgcccaccggaaggagctga
ctgggttgaaggctctcaagggcatcggtcgagatcccggtgcctaatgagtgagctaacttacattaattgcgttgcgctcactgcccgctttcca
gtcgggaaacctgtcgtgccagctgcattaatgaatcggccaacgcgcggggagaggcggtttgcgtattgggcgccagggtggtttttcttttcac
cagtgagacgggcaacagctgattgcccttcaccgcctggccctgagagagttgcagcaagcggtccacgctggtttgccccagcaggcgaaaatcc
tgtttgatggtggttaacggcgggatataacatgagctgtcttcggtatcgtcgtatcccactaccgagatatccgcaccaacgcgcagcccggact
cggtaatggcgcgcattgcgcccagcgccatctgatcgttggcaaccagcatcgcagtgggaacgatgccctcattcagcatttgcatggtttgttg
aaaaccggacatggcactccagtcgccttcccgttccgctatcggctgaatttgattgcgagtgagatatttatgccagccagccagacgcagacgc
gccgagacagaacttaatgggcccgctaacagcgcgatttgctggtgacccaatgcgaccagatgctccacgcccagtcgcgtaccgtcttcatggg
agaaaataatactgttgatgggtgtctggtcagagacatcaagaaataacgccggaacattagtgcaggcagcttccacagcaatggcatcctggtc
atccagcggatagttaatgatcagcccactgacgcgttgcgcgagaagattgtgcaccgccgctttacaggcttcgacgccgcttcgttctaccatc
gacaccaccacgctggcacccagttgatcggcgcgagatttaatcgccgcgacaatttgcgacggcgcgtgcagggccagactggaggtggcaacgc
caatcagcaacgactgtttgcccgccagttgttgtgccacgcggttgggaatgtaattcagctccgccatcgccgcttccactttttcccgcgtttt
cgcagaaacgtggctggcctggttcaccacgcgggaaacggtctgataagagacaccggcatactctgcgacatcgtataacgttactggtttcaca
ttcaccaccctgaattgactctcttccgggcgctatcatgccataccgcgaaaggttttgcgccattcgatggtgtccgggatctcgacgctctccc
ttatgcgactcctgcattaggaagcagcccagtagtaggttgaggccgttgagcaccgccgccgcaaggaatggtgcatgcaaggagatggcgccca
acagtcccccggccacggggcctgccaccatacccacgccgaaacaagcgctcatgagcccgaagtggcgagcccgatcttccccatcggtgatgtc
ggcgatataggcgccagcaaccgcacctgtggcgccggtgatgccggccacgatgcgtccggcgtagaggatcgagatctcgatcccgcgaaat

SUPPLEMENTARY INFORMATION I. Plasmid sequences used in this work

#### ***Sequence of pET21a-His-IF3***

$P_{T7}$ promoter - His6-IF3 - T7 terminator

```
taatacgactcactataggggaattgtgagcggataacaattcccctctagaaataattttgtttaactttaagaaggagatatacatATGAGAGGA
TCGCATCACCATCACCATCACGGATCCATTAAAGGCGGAAAACGAGTTCAAACGGCGCGCCCTAACCGTATCAATGGCGAAATTCGCGCCCAGGAAG
TTCGCTTAACAGGTCTGGAAGGCGAGCAGCTTGGTATTGTGAGTCTGAGAGAAGCTCTGGAGAAAGCAGAAGAAGCCGGAGTAGACTTAGTCGAGAT
CAGCCCTAACGCCGAGCCGCCGGTTTGTCGTATAATGGATTACGGCAAATTCCTCTATGAAAAGAGCAAGTCTTCTAAGGAACAGAAGAAAAAGCAA
AAAGTTATCCAGGTTAAGGAAATTAAATTCCGTCCTGGTACAGATGAAGGCGACTATCAGGTAAAACTCCGCAGCCTGATTCGCTTTCTCGAAGAGG
GTGATAAAGCCAAAATCACGCTGCGTTTCCGCGGTCGTGAGATGGCGCACCAGCAAATCGGTATGGAAGTGCTTAATCGCGTGAAAGACGATTTGCA
AGAACTGGCAGTGGTCGAATCCTTCCCAACGAAGATCGAAGGCCGCCAGATGATCATGGTGCTCGCTCCTAAGAAGAAACAGTAATGAgatccggct
gctaacaaagcccgaaaggaagctgagttggctgctgccaccgctgagcaataactagcataaccccttggggcctctaaacgggtcttgaggggtt
ttttgctgaaaggaggaactatatccggattggcgaatgggacgcgccctgtagcggcgcattaagcgcggcgggtgtggtggttacgcgcagcgtg
accgctacacttgccagcgccctagcgcccgctcctttcgctttcttcccttcctttctcgccacgttcgccggctttccccgtcaagctctaaatc
gggggctccctttagggttccgatttagtgctttacggcacctcgaccccaaaaaacttgattagggtgatggttcacgtagtgggccatcgccctg
atagacggtttttcgccctttgacgttggagtccacgttctttaatagtggactcttgttccaaactggaacaacactcaaccctatctcggtctat
tcttttgatttataagggattttgccgatttcggcctattggttaaaaaatgagctgatttaacaaaaatttaacgcgaattttaacaaaatattaa
cgcttacaatttaggtggcacttttcggggaaatgtgcgcggaacccctatttgtttatttttctaaatacattcaaatatgtatccgctcatgaga
caataaccctgataaatgcttcaataatattgaaaaaggaagagtatgagtattcaacatttccgtgtcgcccttattcccttttttgcggcatttt
gccttcctgtttttgctcacccagaaacgctggtgaaagtaaaagatgctgaagatcagttgggtgcacgagtgggttacatcgaactggatctcaa
cagcggtaagatccttgagagttttcgccccgaagaacgttttccaatgatgagcacttttaaagttctgctatgtggcgcggtattatcccgtatt
gacgccgggcaagagcaactcggtcgccgcatacactattctcagaatgacttggttgagtactcaccagtcacagaaaagcatcttacggatggca
tgacagtaagagaattatgcagtgctgccataaccatgagtgataacactgcggccaacttacttctgacaacgatcggaggaccgaaggagctaac
cgcttttttgcacaacatgggggatcatgtaactcgccttgatcgttgggaaccggagctgaatgaagccataccaaacgacgagcgtgacaccacg
atgcctgcagcaatggcaacaacgttgcgcaaactattaactggcgaactacttactctagcttcccggcaacaattaatagactggatggaggcgg
ataaagttgcaggaccacttctgcgctcggcccttccggctggctggtttattgctgataaatctggagccggtgagcgtgggtctcgcggtatcat
tgcagcactggggccagatggtaagccctcccgtatcgtagttatctacacgacggggagtcaggcaactatggatgaacgaaatagacagatcgct
gagataggtgcctcactgattaagcattggtaactgtcagaccaagtttactcatatatactttagattgatttaaaacttcatttttaatttaaaa
ggatctaggtgaagatcctttttgataatctcatgaccaaaatcccttaacgtgagttttcgttccactgagcgtcagaccccgtagaaaagatcaa
aggatcttcttgagatcctttttttctgcgcgtaatctgctgcttgcaaacaaaaaaaccaccgctaccagcggtggtttgtttgccggatcaagag
ctaccaactctttttccgaaggtaactggcttcagcagagcgcagataccaaatactgtccttctagtgtagccgtagttaggccaccacttcaaga
actctgtagcaccgcctacatacctcgctctgctaatcctgttaccagtggctgctgccagtggcgataagtcgtgtcttaccgggttggactcaag
acgatagttaccggataaggcgcagcggtcgggctgaacggggggttcgtgcacacagcccagcttggagcgaacgacctacaccgaactgagatac
ctacagcgtgagctatgagaaagcgccacgcttcccgaagggagaaaggcggacaggtatccggtaagcggcagggtcggaacaggagagcgcacga
gggagcttccagggggaaacgcctggtatctttatagtcctgtcgggtttcgccacctctgacttgagcgtcgatttttgtgatgctcgtcaggggg
gcggagcctatggaaaaacgccagcaacgcggcctttttacggttcctggccttttgctggccttttgctcacatgttctttcctgcgttatcccct
gattctgtggataaccgtattaccgcctttgagtgagctgataccgctcgccgcagccgaacgaccgagcgcagcgagtcagtgagcgaggaagcgg
aagagcgcctgatgcggtattttctccttacgcatctgtgcggtatttcacaccgcaatggtgcactctcagtacaatctgctctgatgccgcatag
ttaagccagtatacactccgctatcgctacgtgactgggtcatggctgcgccccgacacccgccaacacccgctgacgcgccctgacgggcttgtct
gctcccggcatccgcttacagacaagctgtgaccgtctccgggagctgcatgtgtcagaggttttcaccgtcatcaccgaaacgcgcgaggcagctg
cggtaaagctcatcagcgtggtcgtgaagcgattcacagatgtctgcctgttcatccgcgtccagctcgttgagtttctccagaagcgttaatgtct
ggcttctgataaagcgggccatgttaagggcggttttttcctgtttggtcactgatgcctccgtgtaagggggatttctgttcatgggggtaatgat
accgatgaaacgagagaggatgctcacgatacgggttactgatgatgaacatgcccggttactggaacgttgtgagggtaaacaactggcggtatgg
atgcggcgggaccagagaaaaatcactcagggtcaatgccagcgcttcgttaatacagatgtaggtgttccacagggtagccagcagcatcctgcga
tgcagatccggaacataatggtgcagggcgctgacttccgcgtttccagactttacgaaacacggaaaccgaagaccattcatgttgttgctcaggt
cgcagacgttttgcagcagcagtcgcttcacgttcgctcgcgtatcggtgattcattctgctaaccagtaaggcaaccccgccagcctagccgggtc
ctcaacgacaggagcacgatcatgcgcacccgtggggccgccatgccggcgataatggcctgcttctcgccgaaacgtttggtggcgggaccagtga
cgaaggcttgagcgagggcgtgcaagattccgaataccgcaagcgacaggccgatcatcgtcgcgctccagcgaaagcggtcctcgccgaaaatgac
ccagagcgctgccggcacctgtcctacgagttgcatgataaagaagacagtcataagtgcggcgacgatagtcatgccccgcgcccaccggaaggag
ctgactgggttgaaggctctcaagggcatcggtcgagatcccggtgcctaatgagtgagctaacttacattaattgcgttgcgctcactgcccgctt
tccagtcgggaaacctgtcgtgccagctgcattaatgaatcggccaacgcgcggggagaggcggtttgcgtattgggcgccagggtggtttttcttt
tcaccagtgagacgggcaacagctgattgcccttcaccgcctggccctgagagagttgcagcaagcggtccacgctggtttgccccagcaggcgaaa
atcctgtttgatggtggttaacggcgggatataacatgagctgtcttcggtatcgtcgtatcccactaccgagatatccgcaccaacgcgcagcccg
gactcggtaatggcgcgcattgcgcccagcgccatctgatcgttggcaaccagcatcgcagtgggaacgatgccctcattcagcatttgcatggttt
gttgaaaaccggacatggcactccagtcgccttcccgttccgctatcggctgaatttgattgcgagtgagatatttatgccagccagccagacgcag
acgcgccgagacagaacttaatgggcccgctaacagcgcgatttgctggtgacccaatgcgaccagatgctccacgcccagtcgcgtaccgtcttca
tgggagaaaataatactgttgatgggtgtctggtcagagacatcaagaaataacgccggaacattagtgcaggcagcttccacagcaatggcatcct
ggtcatccagcggatagttaatgatcagcccactgacgcgttgcgcgagaagattgtgcaccgccgctttacaggcttcgacgccgcttcgttctac
catcgacaccaccacgctggcacccagttgatcggcgcgagatttaatcgccgcgacaatttgcgacggcgcgtgcagggccagactggaggtggca
acgccaatcagcaacgactgtttgcccgccagttgttgtgccacgcggttgggaatgtaattcagctccgccatcgccgcttccacttttcccgcg
ttttcgcagaaacgtggctggcctggttcaccacgcgggaaacggtctgataagagacaccggcatactctgcgacatcgtataacgttactggttt
```

cacattcaccaccctgaattgactctcttccgggcgctatcatgccataccgcgaaaggttttgcgccattcgatggtgtccgggatctcgacgctc
tcccttatgcgactcctgcattaggaagcagcccagtagtaggttgaggccgttgagcaccgccgccgcaaggaatggtgcatgcaaggagatggcg
cccaacagtcccccggccacggggcctgccaccatacccacgccgaaacaagcgctcatgagcccgaagtggcgagcccgatcttccccatcggtga
tgtcggcgatataggcgccagcaaccgcacctgtggcgccggtgatgccggccacgatgcgtccggcgtagaggatcgagatctcgatcccgcgaaa
t

### *Sequence of pET21a-EFG-His*

#### $P_{T7}$ promoter – EFG-His6 - T7 terminator

taatacgactcactataggggaattgtgagcggataacaattccctctagaaataattttgtttaactttaagaaggagatatacatATGGCTCGT
ACAACACCCATCGCACGCTACCGTAACATCGGTATCAGTGCGCACATCGACGCCGGTAAAACCACTACTACCGAACGTATTCTGTTCTACACCGGTG
TAAACCATAAAATCGGTGAAGTTCATGACGGCGCTGCAACCATGGACTGGATGGAGCAGGAGCAGGAACGTGGTATTACCATCACTTCCGCTGCGAC
TACTGCATTCTGGTCTGGTATGGCTAAGCAGTATGAGCCGCATCGCATCAACATCATCGACACCCCGGGGCACGTTGACTTCACAATCGAAGTAGAA
CGTTCCATGCGTGTTCTCGATGGTGCGGTAATGGTTTACTGCGCAGTTGGTGGTGTTCAGCCGCAGTCTGAAACCGTATGGCGTCAGGCAAACAAAT
ATAAAGTTCCGCGCATTGCGTTCGTTAACAAAATGGACCGCATGGGTGCGAACTTCCTGAAAGTTGTTAACCAGATCAAAACCCGTCTGGGCGCGAA
CCCGGTTCCGCTGCAGCTGGCGATTGGTGCTGAAGAACATTTCACCGGTGTTGTTGACCTGGTGAAAATGAAAGCTATCAACTGGAACGACGCTGAC
CAGGGCGTAACCTTCGAATACGAAGATATCCCGGCAGACATGGTTGAACTGGCTAACGAATGGCACCAGAACCTGATCGAATCCGCAGCTGAAGCTT
CTGAAGAGCTGATGGAAAAATACCTGGGTGGTGAAGAACTGACTGAAGCAGAAATCAAAGGTGCTCTGCGACAGCGCGTTCTGAACAACGAAATCAT
CCTGGTAACCTGTGGTTCTGCGTTCAAGAACAAAGGTGTTCAGGCGATGCTGGATGCGGTAATTGATTACCTGCCATCCCCGGTTGACGTACCTGCG
ATCAACGGTATCCTGGACGACGGTAAAGACACTCCGGCTGAACGTCACGCAAGTGATGACGAGCCGTTCTCTGCACTGGCGTTCAAAATCGCTACCG
ACCCGTTTGTTGGTAACCTGACCTTCTTCCGTGTTTACTCCGGTGTGGTTAACTCTGGTGATACCGTACTGAACTCCGTGAAAGCTGCACGTGAGCG
TTTCGGTCGTATCGTTCAGATGCACGCTAACAAACGTGAAGAGATCAAAGAAGTTCGCGCGGGCGACATCGCTGCTGCTATCGGTCTGAAAGACGTA
ACCACTGGTGACACCCTGTGTGACCCGGATGCGCCGATCATTCTGGAACGTATGGAATTCCCTGAGCCGGTAATCTCCATCGCAGTTGAACCGAAAA
CCAAAGCTGACCAGGAAAAAATGGGTCTGGCTCTGGGCCGTCTGGCTAAAGAAGACCCGTCTTTCCGTGTATGGACTGACGAAGAATCTAACCAGAC
CATCATCGCGGGTATGGGCGAACTGCACCTCGACATCATCGTTGACCGTATGAAGCGTGAATTCAACGTTGAAGCGAACGTAGGTAAACCGCAGGTT
GCTTACCGTGAAACTATCCGCCAGAAAGTTACCGATGTTGAAGGTAAACACGCGAAACAGTCTGGTGGTCGTGGTCAGTATGGTCATGTTGTTATCG
ACATGTACCCGCTGGAGCCGGGTTCAAACCCGAAAGGCTACGAGTTCATCAACGACATTAAAGGTGGTGTAATCCCTGGCGAATACATCCCGGCCGT
TGATAAAGGTATCCAGGAACAGCTGAAAGCAGGTCCGCTGGCAGGCTACCCGGTAGTAGACATGGGTATTCGTCTGCACTTCGGTTCTTACCATGAC
GTTGACTCCTCTGAACTGGCGTTTAAACTGGCTGCTTCTATCGCCTTTAAAGAAGGCTTTAAGAAAGCGAAACCAGTTCTGCTTGAGCCGATCATGA
AGGTTGAAGTAGAAACTCCGGAAGAGAACACCGGTGACGTTATCGGTGACTTGAGCCGTCGTCGTGGTATGCTCAAAGGTCAGGAATCTGAAGTTAC
TGGCGTTAAGATCCACGCTGAAGTACCGCTGTCTGAAATGTTCGGATACGCAACTCAGCTGCGTTCTCTGACCAAAGGTCGTGCATCATACACTATG
GAATTCCTGAAGTATGATGAAGCGCCGAGTAACGTTGCTCAGGCCGTAATTGAAGCCCGTGGTAAAAGATCTCATCACCATCACCATCACTAATGAg
atccggctgctaacaaagcccgaaaggaagctgagttggctgctgccaccgctgagcaataactagcataaccccttggggcctctaaacgggtctt
gaggggttttttgctgaaaggaggaactatatccggattggcgaatgggacgcgccctgtagcggcgcattaagcgcggcgggtgtggtggttacgc
gcagcgtgaccgctacacttgccagcgccctagcgcccgctcctttcgctttcttcccttcctttctcgccacgttcgccggctttccccgtcaagc
tctaaatcgggggctccctttagggttccgatttagtgctttacggcacctcgaccccaaaaaacttgattagggtgatggttcacgtagtgggcca
tcgccctgatagacggtttttcgccctttgacgttggagtccacgttctttaatagtggactcttgttccaaactggaacaacactcaaccctatct
cggtctattcttttgatttataagggattttgccgatttcggcctattggttaaaaaatgagctgatttaacaaaaatttaacgcgaattttaacaa
aatattaacgcttacaatttaggtggcacttttcggggaaatgtgcgcggaacccctatttgtttatttttctaaatacattcaaatatgtatccgc
tcatgagacaataaccctgataaatgcttcaataatattgaaaaaggaagagtatgagtattcaacatttccgtgtcgcccttattcccttttttgc
ggcattttgccttcctgtttttgctcacccagaaacgctggtgaaagtaaaagatgctgaagatcagttgggtgcacgagtgggttacatcgaactg
gatctcaacagcggtaagatccttgagagttttcgccccgaagaacgttttccaatgatgagcacttttaaagttctgctatgtggcgcggtattat
cccgtattgacgccgggcaagagcaactcggtcgccgcatacactattctcagaatgacttggttgagtactcaccagtcacagaaaagcatcttac
ggatggcatgacagtaagagaattatgcagtgctgccataaccatgagtgataacactgcggccaacttacttctgacaacgatcggaggaccgaag
gagctaaccgcttttttgcacaacatgggggatcatgtaactcgccttgatcgttgggaaccggagctgaatgaagccataccaaacgacgagcgtg
acaccacgatgcctgcagcaatggcaacaacgttgcgcaaactattaactggcgaactacttactctagcttcccggcaacaattaatagactggat
ggaggcggataaagttgcaggaccacttctgcgctcggcccttccggctggctggtttattgctgataaatctggagccggtgagcgtgggtctcgc
ggtatcattgcagcactggggccagatggtaagccctcccgtatcgtagttatctacacgacggggagtcaggcaactatggatgaacgaaatagac
agatcgctgagataggtgcctcactgattaagcattggtaactgtcagaccaagtttactcatatatactttagattgatttaaaacttcatttta
atttaaaaggatctaggtgaagatcctttttgataatctcatgaccaaaatcccttaacgtgagttttcgttccactgagcgtcagaccccgtagaa
aagatcaaaggatcttcttgagatcctttttttctgcgcgtaatctgctgcttgcaaacaaaaaaaccaccgctaccagcggtggtttgtttgccgg
atcaagagctaccaactctttttccgaaggtaactggcttcagcagagcgcagataccaaatactgtccttctagtgtagccgtagttaggccacca
cttcaagaactctgtagcaccgcctacatacctcgctctgctaatcctgttaccagtggctgctgccagtggcgataagtcgtgtcttaccgggttg
gactcaagacgatagttaccggataaggcgcagcggtcgggctgaacggggggttcgtgcacacagcccagcttggagcgaacgacctacaccgaac
tgagatacctacagcgtgagctatgagaaagcgccacgcttcccgaagggagaaaggcggacaggtatccggtaagcggcagggtcggaacaggaga
gcgcacgagggagcttccagggggaaacgcctggtatctttatagtcctgtcgggtttcgccacctctgacttgagcgtcgatttttgtgatgctcg
tcaggggggcggagcctatggaaaaacgccagcaacgcggcctttttacggttcctggccttttgctggccttttgctcacatgttctttcctgcgt
tatcccctgattctgtggataaccgtattaccgcctttgagtgagctgataccgctcgccgcagccgaacgaccgagcgcagcgagtcagtgagcga
ggaagcggaagagcgcctgatgcggtattttctccttacgcatctgtgcggtatttcacaccgcaatggtgcactctcagtacaatctgctctgatg
ccgcatagttaagccagtatacactccgctatcgctacgtgactgggtcatggctgcgccccgacacccgccaacacccgctgacgcgccctgacgg
gcttgtctgctcccggcatccgcttacagacaagctgtgaccgtctccgggagctgcatgtgtcagaggttttcaccgtcatcaccgaaacgcgcga
ggcagctgcggtaaagctcatcagcgtggtcgtgaagcgattcacagatgtctgcctgttcatccgcgtccagctcgttgagtttctccagaagcgt
taatgtctggcttctgataaagcgggccatgttaagggcggttttttcctgtttggtcactgatgcctccgtgtaagggggatttctgttcatgggg
gtaatgataccgatgaaacgagagaggatgctcacgatacgggttactgatgatgaacatgcccggttactggaacgttgtgagggtaaacaactgg
cggtatggatgcggcgggaccagagaaaaatcactcagggtcaatgccagcgcttcgttaatacagatgtaggtgttccacagggtagccagcagca
tcctgcgatgcagatccggaacataatggtgcagggcgctgacttccgcgtttccagactttacgaaacacggaaaccgaagaccattcatgttgtt

gctcaggtcgcagacgttttgcagcagcagtcgcttcacgttcgctcgcgtatcggtgattcattctgctaaccagtaaggcaaccccgccagccta
gccgggtcctcaacgacaggagcacgatcatgcgcacccgtggggccgccatgccggcgataatggcctgcttctcgccgaaacgtttggtggcggg
accagtgacgaaggcttgagcgagggcgtgcaagattccgaataccgcaagcgacaggccgatcatcgtcgcgctccagcgaaagcggtcctcgccg
aaaatgacccagagcgctgccggcacctgtcctacgagttgcatgataaagaagacagtcataagtgcggcgacgatagtcatgccccgcgcccacc
ggaaggagctgactgggttgaaggctctcaagggcatcggtcgagatcccggtgcctaatgagtgagctaacttacattaattgcgttgcgctcact
gcccgctttccagtcgggaaacctgtcgtgccagctgcattaatgaatcggccaacgcgcggggagaggcggtttgcgtattgggcgccagggtggt
ttttcttttcaccagtgagacgggcaacagctgattgcccttcaccgcctggccctgagagagttgcagcaagcggtccacgctggtttgccccagc
aggcgaaaatcctgtttgatggtggttaacggcgggatataacatgagctgtcttcggtatcgtcgtatcccactaccgagatatccgcaccaacgc
gcagcccggactcggtaatggcgcgcattgcgcccagcgccatctgatcgttggcaaccagcatcgcagtgggaacgatgccctcattcagcatttg
catggtttgttgaaaaccggacatggcactccagtcgccttcccgttccgctatcggctgaatttgattgcgagtgagatatttatgccagccagcc
agacgcagacgcgccgagacagaacttaatgggcccgctaacagcgcgatttgctggtgacccaatgcgaccagatgctccacgcccagtcgcgtac
cgtcttcatgggagaaaataatactgttgatgggtgtctggtcagagacatcaagaaataacgccggaacattagtgcaggcagcttccacagcaat
ggcatcctggtcatccagcggatagttaatgatcagcccactgacgcgttgcgcgagaagattgtgcaccgccgctttacaggcttcgacgccgctt
cgttctaccatcgacaccaccacgctggcacccagttgatcggcgcgagatttaatcgccgcgacaatttgcgacggcgcgtgcagggccagactgg
aggtggcaacgccaatcagcaacgactgtttgcccgccagttgttgtgccacgcggttgggaatgtaattcagctccgccatcgccgcttccacttt
ttcccgcgttttcgcagaaacgtggctggcctggttcaccacgcgggaaacggtctgataagagacaccggcatactctgcgacatcgtataacgtt
actggtttcacattcaccaccctgaattgactctcttccgggcgctatcatgccataccgcgaaaggttttgcgccattcgatggtgtccgggatct
cgacgctctcccttatgcgactcctgcattaggaagcagcccagtagtaggttgaggccgttgagcaccgccgccgcaaggaatggtgcatgcaagg
agatggcgcccaacagtcccccggccacggggcctgccaccatacccacgccgaaacaagcgctcatgagcccgaagtggcgagcccgatcttcccc
atcggtgatgtcggcgatataggcgccagcaaccgcacctgtggcgccggtgatgccggccacgatgcgtccggcgtagaggatcgagatctcgatc
ccgcgaaat

### *Sequence of pET21a-EFTu-His*

#### $P_{T7}$ promoter – EFTu-His6 - T7 terminator

taatacgactcactataggggaattgtgagcggataacaattcccctctagaaataattttgtttaactttaagaaggagatatacatATGTCTAAA
GAAAAGTTTGAACGTACAAAACCGCACGTTAACGTCGGTACTATCGGCCACGTTGACCATGGTAAAACAACGCTGACCGCTGCAATCACTACCGTAC
TGGCTAAAACCTACGGCGGTGCTGCTCGCGCATTCGACCAGATCGATAACGCGCCGGAAGAAAAAGCTCGTGGTATCACCATCAACACTTCTCACGT
TGAATACGACACCCCGACCCGTCACTACGCACACGTAGACTGCCCGGGGCACGCCGACTATGTTAAAAACATGATCACCGGTGCTGCGCAGATGGAC
GGCGCGATCCTGGTAGTTGCTGCGACTGACGGCCCGATGCCGCAGACTCGTGAGCACATCCTGCTGGGTCGTCAGGTAGGCGTTCCGTACATCATCG
TGTTCCTGAACAAATGCGACATGGTTGATGACGAAGAGCTGCTGGAACTGGTTGAAATGGAAGTTCGTGAACTTCTGTCTCAGTACGACTTCCCGGG
CGACGACACTCCGATCGTTCGTGGTTCTGCTCTGAAAGCGCTGGAAGGCGACGCAGAGTGGGAAGCGAAAATCCTGGAACTGGCTGGCTTCCTGGAT
TCTTACATTCCGGAACCAGAGCGTGCGATTGACAAGCCGTTCCTGCTGCCGATCGAAGACGTATTCTCCATCTCCGGTCGTGGTACCGTTGTTACCG
GTCGTGTAGAACGCGGTATCATCAAAGTTGGTGAAGAAGTTGAAATCGTTGGTATCAAAGAGACTCAGAAGTCTACCTGTACTGGCGTTGAAATGTT
CCGCAAACTGCTGGACGAAGGCCGTGCTGGTGAGAACGTAGGTGTTCTGCTGCGTGGTATCAAACGTGAAGAAATCGAACGTGGTCAGGTACTGGCT
AAGCCGGGCACCATCAAGCCGCACACCAAGTTCGAATCTGAAGTGTACATTCTGTCCAAAGATGAAGGCGGCCGTCATACTCCGTTCTTCAAAGGCT
ACCGTCCGCAGTTCTACTTCCGTACTACTGACGTGACTGGTACCATCGAACTGCCGGAAGGCGTAGAGATGGTAATGCCGGGCGACAACATCAAAAT
GGTTGTTACCCTGATCCACCCGATCGCGATGGACGACGGTCTGCGTTTCGCAATCCGTGAAGGCGGCCGTACCGTTGGCGCGGGCGTTGTAGCAAAA
GTTCTGGGCAGATCTCATCACCATCACCATCACTAATGAgatccggctgctaacaaagcccgaaaggaagctgagttggctgctgccaccgctgagc
aataactagcataaccccttggggcctctaaacgggtcttgaggggttttttgctgaaaggaggaactatatccggattggcgaatgggacgcgccc
tgtagcggcgcattaagcgcggcgggtgtggtggttacgcgcagcgtgaccgctacacttgccagcgccctagcgcccgctcctttcgctttcttcc
cttcctttctcgccacgttcgccggctttccccgtcaagctctaaatcgggggctccctttagggttccgatttagtgctttacggcacctcgaccc
caaaaaacttgattagggtgatggttcacgtagtgggccatcgccctgatagacggtttttcgccctttgacgttggagtccacgttctttaatagt
ggactcttgttccaaactggaacaacactcaaccctatctcggtctattcttttgatttataagggattttgccgatttcggcctattggttaaaaa
atgagctgatttaacaaaaatttaacgcgaattttaacaaaatattaacgcttacaatttaggtggcacttttcggggaaatgtgcgcggaacccct
atttgtttatttttctaaatacattcaaatatgtatccgctcatgagacaataaccctgataaatgcttcaataatattgaaaaaggaagagtatga
gtattcaacatttccgtgtcgcccttattcccttttttgcggcattttgccttcctgtttttgctcacccagaaacgctggtgaaagtaaaagatgc
tgaagatcagttgggtgcacgagtgggttacatcgaactggatctcaacagcggtaagatccttgagagttttcgccccgaagaacgttttccaatg
atgagcacttttaaagttctgctatgtggcgcggtattatcccgtattgacgccgggcaagagcaactcggtcgccgcatacactattctcagaatg
acttggttgagtactcaccagtcacagaaaagcatcttacggatggcatgacagtaagagaattatgcagtgctgccataaccatgagtgataacac
tgcggccaacttacttctgacaacgatcggaggaccgaaggagctaaccgcttttttgcacaacatgggggatcatgtaactcgccttgatcgttgg
gaaccggagctgaatgaagccataccaaacgacgagcgtgacaccacgatgcctgcagcaatggcaacaacgttgcgcaaactattaactggcgaac
tacttactctagcttcccggcaacaattaatagactggatggaggcggataaagttgcaggaccacttctgcgctcggcccttccggctggctggtt
tattgctgataaatctggagccggtgagcgtgggtctcgcggtatcattgcagcactggggccagatggtaagccctcccgtatcgtagttatctac
acgacggggagtcaggcaactatggatgaacgaaatagacagatcgctgagataggtgcctcactgattaagcattggtaactgtcagaccaagttt
actcatatatactttagattgatttaaaacttcatttttaatttaaaaggatctaggtgaagatcctttttgataatctcatgaccaaaatccctta
acgtgagttttcgttccactgagcgtcagaccccgtagaaaagatcaaaggatcttcttgagatcctttttttctgcgcgtaatctgctgcttgcaa
acaaaaaaaccaccgctaccagcggtggtttgtttgccggatcaagagctaccaactctttttccgaaggtaactggcttcagcagagcgcagatac
caaatactgtccttctagtgtagccgtagttaggccaccacttcaagaactctgtagcaccgcctacatacctcgctctgctaatcctgttaccagt
ggctgctgccagtggcgataagtcgtgtcttaccgggttggactcaagacgatagttaccggataaggcgcagcggtcgggctgaacggggggttcg
tgcacacagcccagcttggagcgaacgacctacaccgaactgagatacctacagcgtgagctatgagaaagcgccacgcttcccgaagggagaaagg
cggacaggtatccggtaagcggcagggtcggaacaggagagcgcacgagggagcttccagggggaaacgcctggtatctttatagtcctgtcgggtt
tcgccacctctgacttgagcgtcgatttttgtgatgctcgtcaggggggcggagcctatggaaaaacgccagcaacgcggcctttttacggttcctg
gccttttgctggccttttgctcacatgttctttcctgcgttatcccctgattctgtggataaccgtattaccgcctttgagtgagctgataccgctc
gccgcagccgaacgaccgagcgcagcgagtcagtgagcgaggaagcggaagagcgcctgatgcggtattttctccttacgcatctgtgcggtatttc
acaccgcaatggtgcactctcagtacaatctgctctgatgccgcatagttaagccagtatacactccgctatcgctacgtgactgggtcatggctgc
gccccgacacccgccaacacccgctgacgcgccctgacgggcttgtctgctcccggcatccgcttacagacaagctgtgaccgtctccgggagctgc
atgtgtcagaggttttcaccgtcatcaccgaaacgcgcgaggcagctgcggtaaagctcatcagcgtggtcgtgaagcgattcacagatgtctgcct
gttcatccgcgtccagctcgttgagtttctccagaagcgttaatgtctggcttctgataaagcgggccatgttaagggcggttttttcctgtttggt
cactgatgcctccgtgtaagggggatttctgttcatgggggtaatgataccgatgaaacgagagaggatgctcacgatacgggttactgatgatgaa
catgcccggttactggaacgttgtgagggtaaacaactggcggtatggatgcggcgggaccagagaaaaatcactcagggtcaatgccagcgcttcg
ttaatacagatgtaggtgttccacagggtagccagcagcatcctgcgatgcagatccggaacataatggtgcagggcgctgacttccgcgtttccag
actttacgaaacacggaaaccgaagaccattcatgttgttgctcaggtcgcagacgttttgcagcagcagtcgcttcacgttcgctcgcgtatcggt
gattcattctgctaaccagtaaggcaaccccgccagcctagccgggtcctcaacgacaggagcacgatcatgcgcacccgtgggccgccatgccgg
cgataatggcctgcttctcgccgaaacgtttggtggcgggaccagtgacgaaggcttgagcgagggcgtgcaagattccgaataccgcaagcgacag
gccgatcatcgtcgcgctccagcgaaagcggtcctcgccgaaaatgacccagagcgctgccggcacctgtcctacgagttgcatgataaagaagaca
gtcataagtgcggcgacgatagtcatgccccgcgcccaccggaaggagctgactgggttgaaggctctcaagggcatcggtcgagatcccggtgcct
aatgagtgagctaacttacattaattgcgttgcgctcactgcccgctttccagtcgggaaacctgtcgtgccagctgcattaatgaatcggccaacg
cgcggggagaggcggtttgcgtattgggcgccagggtggtttttcttttcaccagtgagacgggcaacagctgattgcccttcaccgcctggccctg
agagagttgcagcaagcggtccacgctggtttgccccagcaggcgaaaatcctgtttgatggtggttaacggcgggatataacatgagctgtcttcg
gtatcgtcgtatcccactaccgagatatccgcaccaacgcgcagcccggactcggtaatggcgcgcattgcgcccagcgccatctgatcgttggcaa
ccagcatcgcagtgggaacgatgccctcattcagcatttgcatggtttgttgaaaaccggacatggcactccagtcgccttcccgttccgctatcgg

ctgaatttgattgcgagtgagatatttatgccagccagccagacgcagacgcgccgagacagaacttaatgggcccgctaacagcgcgatttgctgg
tgacccaatgcgaccagatgctccacgcccagtcgcgtaccgtcttcatgggagaaaataatactgttgatgggtgtctggtcagagacatcaagaa
ataacgccggaacattagtgcaggcagcttccacagcaatggcatcctggtcatccagcggatagttaatgatcagcccactgacgcgttgcgcgag
aagattgtgcaccgccgctttacaggcttcgacgccgcttcgttctaccatcgacaccaccacgctggcacccagttgatcggcgcgagatttaatc
gccgcgacaatttgcgacggcgcgtgcagggccagactggaggtggcaacgccaatcagcaacgactgtttgcccgccagttgttgtgccacgcggt
tgggaatgtaattcagctccgccatcgccgcttccactttttcccgcgttttcgcagaaacgtggctggcctggttcaccacgcgggaaacggtctg
ataagagacaccggcatactctgcgacatcgtataacgttactggtttcacattcaccaccctgaattgactctcttccgggcgctatcatgccata
ccgcgaaaggttttgcgccattcgatggtgtccgggatctcgacgctctcccttatgcgactcctgcattaggaagcagcccagtagtaggttgagg
ccgttgagcaccgccgccgcaaggaatggtgcatgcaaggagatggcgcccaacagtcccccggccacggggcctgccaccatacccacgccgaaac
aagcgctcatgagcccgaagtggcgagcccgatcttccccatcggtgatgtcggcgatataggcgccagcaaccgcacctgtggcgccggtgatgcc
ggccacgatgcgtccggcgtagaggatcgagatctcgatcccgcgaaat

### *Sequence of pET21a-EFTs-His*

#### $P_{T7}$ promoter – EFTs-His6 - T7 terminator

taatacgactcactataggggaattgtgagcggataacaattcccctctagaaataattttgtttaactttaagaaggagatatacatATGGCTGAA
ATTACCGCATCCCTGGTAAAAGAGCTGCGTGAGCGTACTGGCGCAGGCATGATGGATTGCAAAAAAGCACTGACTGAAGCTAACGGCGACATCGAGC
TGGCAATCGAAAACATGCGTAAGTCCGGTGCTATTAAAGCAGCGAAAAAAGCAGGCAACGTTGCTGCTGACGGCGTGATCAAAACCAAAATCGACGG
CAACTACGGCATCATTCTGGAAGTTAACTGCCAGACTGACTTCGTTGCAAAAGACGCTGGTTTCCAGGCGTTCGCAGACAAAGTTCTGGACGCAGCT
GTTGCTGGCAAAATCACTGACGTTGAAGTTCTGAAAGCACAGTTCGAAGAAGAACGTGTTGCGCTGGTAGCGAAAATTGGTGAAAACATCAACATTC
GCCGCGTTGCTGCGCTGGAAGGCGACGTTCTGGGTTCTTATCAGCACGGTGCGCGTATCGGCGTTCTGGTTGCTGCTAAAGGCGCTGACGAAGAGCT
GGTTAAACACATCGCTATGCACGTTGCTGCAAGCAAGCCAGAATTCATCAAACCGGAAGACGTATCCGCTGAAGTGGTAGAAAAAGAATACCAGGTA
CAGCTGGATATCGCGATGCAGTCTGGTAAGCCGAAAGAAATCGCAGAGAAAATGGTTGAAGGCCGCATGAAGAAATTCACCGGCGAAGTTTCTCTGA
CCGGTCAGCCGTTCGTTATGGAACCAAGCAAAACTGTTGGTCAGCTGCTGAAAGAGCATAACGCTGAAGTGACTGGCTTCATCCGCTTCGAAGTGGG
TGAAGGCATCGAGAAAGTTGAGACTGACTTTGCAGCAGAAGTTGCTGCGATGTCCAAGCAGTCTAGATCTCATCACCATCACCATCACTAATGAgat
ccggctgctaacaaagcccgaaaggaagctgagttggctgctgccaccgctgagcaataactagcataaccccttggggcctctaaacgggtcttga
ggggttttttgctgaaaggaggaactatatccggattggcgaatgggacgcgccctgtagcggcgcattaagcgcggcgggtgtggtggttacgcgc
agcgtgaccgctacacttgccagcgccctagcgcccgctcctttcgctttcttcccttcctttctcgccacgttcgccggctttccccgtcaagctc
taaatcgggggctccctttagggttccgatttagtgctttacggcacctcgaccccaaaaaacttgattagggtgatggttcacgtagtgggccatc
gccctgatagacggtttttcgccctttgacgttggagtccacgttctttaatagtggactcttgttccaaactggaacaacactcaaccctatctcg
gtctattcttttgatttataagggattttgccgatttcggcctattggttaaaaaatgagctgatttaacaaaaatttaacgcgaattttaacaaaa
tattaacgcttacaatttaggtggcacttttcggggaaatgtgcgcggaacccctatttgtttatttttctaaatacattcaaatatgtatccgctc
atgagacaataaccctgataaatgcttcaataatattgaaaaaggaagagtatgagtattcaacatttccgtgtcgcccttattcccttttttgcgg
cattttgccttcctgtttttgctcacccagaaacgctggtgaaagtaaaagatgctgaagatcagttgggtgcacgagtgggttacatcgaactgga
tctcaacagcggtaagatccttgagagttttcgccccgaagaacgttttccaatgatgagcacttttaaagttctgctatgtggcgcggtattatcc
cgtattgacgccgggcaagagcaactcggtcgccgcatacactattctcagaatgacttggttgagtactcaccagtcacagaaaagcatcttacgg
atggcatgacagtaagagaattatgcagtgctgccataaccatgagtgataacactgcggccaacttacttctgacaacgatcggaggaccgaagga
gctaaccgcttttttgcacaacatgggggatcatgtaactcgccttgatcgttgggaaccggagctgaatgaagccataccaaacgacgagcgtgac
accacgatgcctgcagcaatggcaacaacgttgcgcaaactattaactggcgaactacttactctagcttcccggcaacaattaatagactggatgg
aggcggataaagttgcaggaccacttctgcgctcggcccttccggctggctggtttattgctgataaatctggagccggtgagcgtgggtctcgcgg
tatcattgcagcactggggccagatggtaagccctcccgtatcgtagttatctacacgacggggagtcaggcaactatggatgaacgaaatagacag
atcgctgagataggtgcctcactgattaagcattggtaactgtcagaccaagtttactcatatatactttagattgatttaaaacttcatttttaat
ttaaaaggatctaggtgaagatcctttttgataatctcatgaccaaaatcccttaacgtgagttttcgttccactgagcgtcagaccccgtagaaaa
gatcaaaggatcttcttgagatcctttttttctgcgcgtaatctgctgcttgcaaacaaaaaaaccaccgctaccagcggtggtttgtttgccggat
caagagctaccaactctttttccgaaggtaactggcttcagcagagcgcagataccaaatactgtccttctagtgtagccgtagttaggccaccact
tcaagaactctgtagcaccgcctacatacctcgctctgctaatcctgttaccagtggctgctgccagtggcgataagtcgtgtcttaccgggttgga
ctcaagacgatagttaccggataaggcgcagcggtcgggctgaacggggggttcgtgcacacagcccagcttggagcgaacgacctacaccgaactg
agatacctacagcgtgagctatgagaaagcgccacgcttcccgaagggagaaaggcggacaggtatccggtaagcggcagggtcggaacaggagagc
gcacgagggagcttccagggggaaacgcctggtatctttatagtcctgtcgggtttcgccacctctgacttgagcgtcgatttttgtgatgctcgtc
aggggggcggagcctatggaaaaacgccagcaacgcggcctttttacggttcctggccttttgctggccttttgctcacatgttctttcctgcgtta
tcccctgattctgtggataaccgtattaccgcctttgagtgagctgataccgctcgccgcagccgaacgaccgagcgcagcgagtcagtgagcgagg
aagcggaagagcgcctgatgcggtattttctccttacgcatctgtgcggtatttcacaccgcaatggtgcactctcagtacaatctgctctgatgcc
gcatagttaagccagtatacactccgctatcgctacgtgactgggtcatggctgcgccccgacacccgccaacacccgctgacgcgccctgacgggc
ttgtctgctcccggcatccgcttacagacaagctgtgaccgtctccgggagctgcatgtgtcagaggttttcaccgtcatcaccgaaacgcgcgagg
cagctgcggtaaagctcatcagcgtggtcgtgaagcgattcacagatgtctgcctgttcatccgcgtccagctcgttgagtttctccagaagcgtta
atgtctggcttctgataaagcgggccatgttaagggcggttttttcctgtttggtcactgatgcctccgtgtaagggggatttctgttcatgggggt
aatgataccgatgaaacgagagaggatgctcacgatacgggttactgatgatgaacatgcccggttactggaacgttgtgagggtaaacaactggcg
gtatggatgcggcgggaccagagaaaaatcactcagggtcaatgccagcgcttcgttaatacagatgtaggtgttccacagggtagccagcagcatc
ctgcgatgcagatccggaacataatggtgcagggcgctgacttccgcgtttccagactttacgaaacacggaaaccgaagaccattcatgttgttgc
tcaggtcgcagacgttttgcagcagcagtcgcttcacgttcgctcgcgtatcggtgattcattctgctaaccagtaaggcaaccccgccagcctagc
cgggtcctcaacgacaggagcacgatcatgcgcacccgtggggccgccatgccggcgataatggcctgcttctcgccgaaacgtttggtggcgggac
cagtgacgaaggcttgagcgagggcgtgcaagattccgaataccgcaagcgacaggccgatcatcgtcgcgctccagcgaaagcggtcctcgccgaa
aatgacccagagcgctgccggcacctgtcctacgagttgcatgataaagaagacagtcataagtgcggcgacgatagtcatgccccgcgcccaccgg
aaggagctgactgggttgaaggctctcaagggcatcggtcgagatcccggtgcctaatgagtgagctaacttacattaattgcgttgcgctcactgc
ccgctttccagtcgggaaacctgtcgtgccagctgcattaatgaatcggccaacgcgcggggagaggcggtttgcgtattgggcgccagggtggttt
ttcttttcaccagtgagacgggcaacagctgattgcccttcaccgcctggccctgagagagttgcagcaagcggtccacgctggtttgccccagcag
gcgaaaatcctgtttgatggtggttaacggcgggatataacatgagctgtcttcggtatcgtcgtatcccactaccgagatatccgcaccaacgcgc
agcccggactcggtaatggcgcgcattgcgcccagcgccatctgatcgttggcaaccagcatcgcagtgggaacgatgccctcattcagcatttgca
tggtttgttgaaaaccggacatggcactccagtcgccttcccgttccgctatcggctgaatttgattgcgagtgagatatttatgccagccagccag
acgcagacgcgccgagacagaacttaatgggcccgctaacagcgcgatttgctggtgacccaatgcgaccagatgctccacgcccagtcgcgtaccg
tcttcatgggagaaaataatactgttgatgggtgtctggtcagagacatcaagaaataacgccggaacattagtgcaggcagcttccacagcaatgg
catcctggtcatccagcggatagttaatgatcagcccactgacgcgttgcgcgagaagattgtgcaccgccgctttacaggcttcgacgccgcttcg

ttctaccatcgacaccaccacgctggcacccagttgatcggcgcgagatttaatcgccgcgacaatttgcgacggcgcgtgcagggccagactggag
gtggcaacgccaatcagcaacgactgtttgcccgccagttgttgtgccacgcggttgggaatgtaattcagctccgccatcgccgcttccactttttt
cccgcgttttcgcagaaacgtggctggcctggttcaccacgcgggaaacggtctgataagagacaccggcatactctgcgacatcgtataacgttac
tggtttcacattcaccaccctgaattgactctcttccgggcgctatcatgccataccgcgaaaggttttgcgccattcgatggtgtccgggatctcg
acgctctcccttatgcgactcctgcattaggaagcagcccagtagtaggttgaggccgttgagcaccgccgccgcaaggaatggtgcatgcaaggag
atggcgcccaacagtcccccggccacggggcctgccaccatacccacgccgaaacaagcgctcatgagcccgaagtggcgagcccgatcttccccat
cggtgatgtcggcgatataggcgccagcaaccgcacctgtggcgccggtgatgccggccacgatgcgtccggcgtagaggatcgagatctcgatccc
gcgaaat

### *Sequence of pET21a-RF1-His*

#### $P_{T7}$ promoter – RF1-His6 - T7 terminator

taatacgactcactataggggaattgtgagcggataacaattcccctctagaaataattttgtttaactttaagaaggagatatacatATGAAGCCT
TCTATCGTTGCCAAACTGGAAGCCCTGCATGAACGCCATGAAGAAGTTCAGGCGTTGCTGGGTGACGCGCAAACTATCGCCGACCAGGAACGTTTTC
GCGCATTATCACGCGAATATGCGCAGTTAAGTGATGTTTCGCGCTGTTTTACCGACTGGCAACAGGTTCAGGAAGATATCGAAACCGCACAGATGAT
GCTCGATGATCCTGAAATGCGTGAGATGGCGCAGGATGAACTGCGCGAAGCTAAAGAAAAAAGCGAGCAACTGGAACAGCAATTACAGGTTCTGTTA
CTGCCAAAAGATCCTGATGACGAACGTAACGCCTTCCTCGAAGTCCGAGCCGGAACCGGCGGCGACGAAGCGGCGCTGTTCGCGGGCGATCTGTTCC
GTATGTACAGCCGTTATGCCGAAGCCCGCCGCTGGCGGGTAGAAATCATGAGCGCCAGCGAGGGTGAACATGGTGGTTATAAAGAGATCATCGCCAA
AATTAGCGGTGATGGTGTGTATGGTCGTCTGAAATTTGAATCCGGCGGTCATCGCGTGCAACGTGTTCCTGCTACGGAATCGCAGGGTCGTATTCAT
ACTTCTGCTTGTACCGTTGCGGTAATGCCAGAACTGCCTGACGCAGAACTGCCGGACATCAACCCAGCAGATTTACGCATTGATACTTTCCGCTCGT
CAGGGGCGGGTGGTCAGCACGTTAACACCACCGATTCGGCAATTCGTATTACTCACTTGCCGACCGGGATTGTTGTTGAATGTCAGGACGAACGTTC
ACAACATAAAAACAAAGCTAAAGCACTTTCTGTTCTCGGTGCTCGCATCCACGCTGCTGAAATGGCAAAACGCCAACAGGCCGAAGCGTCTACCCGT
CGTAACCTGCTGGGGAGTGGCGATCGCAGCGACCGTAACCGTACTTACAACTTCCCGCAGGGGCGCGTTACCGATCACCGCATCAACCTGACGCTCT
ACCGCCTGGATGAAGTGATGGAAGGTAAGCTGGATATGCTGATTGAACCGATTATCCAGGAACATCAGGCCGACCAACTGGCGGCGTTGTCCGAGCA
GGAACTCGAGCACCACCACCACCACCACTGAgatccggctgctaacaaagcccgaaaggaagctgagttggctgctgccaccgctgagcaataacta
gcataaccccttggggcctctaaacgggtcttgaggggttttttgctgaaaggaggaactatatccggattggcgaatgggacgcgccctgtagcgg
cgcattaagcgcggcgggtgtggtggttacgcgcagcgtgaccgctacacttgccagcgccctagcgcccgctcctttcgctttcttcccttccttt
ctcgccacgttcgccggctttccccgtcaagctctaaatcgggggctccctttagggttccgatttagtgctttacggcacctcgaccccaaaaaac
ttgattagggtgatggttcacgtagtgggccatcgccctgatagacggtttttcgccctttgacgttggagtccacgttctttaatagtggactctt
gttccaaactggaacaacactcaaccctatctcggtctattcttttgatttataagggattttgccgatttcggcctattggttaaaaaatgagctg
atttaacaaaaatttaacgcgaattttaacaaaatattaacgcttacaatttaggtggcacttttcggggaaatgtgcgcggaacccctatttgttt
atttttctaaatacattcaaatatgtatccgctcatgagacaataaccctgataaatgcttcaataatattgaaaaaggaagagtatgagtattcaa
catttccgtgtcgcccttattcccttttttgcggcattttgccttcctgtttttgctcacccagaaacgctggtgaaagtaaaagatgctgaagatc
agttgggtgcacgagtgggttacatcgaactggatctcaacagcggtaagatccttgagagttttcgccccgaagaacgttttccaatgatgagcac
ttttaaagttctgctatgtggcgcggtattatcccgtattgacgccgggcaagagcaactcggtcgccgcatacactattctcagaatgacttggtt
gagtactcaccagtcacagaaaagcatcttacggatggcatgacagtaagagaattatgcagtgctgccataaccatgagtgataacactgcggcca
acttacttctgacaacgatcggaggaccgaaggagctaaccgcttttttgcacaacatgggggatcatgtaactcgccttgatcgttgggaaccgga
gctgaatgaagccataccaaacgacgagcgtgacaccacgatgcctgcagcaatggcaacaacgttgcgcaaactattaactggcgaactacttact
ctagcttcccggcaacaattaatagactggatggaggcggataaagttgcaggaccacttctgcgctcggcccttccggctggctggtttattgctg
ataaatctggagccggtgagcgtgggtctcgcggtatcattgcagcactggggccagatggtaagccctcccgtatcgtagttatctacacgacggg
gagtcaggcaactatggatgaacgaaatagacagatcgctgagataggtgcctcactgattaagcattggtaactgtcagaccaagtttactcatat
atactttagattgatttaaaacttcatttttaatttaaaaggatctaggtgaagatcctttttgataatctcatgaccaaaatcccttaacgtgagt
tttcgttccactgagcgtcagaccccgtagaaaagatcaaaggatcttcttgagatcctttttttctgcgcgtaatctgctgcttgcaaacaaaaaa
accaccgctaccagcggtggtttgtttgccggatcaagagctaccaactctttttccgaaggtaactggcttcagcagagcgcagataccaaatact
gtccttctagtgtagccgtagttaggccaccacttcaagaactctgtagcaccgcctacatacctcgctctgctaatcctgttaccagtggctgctg
ccagtggcgataagtcgtgtcttaccgggttggactcaagacgatagttaccggataaggcgcagcggtcgggctgaacggggggttcgtgcacaca
gcccagcttggagcgaacgacctacaccgaactgagatacctacagcgtgagctatgagaaagcgccacgcttcccgaagggagaaaggcggacagg
tatccggtaagcggcagggtcggaacaggagagcgcacgagggagcttccagggggaaacgcctggtatctttatagtcctgtcgggtttcgccacc
tctgacttgagcgtcgatttttgtgatgctcgtcaggggggcggagcctatggaaaaacgccagcaacgcggcctttttacggttcctggccttttg
ctggccttttgctcacatgttctttcctgcgttatcccctgattctgtggataaccgtattaccgcctttgagtgagctgataccgctcgccgcagc
cgaacgaccgagcgcagcgagtcagtgagcgaggaagcggaagagcgcctgatgcggtattttctccttacgcatctgtgcggtatttcacaccgca
atggtgcactctcagtacaatctgctctgatgccgcatagttaagccagtatacactccgctatcgctacgtgactgggtcatggctgcgccccgac
acccgccaacacccgctgacgcgccctgacgggcttgtctgctcccggcatccgcttacagacaagctgtgaccgtctccgggagctgcatgtgtca
gaggttttcaccgtcatcaccgaaacgcgcgaggcagctgcggtaaagctcatcagcgtggtcgtgaagcgattcacagatgtctgcctgttcatcc
gcgtccagctcgttgagtttctccagaagcgttaatgtctggcttctgataaagcgggccatgttaagggcggttttttcctgtttggtcactgatg
cctccgtgtaagggggatttctgttcatgggggtaatgataccgatgaaacgagagaggatgctcacgatacgggttactgatgatgaacatgcccg
gttactggaacgttgtgagggtaaacaactggcggtatggatgcggcgggaccagagaaaaatcactcagggtcaatgccagcgcttcgttaataca
gatgtaggtgttccacagggtagccagcagcatcctgcgatgcagatccggaacataatggtgcagggcgctgacttccgcgtttccagactttacg
aaacacggaaaccgaagaccattcatgttgttgctcaggtcgcagacgttttgcagcagcagtcgcttcacgttcgctcgcgtatcggtgattcatt
ctgctaaccagtaaggcaaccccgccagcctagccgggtcctcaacgacaggagcacgatcatgcgcacccgtgggccgccatgccggcgataatg
gcctgcttctcgccgaaacgtttggtggcgggaccagtgacgaaggcttgagcgagggcgtgcaagattccgaataccgcaagcgacaggccgatca
tcgtcgcgctccagcgaaagcggtcctcgccgaaaatgacccagagcgctgccggcacctgtcctacgagttgcatgataaagaagacagtcataag
tgcggcgacgatagtcatgccccgcgcccaccggaaggagctgactgggttgaaggctctcaagggcatcggtcgagatcccggtgcctaatgagtg
agctaacttacattaattgcgttgcgctcactgcccgctttccagtcgggaaacctgtcgtgccagctgcattaatgaatcggccaacgcgcgggga
gaggcggtttgcgtattgggcgccagggtggtttttcttttcaccagtgagacgggcaacagctgattgcccttcaccgcctggccctgagagagtt
gcagcaagcggtccacgctggtttgccccagcaggcgaaaatcctgtttgatggtggttaacggcgggatataacatgagctgtcttcggtatcgtc
gtatcccactaccgagatatccgcaccaacgcgcagcccggactcggtaatggcgcgcattgcgcccagcgccatctgatcgttggcaaccagcatc
gcagtgggaacgatgccctcattcagcatttgcatggtttgttgaaaaccggacatggcactccagtcgccttcccgttccgctatcggctgaattt
gattgcgagtgagatatttatgccagccagccagacgcagacgcgccgagacagaacttaatgggcccgctaacagcgcgatttgctggtgaccaa

tgcgaccagatgctccacgcccagtcgcgtaccgtcttcatgggagaaaataatactgttgatgggtgtctggtcagagacatcaagaaataacgcc
ggaacattagtgcaggcagcttccacagcaatggcatcctggtcatccagcggatagttaatgatcagcccactgacgcgttgcgcgagaagattgt
gcaccgccgctttacaggcttcgacgccgcttcgttctaccatcgacaccaccacgctggcacccagttgatcggcgcgagatttaatcgccgcgac
aatttgcgacggcgcgtgcagggccagactggaggtggcaacgccaatcagcaacgactgtttgcccgccagttgttgtgccacgcggttgggaatg
taattcagctccgccatcgccgcttccactttttcccgcgttttcgcagaaacgtggctggcctggttcaccacgcgggaaacggtctgataagaga
caccggcatactctgcgacatcgtataacgttactggtttcacattcaccaccctgaattgactctcttccgggcgctatcatgccataccgcgaaa
ggttttgcgccattcgatggtgtccgggatctcgacgctctcccttatgcgactcctgcattaggaagcagcccagtagtaggttgaggccgttgag
caccgccgccgcaaggaatggtgcatgcaaggagatggcgcccaacagtcccccggccacggggcctgccaccatacccacgccgaaacaagcgctc
atgagcccgaagtggcgagcccgatcttccccatcggtgatgtcggcgatataggcgccagcaaccgcacctgtggcgccggtgatgccggccacga
tgcgtccggcgtagaggatcgagatctcgatcccgcgaaat

### ***Sequence of pET21a-His-RF3-His***

$P_{T7}$ promoter – His6-RF3 - T7 terminator

```
taatacgactcactataggggaattgtgagcggataacaattcccctctagaaataattttgtttaactttaagaaggagatatacatATGAGAGGA
TCGCATCACCATCACCATCACGGATCCGATGACGATGACAAAATGACGTTGTCTCCTTATTTGCAAGAGGTGGCGAAGCGCCGCACTTTTGCCATTA
TTTCTCACCCGGACGCCGGTAAGACTACCATCACCGAGAAGGTGCTGCTGTTCGGACAGGCCATTCAGACCGCCGGTACAGTAAAAGGCCGTGGTTC
CAACCAGCACGCTAAGTCGGACTGGATGGAGATGGAAAAGCAGCGTGGGATCTCCATTACTACGTCTGTGATGCAGTTTCCGTATCACGATTGCCTG
GTTAACCTGCTCGACACCCCGGGGCACGAAGACTTCTCGGAAGATACCTATCGTACCCTGACGGCGGTGGACTGCTGCCTGATGGTTATCGACGCCG
CAAAAGGTGTTGAAGATCGTACCCGTAAGCTGATGGAAGTTACCCGTCTGCGCGACACGCCGATCCTCACCTTTATGAACAAACTTGACCGTGATAT
CCGCGACCCGATGGAGCTGCTCGATGAAGTTGAGAACGAGCTGAAAATCGGCTGTGCGCCGATCACCTGGCCGATTGGCTGCGGCAAGCTGTTTAAA
GGCGTTTACCACCTTTATAAAGACGAAACCTATCTCTATCAGAGCGGTAAAGGCCACACCATTCAGGAAGTCCGCATTGTTAAAGGGCTGAATAACC
CGGATCTCGATGCTGCGGTTGGTGAAGATCTGGCACAGCAGCTGCGTGACGAACTGGAACTGGTGAAAGGCGCGTCTAACGAGTTCGACAAAGAGCT
GTTCCTTGCGGGCGAAATCACTCCGGTATTCTTCGGTACTGCGCTGGGTAACTTCGGCGTCGATCATATGTTGGATGGCCTGGTGGAGTGGGCACCT
GCGCCGATGCCGCGTCAGACTGATACCCGTACCGTAGAAGCGAGCGAAGATAAATTTACCGGCTTCGTATTTAAAATTCAGGCCAACATGGACCCGA
AACACCGCGACCGCGTGGCGTTTATGCGTGTGGTGTCCGGTAAATATGAAAAAGGCATGAAACTGCGCCAGGTGCGCACTGCGAAAGATGTGGTGAT
CTCCGACGCGCTGACCTTTATGGCGGGTGACCGTTCGCACGTTGAAGAAGCGTATCCGGGCGATATCCTCGGCCTGCACAACCACGGCACCATTCAG
ATCGGCGACACCTTTACCCAGGGTGAGATGATGAAGTTCACCGGTATTCCGAACTTCGCACCAGAACTGTTCCGTCGTATCCGCCTGAAAGATCCGC
TGAAGCAAAAACAGCTGCTCAAAGGGCTGGTACAGCTTTCCGAAGAGGGCGCGGTGCAGGTGTTCCGTCCAATCTCCAACAACGATCTGATCGTTGG
TGCAGTTGGTGTGCTGCAGTTTGATGTGGTGGTAGCGCGCCTGAAGAGCGAATACAACGTTGAAGCAGTGTATGAGTCAGTCAACGTTGCCACTGCC
CGCTGGGTAGAATGTGCAGACGCGAAGAAATTCGAAGAGTTCAAGCGTAAGAACGAAAGCCAACTGGCGCTTGATGGCGGCGATAACCTCGCTTACA
TCGCTACCAGCATGGTCAACCTGCGCCTGGCACAGGAACGTTATCCGGACGTTCAGTTCCACCAGACCCGCGAGCATTAATGAgatccggctgctaa
caaagcccgaaaggaagctgagttggctgctgccaccgctgagcaataactagcataaccccttggggcctctaaacgggtcttgaggggttttttg
ctgaaaggaggaactatatccggattggcgaatgggacgcgccctgtagcggcgcattaagcgcggcgggtgtggtggttacgcgcagcgtgaccgc
tacacttgccagcgccctagcgcccgctcctttcgctttcttcccttcctttctcgccacgttcgccggctttccccgtcaagctctaaatcggggg
ctccctttagggttccgatttagtgctttacggcacctcgaccccaaaaaacttgattagggtgatggttcacgtagtgggccatcgccctgataga
cggtttttcgccctttgacgttggagtccacgttctttaatagtggactcttgttccaaactggaacaacactcaaccctatctcggtctattcttt
tgatttataagggattttgccgatttcggcctattggttaaaaaatgagctgatttaacaaaaatttaacgcgaattttaacaaaatattaacgctt
acaatttaggtggcacttttcggggaaatgtgcgcggaacccctatttgtttatttttctaaatacattcaaatatgtatccgctcatgagacaata
accctgataaatgcttcaataatattgaaaaaggaagagtatgagtattcaacatttccgtgtcgcccttattcccttttttgcggcattttgcctt
cctgtttttgctcacccagaaacgctggtgaaagtaaaagatgctgaagatcagttgggtgcacgagtgggttacatcgaactggatctcaacagcg
gtaagatccttgagagttttcgccccgaagaacgttttccaatgatgagcacttttaaagttctgctatgtggcgcggtattatcccgtattgacgc
cgggcaagagcaactcggtcgccgcatacactattctcagaatgacttggttgagtactcaccagtcacagaaaagcatcttacggatggcatgaca
gtaagagaattatgcagtgctgccataaccatgagtgataacactgcggccaacttacttctgacaacgatcggaggaccgaaggagctaaccgctt
ttttgcacaacatgggggatcatgtaactcgccttgatcgttgggaaccggagctgaatgaagccataccaaacgacgagcgtgacaccacgatgcc
tgcagcaatggcaacaacgttgcgcaaactattaactggcgaactacttactctagcttcccggcaacaattaatagactggatggaggcggataaa
gttgcaggaccacttctgcgctcggcccttccggctggctggtttattgctgataaatctggagccggtgagcgtgggtctcgcggtatcattgcag
cactggggccagatggtaagccctcccgtatcgtagttatctacacgacggggagtcaggcaactatggatgaacgaaatagacagatcgctgagat
aggtgcctcactgattaagcattggtaactgtcagaccaagtttactcatatatactttagattgatttaaaacttcatttttaatttaaaaggatc
taggtgaagatcctttttgataatctcatgaccaaaatcccttaacgtgagttttcgttccactgagcgtcagaccccgtagaaaagatcaaaggat
cttcttgagatcctttttttctgcgcgtaatctgctgcttgcaaacaaaaaaaccaccgctaccagcggtggtttgtttgccggatcaagagctacc
aactctttttccgaaggtaactggcttcagcagagcgcagataccaaatactgtccttctagtgtagccgtagttaggccaccacttcaagaactct
gtagcaccgcctacatacctcgctctgctaatcctgttaccagtggctgctgccagtggcgataagtcgtgtcttaccgggttggactcaagacgat
agttaccggataaggcgcagcggtcgggctgaacggggggttcgtgcacacagcccagcttggagcgaacgacctacaccgaactgagatacctaca
gcgtgagctatgagaaagcgccacgcttcccgaagggagaaaggcggacaggtatccggtaagcggcagggtcggaacaggagagcgcacgaggg
ag
cttccagggggaaacgcctggtatctttatagtcctgtcgggtttcgccacctctgacttgagcgtcgatttttgtgatgctcgtcaggggggcgga
gcctatggaaaaacgccagcaacgcggcctttttacggttcctggccttttgctggccttttgctcacatgttctttcctgcgttatcccctgattc
tgtggataaccgtattaccgcctttgagtgagctgataccgctcgccgcagccgaacgaccgagcgcagcgagtcagtgagcgaggaagcggaagag
cgcctgatgcggtattttctccttacgcatctgtgcggtatttcacaccgcaatggtgcactctcagtacaatctgctctgatgccgcatagttaag
ccagtatacactccgctatcgctacgtgactgggtcatggctgcgccccgacacccgccaacacccgctgacgcgccctgacgggcttgtctgctcc
cggcatccgcttacagacaagctgtgaccgtctccgggagctgcatgtgtcagaggttttcaccgtcatcaccgaaacgcgcgaggcagctgcggta
aagctcatcagcgtggtcgtgaagcgattcacagatgtctgcctgttcatccgcgtccagctcgttgagtttctccagaagcgttaatgtctggctt
ctgataaagcgggccatgttaagggcggttttttcctgtttggtcactgatgcctccgtgtaagggggatttctgttcatgggggtaatgataccga
tgaaacgagagaggatgctcacgatacgggttactgatgatgaacatgcccggttactggaacgttgtgagggtaaacaactggcggtatggatgcg
gcgggaccagagaaaaatcactcagggtcaatgccagcgcttcgttaatacagatgtaggtgttccacagggtagccagcagcatcctgcgatgcag
atccggaacataatggtgcagggcgctgacttccgcgtttccagactttacgaaacacggaaaccgaagaccattcatgttgttgctcaggtcgcag
acgttttgcagcagcagtcgcttcacgttcgctcgcgtatcggtgattcattctgctaaccagtaaggcaaccccgccagcctagccgggtcctcaa
cgacaggagcacgatcatgcgcacccgtggggccgccatgccggcgataatggcctgcttctcgccgaaacgtttggtggcgggaccagtgacgaag
gcttgagcgagggcgtgcaagattccgaataccgcaagcgacaggccgatcatcgtcgcgctccagcgaaagcggtcctcgccgaaaatgacccaga
gcgctgccggcacctgtcctacgagttgcatgataaagaagacagtcataagtgcggcgacgatagtcatgccccgcgcccaccggaaggagctgac
tgggttgaaggctctcaagggcatcggtcgagatcccggtgcctaatgagtgagctaacttacattaattgcgttgcgctcactgcccgctttccag
```

tcgggaaacctgtcgtgccagctgcattaatgaatcggccaacgcgcggggagaggcggtttgcgtattgggcgccagggtggtttttcttttcacc
agtgagacgggcaacagctgattgcccttcaccgcctggccctgagagagttgcagcaagcggtccacgctggtttgccccagcaggcgaaaatcct
gtttgatggtggttaacggcgggatataacatgagctgtcttcggtatcgtcgtatcccactaccgagatatccgcaccaacgcgcagcccggactc
ggtaatggcgcgcattgcgcccagcgccatctgatcgttggcaaccagcatcgcagtgggaacgatgccctcattcagcatttgcatggtttgttga
aaaccggacatggcactccagtcgccttcccgttccgctatcggctgaatttgattgcgagtgagatatttatgccagccagccagacgcagacgcg
ccgagacagaacttaatgggcccgctaacagcgcgatttgctggtgacccaatgcgaccagatgctccacgcccagtcgcgtaccgtcttcatggga
gaaaataatactgttgatgggtgtctggtcagagacatcaagaaataacgccggaacattagtgcaggcagcttccacagcaatggcatcctggtca
tccagcggatagttaatgatcagcccactgacgcgttgcgcgagaagattgtgcaccgccgctttacaggcttcgacgccgcttcgttctaccatcg
acaccaccacgctggcacccagttgatcggcgcgagatttaatcgccgcgacaatttgcgacggcgcgtgcagggccagactggaggtggcaacgcc
aatcagcaacgactgtttgcccgccagttgttgtgccacgcggttgggaatgtaattcagctccgccatcgccgcttccactttttcccgcgttttc
gcagaaacgtggctggcctggttcaccacgcgggaaacggtctgataagagacaccggcatactctgcgacatcgtataacgttactggtttcacat
tcaccaccctgaattgactctcttccgggcgctatcatgccataccgcgaaaggttttgcgccattcgatggtgtccgggatctcgacgctctccct
tatgcgactcctgcattaggaagcagcccagtagtaggttgaggccgttgagcaccgccgccgcaaggaatggtgcatgcaaggagatggcgcccaa
cagtcccccggccacggggcctgccaccatacccacgccgaaacaagcgctcatgagcccgaagtggcgagcccgatcttccccatcggtgatgtcg
gcgatataggcgccagcaaccgcacctgtggcgccggtgatgccggccacgatgcgtccggcgtagaggatcgagatctcgatcccgcgaaat

### ***Sequence of pET21a-RRF-His***

#### $P_{T7}$ promoter – RRF-His6 - T7 terminator

taatacgactcactataggggaattgtgagcggataacaattcccctctagaaataattttgtttaactttaagaaggagatatacatATGATTAGC
GATATCAGAAAAGATGCTGAAGTACGCATGGACAAATGCGTAGAAGCGTTCAAAACCCAAATCAGCAAAATACGCACGGGTCGTGCTTCTCCCAGCC
TGCTGGATGGCATTGTCGTGGAATATTACGGCACGCCGACGCCGCTGCGTCAGCTGGCAAGCGTAACGGTAGAAGATTCCCGTACACTGAAAATCAA
CGTGTTTGATCGTTCAATGTCTCCGGCCGTTGAAAAAGCGATTATGGCGTCCGATCTTGGCCTGAACCCGAACTCTGCGGGTAGCGACATCCGTGTT
CCGCTGCCGCCGCTGACGGAAGAACGTCGTAAAGATCTGACCAAAATCGTTCGTGGTGAAGCAGAACAAGCGCGTGTTGCAGTACGTAACGTGCGTC
GTGACGCGAACGACAAAGTGAAAGCACTGTTGAAAGATAAAGAGATCAGCGAAGACGACGATCGCCGTTCTCAGGACGATGTACAGAAACTGACTGA
TGCTGCAATCAAGAAAATTGAAGCGGCGCTGGCAGACAAAGAAGCAGAACTGATGCAGTTCGGATCCAGATCTCATCACCATCACCATCACTAATGA
gatccggctgctaacaaagcccgaaaggaagctgagttggctgctgccaccgctgagcaataactagcataaccccttggggcctctaaacgggtct
tgaggggttttttgctgaaaggaggaactatatccggattggcgaatgggacgcgccctgtagcggcgcattaagcgcggcgggtgtggtggttacg
cgcagcgtgaccgctacacttgccagcgccctagcgcccgctcctttcgctttcttcccttcctttctcgccacgttcgccggctttccccgtcaag
ctctaaatcgggggctccctttagggttccgatttagtgctttacggcacctcgaccccaaaaaacttgattagggtgatggttcacgtagtgggcc
atcgccctgatagacggtttttcgccctttgacgttggagtccacgttctttaatagtggactcttgttccaaactggaacaacactcaaccctatc
tcggtctattcttttgatttataagggattttgccgatttcggcctattggttaaaaaatgagctgatttaacaaaaatttaacgcgaattttaaca
aaatattaacgcttacaatttaggtggcacttttcggggaaatgtgcgcggaacccctatttgtttatttttctaaatacattcaaatatgtatccg
ctcatgagacaataaccctgataaatgcttcaataatattgaaaaaggaagagtatgagtattcaacatttccgtgtcgcccttattcccttttttg
cggcattttgccttcctgtttttgctcacccagaaacgctggtgaaagtaaaagatgctgaagatcagttgggtgcacgagtgggttacatcgaact
ggatctcaacagcggtaagatccttgagagttttcgccccgaagaacgttttccaatgatgagcacttttaaagttctgctatgtggcgcggtatta
tcccgtattgacgccgggcaagagcaactcggtcgccgcatacactattctcagaatgacttggttgagtactcaccagtcacagaaaagcatctta
cggatggcatgacagtaagagaattatgcagtgctgccataaccatgagtgataacactgcggccaacttacttctgacaacgatcggaggaccgaa
ggagctaaccgcttttttgcacaacatgggggatcatgtaactcgccttgatcgttgggaaccggagctgaatgaagccataccaaacgacgagcgt
gacaccacgatgcctgcagcaatggcaacaacgttgcgcaaactattaactggcgaactacttactctagcttcccggcaacaattaatagactgga
tggaggcggataaagttgcaggaccacttctgcgctcggcccttccggctggctggtttattgctgataaatctggagccggtgagcgtgggtctcg
cggtatcattgcagcactggggccagatggtaagccctcccgtatcgtagttatctacacgacggggagtcaggcaactatggatgaacgaaataga
cagatcgctgagataggtgcctcactgattaagcattggtaactgtcagaccaagtttactcatatatactttagattgatttaaaacttcatttt
aatttaaaaggatctaggtgaagatcctttttgataatctcatgaccaaaatcccttaacgtgagttttcgttccactgagcgtcagaccccgtaga
aaagatcaaaggatcttcttgagatcctttttttctgcgcgtaatctgctgcttgcaaacaaaaaaaccaccgctaccagcggtggtttgtttgccg
gatcaagagctaccaactctttttccgaaggtaactggcttcagcagagcgcagataccaaatactgtccttctagtgtagccgtagttaggccacc
acttcaagaactctgtagcaccgcctacatacctcgctctgctaatcctgttaccagtggctgctgccagtggcgataagtcgtgtcttaccgggtt
ggactcaagacgatagttaccggataaggcgcagcggtcgggctgaacggggggttcgtgcacacagcccagcttggagcgaacgacctacaccgaa
ctgagatacctacagcgtgagctatgagaaagcgccacgcttcccgaagggagaaaggcggacaggtatccggtaagcggcagggtcggaacaggag
agcgcacgagggagcttccagggggaaacgcctggtatctttatagtcctgtcgggtttcgccacctctgacttgagcgtcgatttttgtgatgctc
gtcaggggggcggagcctatggaaaaacgccagcaacgcggcctttttacggttcctggccttttgctggccttttgctcacatgttctttcctgcg
ttatcccctgattctgtggataaccgtattaccgcctttgagtgagctgataccgctcgccgcagccgaacgaccgagcgcagcgagtcagtgagcg
aggaagcggaagagcgcctgatgcggtattttctccttacgcatctgtgcggtatttcacaccgcaatggtgcactctcagtacaatctgctctgat
gccgcatagttaagccagtatacactccgctatcgctacgtgactgggtcatggctgcgccccgacacccgccaacacccgctgacgcgccctgacg
ggcttgtctgctcccggcatccgcttacagacaagctgtgaccgtctccgggagctgcatgtgtcagaggttttcaccgtcatcaccgaaacgcgcg
aggcagctgcggtaaagctcatcagcgtggtcgtgaagcgattcacagatgtctgcctgttcatccgcgtccagctcgttgagtttctccagaagcg
ttaatgtctggcttctgataaagcgggccatgttaagggcggttttttcctgtttggtcactgatgcctccgtgtaagggggatttctgttcatggg
ggtaatgataccgatgaaacgagagaggatgctcacgatacgggttactgatgatgaacatgcccggttactggaacgttgtgagggtaaacaactg
gcggtatggatgcggcgggaccagagaaaaatcactcagggtcaatgccagcgcttcgttaatacagatgtaggtgttccacagggtagccagcagc
atcctgcgatgcagatccggaacataatggtgcagggcgctgacttccgcgtttccagactttacgaaacacggaaaccgaagaccattcatgttgt
tgctcaggtcgcagacgttttgcagcagcagtcgcttcacgttcgctcgcgtatcggtgattcattctgctaaccagtaaggcaaccccgccagcct
agccgggtcctcaacgacaggagcacgatcatgcgcacccgtggggccgccatgccggcgataatggcctgcttctcgccgaaacgtttggtggcgg
gaccagtgacgaaggcttgagcgagggcgtgcaagattccgaataccgcaagcgacaggccgatcatcgtcgcgctccagcgaaagcggtcctcgcc
gaaaatgacccagagcgctgccggcacctgtcctacgagttgcatgataaagaagacagtcataagtgcggcgacgatagtcatgccccgcgcccac
cggaaggagctgactgggttgaaggctctcaagggcatcggtcgagatcccggtgcctaatgagtgagctaacttacattaattgcgttgcgctcac
tgcccgctttccagtcgggaaacctgtcgtgccagctgcattaatgaatcggccaacgcgcggggagaggcggtttgcgtattgggcgccagggtgg
tttttcttttcaccagtgagacgggcaacagctgattgcccttcaccgcctggccctgagagagttgcagcaagcggtccacgctggtttgccccag
caggcgaaaatcctgtttgatggtggttaacggcgggatataacatgagctgtcttcggtatcgtcgtatcccactaccgagatatccgcaccaacg
cgcagcccggactcggtaatggcgcgcattgcgcccagcgccatctgatcgttggcaaccagcatcgcagtgggaacgatgccctcattcagcattt
gcatggtttgttgaaaaccggacatggcactccagtcgccttcccgttccgctatcggctgaatttgattgcgagtgagatatttatgccagccagc
cagacgcagacgcgccgagacagaacttaatgggcccgctaacagcgcgatttgctggtgacccaatgcgaccagatgctccacgcccagtcgcgta
ccgtcttcatgggagaaaataatactgttgatgggtgtctggtcagagacatcaagaaataacgccggaacattagtgcaggcagcttccacagcaa
tggcatcctggtcatccagcggatagttaatgatcagcccactgacgcgttgcgcgagaagattgtgcaccgccgctttacaggcttcgacgccgct
tcgttctaccatcgacaccaccacgctggcacccagttgatcggcgcgagatttaatcgccgcgacaatttgcgacggcgcgtgcagggccagactg
gaggtggcaacgccaatcagcaacgactgtttgcccgccagttgttgtgccacgcggttgggaatgtaattcagctccgccatcgccgcttccactt
tttcccgcgttttcgcagaaacgtggctggcctggttcaccacgcgggaaacggtctgataagagacaccggcatactctgcgacatcgtataacgt

tactggtttcacattcaccaccctgaattgactctcttccgggcgctatcatgccataccgcgaaaggttttgcgccattcgatggtgtccgggatc
tcgacgctctcccttatgcgactcctgcattaggaagcagcccagtagtaggttgaggccgttgagcaccgccgccgcaaggaatggtgcatgcaag
gagatggcgcccaacagtcccccggccacggggcctgccaccatacccacgccgaaacaagcgctcatgagcccgaagtggcgagcccgatcttccc
catcggtgatgtcggcgatataggcgccagcaaccgcacctgtggcgccggtgatgccggccacgatgcgtccggcgtagaggatcgagatctcgat
cccgcgaaat

### ***Sequence of pET21a-His-CKM***

#### $P_{T7}$ promoter – His6-CKM - T7 terminator

taatacgactcactataggggaattgtgagcggataacaattccсctctagaaataattttgtttaactttaagaaggagatatacatATGAGAGGA
TCGCATCACCATCACCATCACGGATCCATGCCGTTCAGCAGCACCCACAACAAACACAAGCTGAAGTTCTCAGCCGAGGAGGAATTCCCGGACCTCT
CGAAGCACAACAACCACATGGCCAAAGTCCTCACCCCGGAGCTCTACAAACGCCTCCGGGATAAGGAGACCCCGAGCGGATTCACCCTCGACGACGT
CATCCAAACCGGGGTCGATAACCCCGGTCACCCATTCATCATGACGGTGGGCTGCGTGGCGGGGGACGAGGAGTCCTACGAGGTGTTCAAGGATCTC
TTCGACCCCGTCATCCAGGACCGCCACGGGGGGTACAAACCCACCGATAAGCACCGCACCGACCTCAACCACGAGAACCTCAAGGGGGGTGACGACC
TGGACCCCAAATACGTGCTGAGCAGCCGCGTGCGCACGGGGAGAAGCATTAAGGGGTACTCCCTGCCCCCACACTGCAGCCGTGGGGAGCGGCGCGC
CGTCGAGAAGCTGTCCGTGGAAGCCCTGAACAGCCTGGAGGGGGAGTTCAAGGGCCGCTATTACCCTCTGAAGGCCATGACGGAGCAGGAGCAGCAG
CAGCTGATCGACGACCACTTCCTGTTCGATAAACCCGTCTCCCCACTGCTGCTCGCATCCGGGATGGCCCGAGATTGGCCCGACGCCAGGGGCATCT
GGCACAACGACAACAAGACGTTCCTGGTGTGGGTGAATGAGGAGGACCACCTGAGGGTCATCTCCATGGAGAAGGGAGGAAATATGAAGGAGGTCTT
CCGGCGCTTCTGCGTCGGCCTCAAGAAGATCGAGGAGATCTTCAAGAAGGCCGGGCACCCCTTCATGTGGACGGAGCACCTGGGTTACATCCTGACG
TGCCCCTCCAATTTGGGCACGGGGCTGCGGGGGGGGGTCCACGTGAAGCTCCCCAAACTCAGCCAGCACCCCAAATTCGAGGAGATCCTCCATAGGC
TGCGCCTGCAGAAACGGGGCACCGGCGGGGTGGATACGGCGGCGGTGGGCGCCGTCTTTGACATCTCCAACGCCGACCGGCTGGGCTTCTCGGAGGT
GGAGCAGGTGCAGATGGTGGTGGACGGCGTCAAGCTCATGGTGGAGATGGAGAAGAAGCTGGAGCAGAACCAGCCCATAGACGACATGATCCCGGCC
CAGAAGTAATGAgatccggctgctaacaaagcccgaaaggaagctgagttggctgctgccaccgctgagcaataactagcataaccccttggggcct
ctaaacgggtcttgaggggttttttgctgaaaggaggaactatatccggattggcgaatgggacgcgccctgtagcggcgcattaagcgcggcgggt
gtggtggttacgcgcagcgtgaccgctacacttgccagcgccctagcgcccgctcctttcgctttcttcccttcctttctcgccacgttcgccggct
ttccccgtcaagctctaaatcgggggctccctttagggttccgatttagtgctttacggcacctcgaccccaaaaaacttgattagggtgatggttc
acgtagtgggccatcgccctgatagacggtttttcgccctttgacgttggagtccacgttctttaatagtggactcttgttccaaactggaacaaca
ctcaaccctatctcggtctattcttttgatttataagggattttgccgatttcggcctattggttaaaaaatgagctgatttaacaaaaatttaacg
cgaattttaacaaaatattaacgcttacaatttaggtggcacttttcggggaaatgtgcgcggaacccctatttgtttatttttctaaatacattca
aatatgtatccgctcatgagacaataaccctgataaatgcttcaataatattgaaaaaggaagagtatgagtattcaacatttccgtgtcgccctta
ttcccttttttgcggcattttgccttcctgtttttgctcacccagaaacgctggtgaaagtaaaagatgctgaagatcagttgggtgcacgagtggg
ttacatcgaactggatctcaacagcggtaagatccttgagagttttcgccccgaagaacgttttccaatgatgagcacttttaaagttctgctatgt
ggcgcggtattatcccgtattgacgccgggcaagagcaactcggtcgccgcatacactattctcagaatgacttggttgagtactcaccagtcacag
aaaagcatcttacggatggcatgacagtaagagaattatgcagtgctgccataaccatgagtgataacactgcggccaacttacttctgacaacgat
cggaggaccgaaggagctaaccgcttttttgcacaacatgggggatcatgtaactcgccttgatcgttgggaaccggagctgaatgaagccatacca
aacgacgagcgtgacaccacgatgcctgcagcaatggcaacaacgttgcgcaaactattaactggcgaactacttactctagcttcccggcaacaat
taatagactggatggaggcggataaagttgcaggaccacttctgcgctcggcccttccggctggctggtttattgctgataaatctggagccggtga
gcgtgggtctcgcggtatcattgcagcactggggccagatggtaagccctcccgtatcgtagttatctacacgacggggagtcaggcaactatggat
gaacgaaatagacagatcgctgagataggtgcctcactgattaagcattggtaactgtcagaccaagtttactcatatatactttagattgatttaa
aacttcatttttaatttaaaaggatctaggtgaagatcctttttgataatctcatgaccaaaatcccttaacgtgagttttcgttccactgagcgtc
agaccccgtagaaaagatcaaaggatcttcttgagatcctttttttctgcgcgtaatctgctgcttgcaaacaaaaaaaccaccgctaccagcggtg
gtttgtttgccggatcaagagctaccaactctttttccgaaggtaactggcttcagcagagcgcagataccaaatactgtccttctagtgtagccgt
agttaggccaccacttcaagaactctgtagcaccgcctacatacctcgctctgctaatcctgttaccagtggctgctgccagtggcgataagtcgtg
tcttaccgggttggactcaagacgatagttaccggataaggcgcagcggtcgggctgaacggggggttcgtgcacacagcccagcttggagcgaacg
acctacaccgaactgagatacctacagcgtgagctatgagaaagcgccacgcttcccgaagggagaaaggcggacaggtatccggtaagcggcaggg
tcggaacaggagagcgcacgagggagcttccagggggaaacgcctggtatctttatagtcctgtcgggtttcgccacctctgacttgagcgtcgatt
tttgtgatgctcgtcaggggggcggagcctatggaaaaacgccagcaacgcggcctttttacggttcctggccttttgctggccttttgctcacatg
ttctttcctgcgttatcccctgattctgtggataaccgtattaccgcctttgagtgagctgataccgctcgccgcagccgaacgaccgagcgcagcg
agtcagtgagcgaggaagcggaagagcgcctgatgcggtattttctccttacgcatctgtgcggtatttcacaccgcaatggtgcactctcagtaca
atctgctctgatgccgcatagttaagccagtatacactccgctatcgctacgtgactgggtcatggctgcgccccgacacccgccaacacccgctga
cgcgccctgacgggcttgtctgctcccggcatccgcttacagacaagctgtgaccgtctccgggagctgcatgtgtcagaggttttcaccgtcatca
ccgaaacgcgcgaggcagctgcggtaaagctcatcagcgtggtcgtgaagcgattcacagatgtctgcctgttcatccgcgtccagctcgttgagtt
tctccagaagcgttaatgtctggcttctgataaagcgggccatgttaagggcggtttttcctgtttggtcactgatgcctccgtgtaagggggatt
tctgttcatgggggtaatgataccgatgaaacgagagaggatgctcacgatacgggttactgatgatgaacatgcccggttactggaacgttgtgag
ggtaaacaactggcggtatggatgcggcgggaccagagaaaaatcactcagggtcaatgccagcgcttcgttaatacagatgtaggtgttccacagg
gtagccagcagcatcctgcgatgcagatccggaacataatggtgcagggcgctgacttccgcgtttccagactttacgaaacacggaaaccgaagac
cattcatgttgttgctcaggtcgcagacgttttgcagcagcagtcgcttcacgttcgctcgcgtatcggtgattcattctgctaaccagtaaggcaa
ccccgccagcctagccgggtcctcaacgacaggagcacgatcatgcgcacccgtggggccgccatgccggcgataatggcctgcttctcgccgaaac
gtttggtggcgggaccagtgacgaaggcttgagcgagggcgtgcaagattccgaataccgcaagcgacaggccgatcatcgtcgcgctccagcgaaa
gcggtcctcgccgaaaatgacccagagcgctgccggcacctgtcctacgagttgcatgataaagaagacagtcataagtgcggcgacgatagtcatg
ccccgcgcccaccggaaggagctgactgggttgaaggctctcaagggcatcggtcgagatcccggtgcctaatgagtgagctaacttacattaattg
cgttgcgctcactgcccgctttccagtcgggaaacctgtcgtgccagctgcattaatgaatcggccaacgcgcggggagaggcggtttgcgtattgg
gcgccagggtggtttttcttttcaccagtgagacgggcaacagctgattgcccttcaccgcctggccctgagagagttgcagcaagcggtccacgct
ggtttgccccagcaggcgaaaatcctgtttgatggtggttaacggcgggatataacatgagctgtcttcggtatcgtcgtatcccactaccgagata
tccgcaccaacgcgcagcccggactcggtaatggcgcgcattgcgcccagcgccatctgatcgttggcaaccagcatcgcagtgggaacgatgccct
cattcagcatttgcatggtttgttgaaaaccggacatggcactccagtcgccttcccgttccgctatcggctgaatttgattgcgagtgagatattt

atgccagccagccagacgcagacgcgccgagacagaacttaatgggcccgctaacagcgcgatttgctggtgacccaatgcgaccagatgctccacg
cccagtcgcgtaccgtcttcatgggagaaaataatactgttgatgggtgtctggtcagagacatcaagaaataacgccggaacattagtgcaggcag
cttccacagcaatggcatcctggtcatccagcggatagttaatgatcagcccactgacgcgttgcgcgagaagattgtgcaccgccgctttacaggc
ttcgacgccgcttcgttctaccatcgacaccaccacgctggcacccagttgatcggcgcgagatttaatcgccgcgacaatttgcgacggcgcgtgc
agggccagactggaggtggcaacgccaatcagcaacgactgtttgcccgccagttgttgtgccacgcggttgggaatgtaattcagctccgccatcg
ccgcttccactttttcccgcgttttcgcagaaacgtggctggcctggttcaccacgcgggaaacggtctgataagagacaccggcatactctgcgac
atcgtataacgttactggtttcacattcaccaccctgaattgactctcttccgggcgctatcatgccataccgcgaaaggttttgcgccattcgatg
gtgtccgggatctcgacgctctcccttatgcgactcctgcattaggaagcagcccagtagtaggttgaggccgttgagcaccgccgccgcaaggaat
ggtgcatgcaaggagatggcgcccaacagtcccccggccacggggcctgccaccatacccacgccgaaacaagcgctcatgagcccgaagtggcgag
cccgatcttccccatcggtgatgtcggcgatataggcgccagcaaccgcacctgtggcgccggtgatgccggccacgatgcgtccggcgtagaggat
cgagatctcgatcccgcgaaat

### *Sequence of pET21a-His-NDK*

#### $P_{T7}$ promoter – His6-NDK - T7 terminator

taatacgactcactataggggaattgtgagcggataacaattcccctctagaaataattttgtttaactttaagaaggagatatacatATGAGAGGA
TCGCATCACCATCACCATCACGGATCCGATGACGATGACAAAGCTATTGAACGTACTTTTTCCATCATCAAACCGAACGCGGTAGCAAAAAACGTCA
TTGGTAATATCTTTGCGCGCTTTGAAGCTGCAGGGTTCAAAATTGTTGGCACCAAAATGCTGCACCTGACCGTTGAACAGGCACGTGGCTTTTATGC
TGAACACGATGGAAAACCGTTCTTTGATGGTCTGGTTGAATTCATGACCTCTGGCCCGATCGTGGTTTCCGTGCTGGAAGGTGAAAACGCCGTTCAG
CGTCACCGCGATCTGCTGGGCGCGACCAATCCGGCAAACGCACTGGCTGGTACTCTGCGCGCTGATTACGCTGACAGCCTGACCGAAAACGGTACCC
ACGGTTCTGATTCCGTCGAATCTGCCGCTCGCGAAATCGCTTATTTCTTTGGCGAAGGCGAAGTGTGCCCGCGCACCCGTTAATGAgatccggctgc
taacaaagcccgaaaggaagctgagttggctgctgccaccgctgagcaataactagcataaccccttggggcctctaaacgggtcttgaggggtttt
ttgctgaaaggaggaactatatccggattggcgaatgggacgcgccctgtagcggcgcattaagcgcggcgggtgtggtggttacgcgcagcgtgac
cgctacacttgccagcgccctagcgcccgctcctttcgctttcttcccttcctttctcgccacgttcgccggctttccccgtcaagctctaaatcgg
gggctccctttagggttccgatttagtgctttacggcacctcgaccccaaaaaacttgattagggtgatggttcacgtagtgggccatcgccctgat
agacggtttttcgccctttgacgttggagtccacgttctttaatagtggactcttgttccaaactggaacaacactcaaccctatctcggtctattc
ttttgatttataagggattttgccgatttcggcctattggttaaaaaatgagctgatttaacaaaaatttaacgcgaattttaacaaaatattaacg
cttacaatttaggtggcacttttcggggaaatgtgcgcggaacccctatttgtttatttttctaaatacattcaaatatgtatccgctcatgagaca
ataaccctgataaatgcttcaataatattgaaaaaggaagagtatgagtattcaacatttccgtgtcgcccttattcccttttttgcggcattttgc
cttcctgtttttgctcacccagaaacgctggtgaaagtaaaagatgctgaagatcagttgggtgcacgagtgggttacatcgaactggatctcaaca
gcggtaagatccttgagagttttcgccccgaagaacgttttccaatgatgagcacttttaaagttctgctatgtggcgcggtattatcccgtattga
cgccgggcaagagcaactcggtcgccgcatacactattctcagaatgacttggttgagtactcaccagtcacagaaaagcatcttacggatggcatg
acagtaagagaattatgcagtgctgccataaccatgagtgataacactgcggccaacttacttctgacaacgatcggaggaccgaaggagctaaccg
cttttttgcacaacatgggggatcatgtaactcgccttgatcgttgggaaccggagctgaatgaagccataccaaacgacgagcgtgacaccacgat
gcctgcagcaatggcaacaacgttgcgcaaactattaactggcgaactacttactctagcttcccggcaacaattaatagactggatggaggcggat
aaagttgcaggaccacttctgcgctcggcccttccggctggctggtttattgctgataaatctggagccggtgagcgtgggtctcgcggtatcattg
cagcactggggccagatggtaagccctcccgtatcgtagttatctacacgacggggagtcaggcaactatggatgaacgaaatagacagatcgctga
gataggtgcctcactgattaagcattggtaactgtcagaccaagtttactcatatatactttagattgatttaaaacttcatttttaatttaaaagg
atctaggtgaagatcctttttgataatctcatgaccaaaatcccttaacgtgagttttcgttccactgagcgtcagaccccgtagaaaagatcaaag
gatcttcttgagatcctttttttctgcgcgtaatctgctgcttgcaaacaaaaaaaccaccgctaccagcggtggtttgtttgccggatcaagagct
accaactctttttccgaaggtaactggcttcagcagagcgcagataccaaatactgtccttctagtgtagccgtagttaggccaccacttcaagaac
tctgtagcaccgcctacatacctcgctctgctaatcctgttaccagtggctgctgccagtggcgataagtcgtgtcttaccgggttggactcaagac
gatagttaccggataaggcgcagcggtcgggctgaacggggggttcgtgcacacagcccagcttggagcgaacgacctacaccgaactgagatacct
acagcgtgagctatgagaaagcgccacgcttcccgaagggagaaaggcggacaggtatccggtaagcggcagggtcggaacaggagagcgcacgagg
gagcttccagggggaaacgcctggtatctttatagtcctgtcgggtttcgccacctctgacttgagcgtcgatttttgtgatgctcgtcaggggggc
ggagcctatggaaaaacgccagcaacgcggcctttttacggttcctggccttttgctggccttttgctcacatgttctttcctgcgttatcccctga
ttctgtggataaccgtattaccgcctttgagtgagctgataccgctcgccgcagccgaacgaccgagcgcagcgagtcagtgagcgaggaagcggaa
gagcgcctgatgcggtattttctccttacgcatctgtgcggtatttcacaccgcaatggtgcactctcagtacaatctgctctgatgccgcatagtt
aagccagtatacactccgctatcgctacgtgactgggtcatggctgcgccccgacacccgccaacacccgctgacgcgccctgacgggcttgtctgc
tcccggcatccgcttacagacaagctgtgaccgtctccgggagctgcatgtgtcagaggttttcaccgtcatcaccgaaacgcgcgaggcagctgcg
gtaaagctcatcagcgtggtcgtgaagcgattcacagatgtctgcctgttcatccgcgtccagctcgttgagtttctccagaagcgttaatgtctgg
cttctgataaagcgggccatgttaagggcggttttttcctgtttggtcactgatgcctccgtgtaagggggatttctgttcatgggggtaatgatac
cgatgaaacgagagaggatgctcacgatacgggttactgatgatgaacatgcccggttactggaacgttgtgagggtaaacaactggcggtatggat
gcggcgggaccagagaaaaatcactcagggtcaatgccagcgcttcgttaatacagatgtaggtgttccacagggtagccagcagcatcctgcgatg
cagatccggaacataatggtgcagggcgctgacttccgcgtttccagactttacgaaacacggaaaccgaagaccattcatgttgttgctcaggtcg
cagacgttttgcagcagcagtcgcttcacgttcgctcgcgtatcggtgattcattctgctaaccagtaaggcaaccccgccagcctagccgggtcct
caacgacaggagcacgatcatgcgcacccgtggggccgccatgccggcgataatggcctgcttctcgccgaaacgtttggtggcgggaccagtgacg
aaggcttgagcgagggcgtgcaagattccgaataccgcaagcgacaggccgatcatcgtcgcgctccagcgaaagcggtcctcgccgaaaatgaccc
agagcgctgccggcacctgtcctacgagttgcatgataaagaagacagtcataagtgcggcgacgatagtcatgccccgcgcccaccggaaggagct
gactgggttgaaggctctcaagggcatcggtcgagatcccggtgcctaatgagtgagctaacttacattaattgcgttgcgctcactgcccgctttc
cagtcgggaaacctgtcgtgccagctgcattaatgaatcggccaacgcgcggggagaggcggtttgcgtattgggcgccagggtggtttttcttttc
accagtgagacgggcaacagctgattgcccttcaccgcctggccctgagagagttgcagcaagcggtccacgctggtttgccccagcaggcgaaaat
cctgtttgatggtggttaacggcgggatataacatgagctgtcttcggtatcgtcgtatcccactaccgagatatccgcaccaacgcgcagcccgga
ctcggtaatggcgcgcattgcgcccagcgccatctgatcgttggcaaccagcatcgcagtgggaacgatgccctcattcagcatttgcatggtttgt
tgaaaaccggacatggcactccagtcgccttcccgttccgctatcggctgaatttgattgcgagtgagatatttatgccagccagccagacgcagac
gcgccgagacagaacttaatgggcccgctaacagcgcgatttgctggtgacccaatgcgaccagatgctccacgcccagtcgcgtaccgtcttcatg
ggagaaaataatactgttgatgggtgtctggtcagagacatcaagaaataacgccggaacattagtgcaggcagcttccacagcaatggcatcctgg
tcatccagcggatagttaatgatcagcccactgacgcgttgcgcgagaagattgtgcaccgccgctttacaggcttcgacgccgcttcgttctacca
tcgacaccaccacgctggcacccagttgatcggcgcgagatttaatcgccgcgacaatttgcgacggcgcgtgcagggccagactggaggtggcaac
gccaatcagcaacgactgtttgcccgccagttgttgtgccacgcggttgggaatgtaattcagctccgccatcgccgcttccactttttcccgcgtt
ttcgcagaaacgtggctggcctggttcaccacgcgggaaacggtctgataagagacaccggcatactctgcgacatcgtataacgttactggtttca
cattcaccaccctgaattgactctcttccgggcgctatcatgccataccgcgaaaggttttgcgccattcgatggtgtccgggatctcgacgctctc

ccttatgcgactcctgcattaggaagcagcccagtagtaggttgaggccgttgagcaccgccgccgcaaggaatggtgcatgcaaggagatggcgcc
caacagtcccccggccacggggcctgccaccatacccacgccgaaacaagcgctcatgagcccgaagtggcgagcccgatcttccccatcggtgatg
tcggcgatataggcgccagcaaccgcacctgtggcgccggtgatgccggccacgatgcgtccggcgtagaggatcgagatctcgatcccgcgaaat

#### ***Sequence of pP$_{T7}$-MGA-UTR1-deGFP-T$_{T7}$***

P$_{T7}$ promoter-MGA-UTR1-deGFP-T7 terminator

taatacgactcactatagggagaccacaacggtttccctctagagggatcccgactggcgagagccaggtaacgaatggatccaataattttgttta
actttaagaaggagatataccATGGAGCTTTTCACTGGCGTTGTTCCCATCCTGGTCGAGCTGGACGGCGACGTAAACGGCCACAAGTTCAGCGTGT
CCGGCGAGGGCGAGGGCGATGCCACCTACGGCAAGCTGACCCTGAAGTTCATCTGCACCACCGGCAAGCTGCCCGTGCCCTGGCCCACCCTCGTGAC
CACCCTGACCTACGGCGTGCAGTGCTTCAGCCGCTACCCCGACCACATGAAGCAGCACGACTTCTTCAAGTCCGCCATGCCCGAAGGCTACGTCCAG
GAGCGCACCATCTTCTTCAAGGACGACGGCAACTACAAGACCCGCGCCGAGGTGAAGTTCGAGGGCGACACCCTGGTGAACCGCATCGAGCTGAAGG
GCATCGACTTCAAGGAGGACGGCAACATCCTGGGGCACAAGCTGGAGTACAACTACAACAGCCACAACGTCTATATCATGGCCGACAAGCAGAAGAA
CGGCATCAAGGTGAACTTCAAGATCCGCCACAACATCGAGGACGGCAGCGTGCAGCTCGCCGACCACTACCAGCAGAACACCCCCATCGGCGACGGC
CCCGTGCTGCTGCCCGACAACCACTACCTGAGCACCCAGTCCGCCCTGAGCAAAGACCCCAACGAGAAGCGCGATCACATGGTCCTGCTGGAGTTCG
TGACCGCCGCCGGGATCTAActcgagccttaggagatccggctgctaacaaagcccgaaaggaagctgagttggctgctgccaccgctgagcaataa
ctagcataaccccttggggcctctaaacgggtcttgaggggttttttgctgaaaggaggaactatatccggatatccacaggacgggtgtggtcgcc
atgatcgcgtagtcgatagtggctccaagtagcgaagcgagcaggactgggcggcggccaaagcggtcggacagtgctccgagaacgggtgcgcata
gaaattgcatcaacgcatatagcgctagcagcacgccatagtgactggcgatgctgtcggaatggacgatatcccgcaagaggcccggcagtaccgg
cataaccaagcctatgcctacagcatccagggtgacggtgccgaggatgacgatgagcgcattgttagatttcatacacggtgcctgactgcgttag
caatttaactgtgataaactaccgcattaaagcttatcgatgataagctgtcaaacatgagaattcgtaatcatgtcatagctgtttcctgtgtgaa
attgttatccgctcacaattccacacaacatacgagccggaagcataaagtgtaaagcctggggtgcctaatgagtgagctaactcacattaattgc
gttgcgctcactgcccgctttccagtcgggaaacctgtcgtgccagctgcattaatgaatcggccaacgcgcggggagaggcggtttgcgtattggg
cgctcttccgcttcctcgctcactgactcgctgcgctcggtcgttcggctgcggcgagcggtatcagctcactcaaaggcggtaatacggttatcca
cagaatcaggggataacgcaggaaagaacatgtgagcaaaaggccagcaaaaggccaggaaccgtaaaaaggccgcgttgctggcgtttttccatag
gctccgcccccctgacgagcatcacaaaaatcgacgctcaagtcagaggtggcgaaacccgacaggactataaagataccaggcgtttccccctgga
agctccctcgtgcgctctcctgttccgaccctgccgcttaccggatacctgtccgcctttctcccttcgggaagcgtggcgctttctcatagctcac
gctgtaggtatctcagttcggtgtaggtcgttcgctccaagctgggctgtgtgcacgaaccccccgttcagcccgaccgctgcgccttatccggtaa
ctatcgtcttgagtccaacccggtaagacacgacttatcgccactggcagcagccactggtaacaggattagcagagcgaggtatgtaggcggtgct
acagagttcttgaagtggtggcctaactacggctacactagaagaacagtatttggtatctgcgctctgctgaagccagttaccttcggaaaaagag
ttggtagctcttgatccggcaaacaaaccaccgctggtagcggtggtttttttgtttgcaagcagcagattacgcgcagaaaaaaaggatctcaaga
agatcctttgatcttttctacggggtctgacgctcagtggaacgaaaactcacgttaagggattttggtcatgagattatcaaaaaggatcttcacc
tagatccttttaaattaaaaatgaagttttaaatcaatctaaagtatatatgagtaaacttggtctgacagttaccaatgcttaatcagtgaggcac
ctatctcagcgatctgtctatttcgttcatccatagttgcctgactccccgtcgtgtagataactacgatacgggagggcttaccatctggccccag
tgctgcaatgataccgcgagacccacgctcaccggctccagatttatcagcaataaaccagccagccggaagggccgagcgcagaagtggtcctgca
actttatccgcctccatccagtctattaattgttgccgggaagctagagtaagtagttcgccagttaatagtttgcgcaacgttgttgccattgcta
caggcatcgtggtgtcacgctcgtcgtttggtatggcttcattcagctccggttcccaacgatcaaggcgagttacatgatcccccatgttgtgcaa
aaaagcggttagctccttcggtcctccgatcgttgtcagaagtaagttggccgcagtgttatcactcatggttatggcagcactgcataattctctt
actgtcatgccatccgtaagatgcttttctgtgactggtgagtactcaaccaagtcattctgagaatagtgtatgcggcgaccgagttgctcttgcc
cggcgtcaatacgggataataccgcgccacatagcagaactttaaaagtgctcatcattggaaaacgttcttcggggcgaaaactctcaaggatctt
accgctgttgagatccagttcgatgtaacccactcgtgcacccaactgatcttcagcatcttttactttcaccagcgtttctgggtgagcaaaaaca
ggaaggcaaaatgccgcaaaaaagggaataagggcgacacggaaatgttgaatactcatactcttcctttttcaatattattgaagcatttatcagg
gttattgtctcatgagcggatacatatttgaatgtatttagaaaaataaacaaataggggttccgcgcacatttccccgaaaagtgccacctgacgt
ctaagaaaccattattatcatgacattaacctataaaaataggcgtatcacgaggccctttcgtctcgcgcgtttcggtgatgacggtgaaaacctc
tgacacatgcagctcccggagacggtcacagcttgtctgtaagcggatgccgggagcagacaagcccgtcagggcgcgtcagcgggtgttggcgggt
gtcggggctggcttaactatgcggcatcagagcagattgtactgagagtgcaccatatatgcggtgtgaaataccgcacagatgcgtaaggagaaaa
taccgcatcaggcgccattcgccattcaggctgcgcaactgttgggaagggcgatcggtgcgggcctcttcgctattacgccagctggcgaaagggg
gatgtgctgcaaggcgattaagttgggtaacgccagggttttcccagtcacgacgttgtaaaacgacggccagtgccaagcttgcatgcaaggagat
ggcgcccaacagtcccccggccacggggcctgccaccatacccacgccgaaacaagcgctcatgagcccgaagtggcgagcccgatcttccccatcg
gtgatgtcggcgatataggcgccagcaaccgcacctgtggcgccggtgatgccggccacgatgcgtccggcgtagaggatcgagatctcgatcccgc
gaaat

#### ***Sequence of pP$_{T7}$-UTR1-deGFP-T$_{T7}$***

P$_{T7}$ promoter- UTR1-deGFP-T7 terminator

taatacgactcactatagggGctagccaataattttgtttaactttaagaaggagatataccATGGAGCTTTTCACTGGCGTTGTTCCCATCCTGGT
CGAGCTGGACGGCGACGTAAACGGCCACAAGTTCAGCGTGTCCGGCGAGGGCGAGGGCGATGCCACCTACGGCAAGCTGACCCTGAAGTTCATCTGC
ACCACCGGCAAGCTGCCCGTGCCCTGGCCCACCCTCGTGACCACCCTGACCTACGGCGTGCAGTGCTTCAGCCGCTACCCCGACCACATGAAGCAGC
ACGACTTCTTCAAGTCCGCCATGCCCGAAGGCTACGTCCAGGAGCGCACCATCTTCTTCAAGGACGACGGCAACTACAAGACCCGCGCCGAGGTGAA
GTTCGAGGGCGACACCCTGGTGAACCGCATCGAGCTGAAGGGCATCGACTTCAAGGAGGACGGCAACATCCTGGGGCACAAGCTGGAGTACAACTAC
AACAGCCACAACGTCTATATCATGGCCGACAAGCAGAAGAACGGCATCAAGGTGAACTTCAAGATCCGCCACAACATCGAGGACGGCAGCGTGCAGC
TCGCCGACCACTACCAGCAGAACACCCCCATCGGCGACGGCCCCGTGCTGCTGCCCGACAACCACTACCTGAGCACCCAGTCCGCCCTGAGCAAAGA
CCCCAACGAGAAGCGCGATCACATGGTCCTGCTGGAGTTCGTGACCGCCGCCGGGATCTAActcgagctagcataaccccttggggcctctaaacgg
gtcttgaggggttttttggtcgaccgatgcccttgagagccttcaacccagtcagctccttccggtgggcgcggggcatgactatcgtcgccgcact
tatgactgtcttctttatcatgcaactcgtaggacaggtgccggcagcgctcttccgcttcctcgctcactgactcgctgcgctcggtcgttcggct
gcggcgagcggtatcagctcactcaaaggcggtaatacggttatccacagaatcaggggataacgcaggaaagaacatgtgagcaaaaggccagcaa
aaggccaggaaccgtaaaaaggccgcgttgctggcgtttttccataggctccgcccccctgacgagcatcacaaaaatcgacgctcaagtcagaggt
ggcgaaacccgacaggactataaagataccaggcgtttccccctggaagctccctcgtgcgctctcctgttccgaccctgccgcttaccggatacct
gtccgcctttctcccttcgggaagcgtggcgctttctcaatgctcacgctgtaggtatctcagttcggtgtaggtcgttcgctccaagctgggctgt
gtgcacgaaccccccgttcagcccgaccgctgcgccttatccggtaactatcgtcttgagtccaacccggtaagacacgacttatcgccactggcag
cagccactggtaacaggattagcagagcgaggtatgtaggcggtgctacagagttcttgaagtggtggcctaactacggctacactagaaggacagt
atttggtatctgcgctctgctgaagccagttaccttcggaaaaagagttggtagctcttgatccggcaaacaaaccaccgctggtagcggtggtttt
tttgtttgcaagcagcagattacgcgcagaaaaaaaggatctcaagaagatcctttgatcttttctacggggtctgacgctcagtggaacgaaaact
cacgttaagggattttggtcatgagattatcaaaaaggatcttcacctagatccttttaaattaaaaatgaagttttaaatcaatctaaagtatata
tgagtaaacttggtctgacagttaccaatgcttaatcagtgaggcacctatctcagcgatctgtctatttcgttcatccatagttgcctgactcccc
gtcgtgtagataactacgatacgggagggcttaccatctggccccagtgctgcaatgataccgcgagacccacgctcaccggctccagatttatcag
caataaaccagccagccggaagggccgagcgcagaagtggtcctgcaactttatccgcctccatccagtctattaattgttgccgggaagctagagt
aagtagttcgccagttaatagtttgcgcaacgttgttgccattgctacaggcatcgtggtgtcacgctcgtcgtttggtatggcttcattcagctcc
ggttcccaacgatcaaggcgagttacatgatcccccatgttgtgcaaaaaagcggttagctccttcggtcctccgatcgttgtcagaagtaagttgg
ccgcagtgttatcactcatggttatggcagcactgcataattctcttactgtcatgccatccgtaagatgcttttctgtgactggtgagtactcaac
caagtcattctgagaatagtgtatgcggcgaccgagttgctcttgcccggcgtcaatacgggataataccgcgccacatagcagaactttaaaagtg
ctcatcattggaaaacgttcttcggggcgaaaactctcaaggatcttaccgctgttgagatccagttcgatgtaacccactcgtgcacccaactgat
cttcagcatcttttactttcaccagcgtttctgggtgagcaaaaacaggaaggcaaaatgccgcaaaaaagggaataagggcgacacggaaatgttg
aatactcatactcttcctttttcaatattattgaagcatttatcagggttattgtctcatgagcggatacatatttgaatgtatttagaaaaataaa
caaataggggttccgcgcacatttccccgaaaagtgccacctgacgtctaagaaaccattattatcatgacattaacctataaaaataggcgtatca
cgaggccctttcgtcttcaagaattctggcgaatcctctgaccagccagaaaacgacctttctgtggtgaaaccggatgctgcaattcagagcggca
gcaagtgggggacagcagaagacctgaccgccgcagagtggatgtttgacatggtgaagactatcgcaccatcagccagaaaaccgaattttgctgg
gtgggctaacgatatccgcggatcccgactgatgcgtgaacgtgacggacgtaaccaccgcgacatgtgtgtgctgttc
