## Supplementary Information II. Extended Methods for "Optimizing protein production in the One-Pot Pure system: insights into reaction composition and expression efficiency"

***One-Pot PURE protein preparation***

The protocol was adapted from Grasemann *et al.*[1] with a few modifications detailed below.

*One-Pot Culture*

The overnight culture for One-Pot protein preparation was grown in 350 $\mu$L LB with 100 $\mu$g/mL carbenicillin and 1% glucose in a 1.3 mL 96-well plate (NEST Biotech). We found this volume to be optimal for growth while providing sufficient volume for optical density (OD) measurement and final culture inoculation. The frozen glycerol stock was removed from the -80 ˚C and placed on dry ice to prevent thawing. A cryo-replicator (Enzyscreen, CR1000) was used to transfer an aliquot of the frozen glycerol stock into an overnight culture plate, and we strongly recommend using a cryo-replicator to minimize cross-well contamination. A Breathe-Easier membrane (Diversified Biotech) was used to seal the overnight culture block, and the culture was grown for 12-16 hours overnight at 37 ˚C, 1000 rpm.

Additional overnight cultures for EF-Tu were prepared by inoculating frozen glycerol stock into two separate culture tubes with 3 mL LB with 100 $\mu$g/mL carbenicillin and 1% glucose and grown at 37 ˚C, 250 rpm for 12-16 hours. This was done to ensure optimal cell growth from frozen glycerol stock into liquid culture. Alternatively, the frozen glycerol for EF-Tu could be streaked onto an LB-agar plate with carbenicillin and glucose the day before overnight culture, and a population scrape of multiple colonies could be used to inoculate a singular 5.5 mL overnight culture for EF-Tu.

The following day, the $OD_{600}$ of the overnight culture was measured by diluting 100 $\mu$L of the overnight culture with 100 $\mu$L of 1xPBS in a clear-bottom 96-well plate (Corning) on a BioTek Synergy plate reader. For single strain One-Pot PURE, all cultures should exhibit similar, saturated $OD_{600}$ between 1.4 and 1.5.

For the starter culture, 1 L of pre-warmed LB was completed with 100 $\mu$g/mL carbenicillin and placed into a 2.5 L baffled flask (UltraYield). 150 $\mu$L of each overnight culture from the 96-well plate and 5 mL of the EF-Tu overnight culture were transferred into the final culture. The final culture was sealed with a breathable membrane (AirOtop) and grown at 37 ˚C, 250 rpm until $OD_{600}$ reached above 0.8 in 2 - 2.5 hours, after which 0.1 mM of IPTG was added to initiate protein expression at 37 ˚C, 250 rpm after IPTG induction.

After 3 hours of protein expression, the cell pellets were harvested by centrifuging the culture at 5,000 xg for 20 minutes at 4 ˚C. The cell pellet was transferred from the centrifuge bottle into the 50 mL Falcon tube by resuspending it with 25 mL spent LB and then transferred into a 50 mL Falcon tube to spin at 4000 xg for 10 minutes at 4 ˚C. The supernatant was discarded, and the cell pellet was flash-frozen in liquid nitrogen and placed at -80 ˚C until protein purification.

*36-Pot Culture*

The previously mentioned cell culture protocol was further modified for preparing 36-Pot PURE.

Instead of combining the overnight cultures, the strains were grown and individually induced, only being combined before the centrifugation step.

The 5 mL of EF-Tu overnight culture was transferred into a 2.5 L flask containing 470 mL of pre-warmed LB media with 100 $\mu$g/mL ampicillin. For all other strains, 150 $\mu$L of each was transferred into 50mL Falcon tubes containing 15.1mL of pre-warmed LB media with 100 $\mu$g/mL ampicillin. The cultures were then grown at 37 ˚C, 250 rpm, and monitored until $OD_{600}$ reached 0.8 (taking ~2 hours), after which 0.1 mM of IPTG was added to initiate protein expression. After 3 hours of protein expression, all 36 cultures were combined and pelleted as previously described.

*Protein Purification*

On the day of protein purification, the frozen cell pellet was thawed on ice for 30 minutes. 10 mL of Lysis Buffer (50 mM HEPES-KOH pH 7.6, 100 mM $NH_4Cl$, 500 mM NaCl, 10 mM $MgCl_2$, 1 tablet of cOmplete mini EDTA-free protease inhibitor (Roche), and 1 mM TCEP) was used to fully resuspend the cell pellet. The cell pellet resuspension was placed in an ice-water bath and sonicated at 20 kHz, 60 % amplitude, with 2-3 repeats of a 5-minute cycle with 5 seconds on and 5 seconds off pulses. The lysed cells were centrifuged at 12,000 xg for 30 minutes to purify the cell lysate from debris for further purification.

An Econo-Pac chromatography column (Bio-Rad) was loaded with 2 mL of Ni-Sepharose resin (Cytiva). After draining the storage buffer by gravity flow, 50 mL of double-distilled water was passed through the column, followed by 30 mL of the Wash Buffer (50 mM HEPES-KOH pH 7.6, 100 mM $NH_4Cl$, 500 mM NaCl, 10 mM $MgCl_2$, 1 mM TCEP, and 20 mM Imidazole-HCl pH 7) to equilibrate the column environment. After equilibration, a stopper was placed at the end of the column, and the cell lysate was added to the resin. The lysate-resin resuspension was placed on a rocker and incubated at 4 ˚C for 1 hour to maximize protein binding to Ni-Sepharose resin. After incubation, the lysate was drained by gravity flow, followed by a wash step with 40 mL of Wash Buffer. To elute the His-tagged One-Pot PURE proteins, 5 mL of Elution Buffer (50 mM HEPES-KOH pH 7.6, 100 mM $NH_4Cl$, 500 NaCl, 10 mM $MgCl_2$, 1 mM TCEP, and 500 mM Imidazole-HCl pH 7).

The eluted proteins are dialyzed in 400 mL of Dialysis Buffer (50 mM HEPES-KOH pH 7.6, 100 mM KCl, 10 mM $MgCl_2$, 30 v/v% Glycerol, 1 mM TCEP) using a 3.5k MWCO dialysis cassette (Thermo Scientific) at 4 ˚C for 30 minutes. The Dialysis Buffer was exchanged 3 more times for a total of 4 cascading steps to maximize Imidazole removal.

Following dialysis, One-Pot PURE proteins were concentrated to approximately 1 mL using a 3 kDa MWCO centrifugal filter (Amicon). The concentration step takes approximately 1 hour at 3500 xg and 4 ˚C. Protein concentrations were measured using the Pierce BCA Protein Assay kit (Thermo Scientific). The proteins are then divided into 5-10 $\mu$L aliquots in PCR strips using a multi-dispenser pipet, flash-frozen in liquid nitrogen, and stored at -80 ˚C.

### *One-Pot PURE energy mix preparation*

The energy mix preparation was prepared following the protocol of Grasemann *et al.*[1] with slight modifications. In brief, the 2.5x Energy Mix was prepared without magnesium acetate, such that the optimal $Mg^{2+}$ concentration could be calibrated to each batch of One-Pot PURE proteins. To prepare the amino acid mixture, 1 M KOH was used to dissolve 13 amino acids (Asn, Asp, Cys, Glu, Gln, His, Ile, Leu, Lys, Met, Phe, Trp, Tyr) to make 500 mM stocks. The remaining amino acids (Ala, Arg, Gly, Pro, Ser, Thr, Val) were dissolved in water. Except for 4 amino acids (Ala, Gly, Leu, Val) added at 1.67x concentration (final concentration 16.7 mM), equal volumes of the 500 mM amino acid stocks were combined and titrated to pH 7.5 using acetic acid to make the 10 mM amino acids solution.

### *Preparation of purified T7 RNA Polymerase*

T7 RNA Polymerase (RNAP) was purified using the Caltech Protein Expression Center (PEC). Briefly, plasmid coding for T7 RNAP expression (Addgene #124138) was transformed into *E. coli* M15/pREP4 and grew overnight on an LB-agar plate with 100 $\mu$g/mL carbenicillin and 25 $\mu$g/mL kanamycin. One colony was used to inoculate a 5 mL overnight culture of LB with carbenicillin and kanamycin grown at 37 ˚C and 250 rpm. The entire overnight culture was used to inoculate the final culture in 1 L Terrific Broth with carbenicillin and kanamycin. Cells were grown until an OD between 0.4-0.8 before induction of protein expression with 1 mM IPTG. Protein expression was done overnight at 27 ˚C at 250 rpm.

Following protein expression, the cell pellet was harvested by centrifuge culture at 5000 xg for 10 min in 4 ˚C. Cell pellets were resuspended in 150 mL 1xTBS supplemented with EDTA-free cOmplete Protease Inhibitor (Roche) and passed through the cell disruptor. The cell lysate was centrifuged at 15,000 xg to remove cell debris before loading on a TBS-equilibrated HisTrap column at a flow rate of 1 mL/minute. The HisTrap column was washed with 10 column volumes of the PEC Wash Buffer (30 mM HEPES-KOH pH 7.6, 150 mM NaCl, 20 mM Imidazole-HCl pH 7) and eluted with 5 column volumes of PEC Elution Buffer (30 mM HEPES-KOH pH 7.6, 150 mM NaCl, 250 mM Imidazole-HCl pH 7) at 0.5 mL/minute. The eluted protein was buffer-exchanged and concentrated using the PEC Dialysis Buffer (50 mM $NaHPO_4$, 100 mM NaCl, 2% DMSO, pH 7.5 with KOH). The protein concentration was determined using Pierce BCA Protein Assay (Thermo Scientific) before being aliquoted into working volumes, flash-frozen in liquid nitrogen, and stored at -80 ˚C until use.

### *Preparation of purified deGFP standards*

Green fluorescence protein deGFP was purified through Caltech PEC in a similar workflow as the T7 RNAP purification described previously. Slight differences in *E. coli* strain, expression vector, and downstream protein processing are detailed below. Briefly, plasmid coding for deGFP expression (sequence in **Supplementary Information I**) was transformed into *E. coli* BL21(DE3) and grew overnight on an LB-agar plate with 100 $\mu$g/mL carbenicillin. Following cell growth, expression, and purification, an additional size-exclusion chromatography step was performed to maximize protein purity. Purified deGFP proteins were exchanged to SEC Buffer (20 mM Tris-HCl pH 7.4 and 300 mM NaCl), aliquoted into working volumes, flash frozen in liquid nitrogen, and stored at -80 ˚C until use.

***Liquid chromatography with tandem mass spectrometry (LC-MS/MS) workflow***

Purified proteins were digested using EasyPep™ Mini MS Sample Prep Kits (Thermo Fisher Scientific) according to the manufacturer's protocol. After desalting and drying, peptides were suspended in Solvent A (0.2% formic acid and 2% acetonitrile) for further LC-MS/MS analysis. LC-MS/MS analyses of the digested peptides were performed on an EASY-nLC 1200 (Thermo Fisher Scientific) coupled to a Q Exactive HF hybrid quadrupole-Orbitrap mass spectrometer (Thermo Fisher Scientific). Peptides were separated on an Aurora UHPLC column (25 cm × 75 $\mu$m, 1.6 $\mu$m C18, AUR2-25075C18A, Ion Opticks) with a flow rate of 0.35 $\mu$L/min for a total duration of 65 min ionized at 1.8 kV in the positive ion mode. The gradient was composed of 6% Solvent B (80% ACN and 0.2% formic acid) for 3.5 min, 6–25% Solvent B for 42 min, 25–40% Solvent B for 14.5 min, and 40–98% Solvent B for 15 min.

MS1 scans were acquired at a resolution of 60,000 from 350 to 1600 m/z, with an AGC target of $3 \times 10^6$, and a maximum injection time of 15 ms. In MS2 scans, 12 abundant ions were acquired at a resolution of 30,000, with an AGC target of $1 \times 10^5$, a maximum injection time of 45 ms, and a normalized collision energy of 28. Dynamic exclusion was set to 45 sec, and ions with charges +1, +6, +7, +8, and greater than +8 were excluded. The temperature of the ion transfer tube was set to 275 °C, and the S-lens RF level was set to 60.

MS2 fragmentation spectra were searched with Proteome Discoverer SEQUEST (version 2.5; Thermo Scientific, Waltham, MA) against an *in silico* tryptic digested *E. coli* BL21 (DE3) and *E. coli* M15 database. The maximum number of missed cleavages was set to 2. Dynamic modifications were set to oxidation on methionine (M, + 15.995 Da), protein N-terminal acetylation (+42.011 Da), and Met-loss (–130.040 Da). Carbamidomethylation on cysteine residues (C, + 57.021 Da) was set as a fixed modification. The maximum parental mass error was set to 10 ppm, and the MS2 mass tolerance was set to 0.6 Da. The false discovery threshold was strictly set to 0.01 using the Percolator Node validated by q-value. The relative abundance of parental peptides was calculated by integrating the area under the curve of the MS1 peaks using the Minora LFQ node.

***Computational prediction of OmpT proteolysis on PURE proteins***

We constructed an amino acid probability matrix to score 8-residue subsequences based on a predefined set of probability patterns for each amino acid (**Table SI.II 1**), which was sourced from empirical results aggregated on MEROPS, the Peptidase Database. Each entry in this matrix represents the frequency of observing a particular amino acid at a specific position in an 8-amino-acid sequence.[2]

**Table SI.II 1.** Amino acid substrate specificity matrix for OmpT cleavage.

| **Cleavage Pattern Specificity Matrix (**-/-/a/RK † Rkq/a/a/g**)** | | | | | | | | |
|---|---|---|---|---|---|---|---|---|
| **Amino Acid** | **P4** | **P3** | **P2** | **P1** | **P1'** | **P2'** | **P3'** | **P4'** |
| G | 4 | 5 | 7 | 0 | 0 | 8 | 6 | 17 |
| P | 1 | 1 | 0 | 0 | 0 | 0 | 1 | 3 |
| A | 3 | 7 | 14 | 0 | 0 | 12 | 12 | 6 |
| V | 1 | 2 | 0 | 0 | 2 | 7 | 3 | 0 |
| L | 9 | 6 | 4 | 0 | 0 | 5 | 2 | 0 |
| I | 2 | 1 | 2 | 0 | 0 | 5 | 0 | 2 |
| M | 2 | 0 | 0 | 0 | 1 | 0 | 1 | 2 |
| F | 1 | 2 | 1 | 0 | 0 | 0 | 1 | 0 |
| U | 0 | 2 | 5 | 0 | 0 | 1 | 0 | 0 |
| W | 4 | 2 | 0 | 0 | 0 | 0 | 3 | 0 |
| S | 0 | 2 | 2 | 1 | 1 | 2 | 1 | 1 |
| T | 1 | 2 | 3 | 0 | 0 | 1 | 1 | 2 |
| C | 1 | 0 | 0 | 0 | 0 | 0 | 0 | 0 |
| N | 0 | 0 | 2 | 0 | 0 | 1 | 0 | 0 |
| Q | 1 | 1 | 4 | 0 | 7 | 0 | 4 | 2 |
| D | 4 | 1 | 0 | 0 | 0 | 0 | 0 | 4 |
| E | 5 | 0 | 5 | 0 | 0 | 1 | 4 | 4 |
| K | 2 | 5 | 0 | 16 | 13 | 4 | 1 | 2 |
| R | 5 | 8 | 1 | 37 | 30 | 3 | 7 | 3 |
| H | 1 | 0 | 0 | 0 | 0 | 0 | 0 | 1 |

*Normalization of the Pattern Matrix*

The matrix entries, representing counts, were normalized to probabilities so that each set of positional values for an amino acid sums to 1. The normalized probability for a given amino acid, $aa$, at position, $i$, was computed as:

$$P(aa_i) = \frac{x_i}{\sum_{i=1}^{8} x_i}$$

where $x_i$ is the raw count for the amino acid at position $i$, and the sum is taken over all positional values for that amino acid.

*Scoring Function for Subsequence Patterns*

To calculate the score of a subsequence, we defined a probabilistic scoring function that multiplies the probabilities for each amino acid in the subsequence based on its position:

$$Score(S) = \prod_{i=1}^{8} P(S_i)$$

where $S_i$ is the amino acid in position $i$ in the subsequence $S$, and $P(S_i)$ is the probability of observing that amino acid at the position $i$ according to the normalized matrix. If the subsequence contains an amino acid not present in the matrix at that position, the score is set to zero.

*Pattern Search Algorithm*

The search for subsequences with high probability scores was performed using a sliding window approach. For each sequence in a FASTA file, we iteratively considered all possible 8-residue subsequences, calculated their scores, and retained those that exceeded a predefined score threshold. The steps are as follows:

1. Parse the FASTA file to extract protein sequences.
2. Slide an 8-residue window across each sequence.
3. For each 8-residue subsequence, compute its score using the previously defined formula.
4. Retain the subsequence if its score exceeds a specified threshold (in this case, $10^{-5}$).

*AlphaFold3 predictions*

To characterize the susceptibility of One-Pot PURE proteins with predicted cleavage sites to OmpT proteolysis, we used the AlphaFold3 (AF3) protein structure prediction tool[3] to model the protein structure and visualize the structure using PyMOL. Proteins with susceptible peptide(s) exposed on the external surface are more likely to be susceptible to OmpT proteolysis, as the peptide sites are accessible. Proteins with susceptible peptide sites buried were determined to be likely to be cleaved by OmpT.
