## Supplementary Inforation III. Supplementary Figures for "Optimizing protein production in the One-Pot Pure system: insights into reaction composition and expression efficiency"

### Table of Figures


| | | |
|---|---|---|
| **Figure SI.III 1** | Monoculture growth profile of *E. coli* strains M15/pREP4 and BL21(DE3) expressing PURE proteins | 3 |
| **Figure SI.III 2** | PURE protein expression and monoculture growth profile using strains directly obtained from a collaborator's lab | 4 |
| **Figure SI.III 3** | Cell growth and protein expression comparison of *E. coli* BL21(DE3) cells expressing 10 PURE proteins grown in LB and LB with 1 % w/v glucose | 5 |
| **Figure SI.III 4** | AlphaFold simulation of the eight PURE proteins containing peptides susceptible to OmpT proteolysis: myokinase (MK), elongation factor – thermal stable (EF-Ts), creatine kinase (CK), glycyl-tRNA synthetase – beta subunit (GlyRS β), T7 RNA polymerase (T7 RNAP), isoleucyl-tRNA synthetase (IleRS), threonyl-tRNA synthetase (ThrRS), valyl-tRNA synthetase (ValRS). Nearly all susceptible peptides (highlighted in purple) are exposed on the protein's surface, except for peptide WARMIMM (R517) for ValRS and EVNKVIEQ peptide (K821) for IleRS. | 6 |
| **Figure SI.III 5** | Cell growth and protein expression comparison of *E. coli* M15/pREP4 and BL21(DE3) cells expressing same PURE proteins | 7 |
| **Figure SI.III 6** | Comparison of the relative abundance of individual PURE proteins in the Single Strain One-Pot PURE and commercial PURE systems. | 8 |
| **Figure SI.III 7** | Productivity comparison of the collaborator-produced batches of Dual Strain 36-Pot A and B, and Single Strain One-Pot PURE ($Single_C$). | 9 |
| **Figure SI.III 8** | Kinetic time trace of deGFP production for Dual- and Single-Strain One-Pot PURE system in commercial $PURE_A$ energy mix | 10 |
| **Figure SI.III 9** | Kinetic time trace of deGFP production for Dual- and Single Strain One-Pot PURE system in commercial $PURE_B$ energy mix | 11 |

| | | |
|---|---|---|
| **Figure SI.III 10** | Plate reader fluorescence calibration for purified deGFP and synthesized malachite green aptamer (MGA) | 12 |

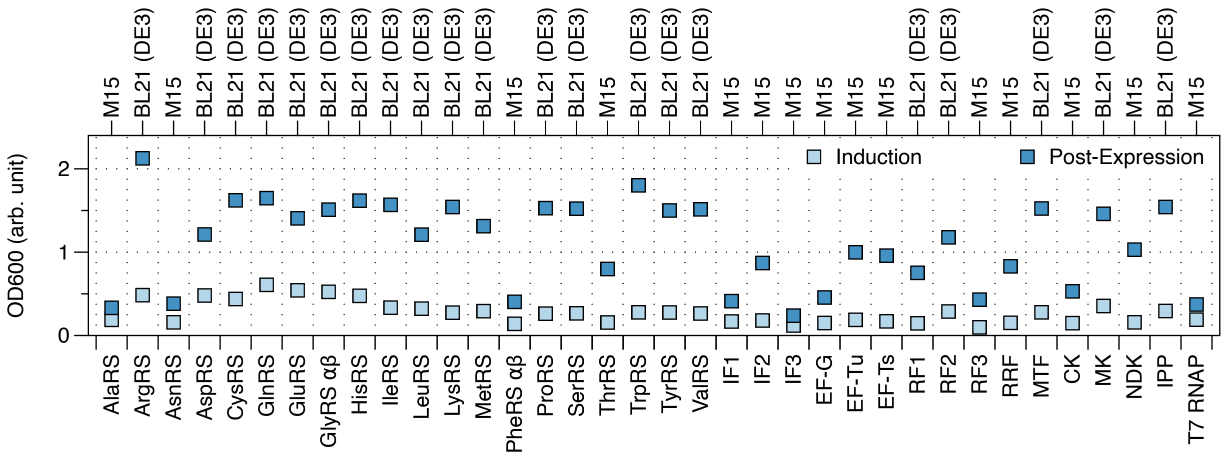


**Figure SI.III 1.** Monoculture growth profile of *E. coli* M15/pREP4 and BL21(DE3) strains expressing PURE proteins. Light blue squares represent cells' $OD_{600}$ measured at the time of IPTG induction. Darker blue squares represent cells' $OD_{600}$ measured 3 hours post-protein expression.

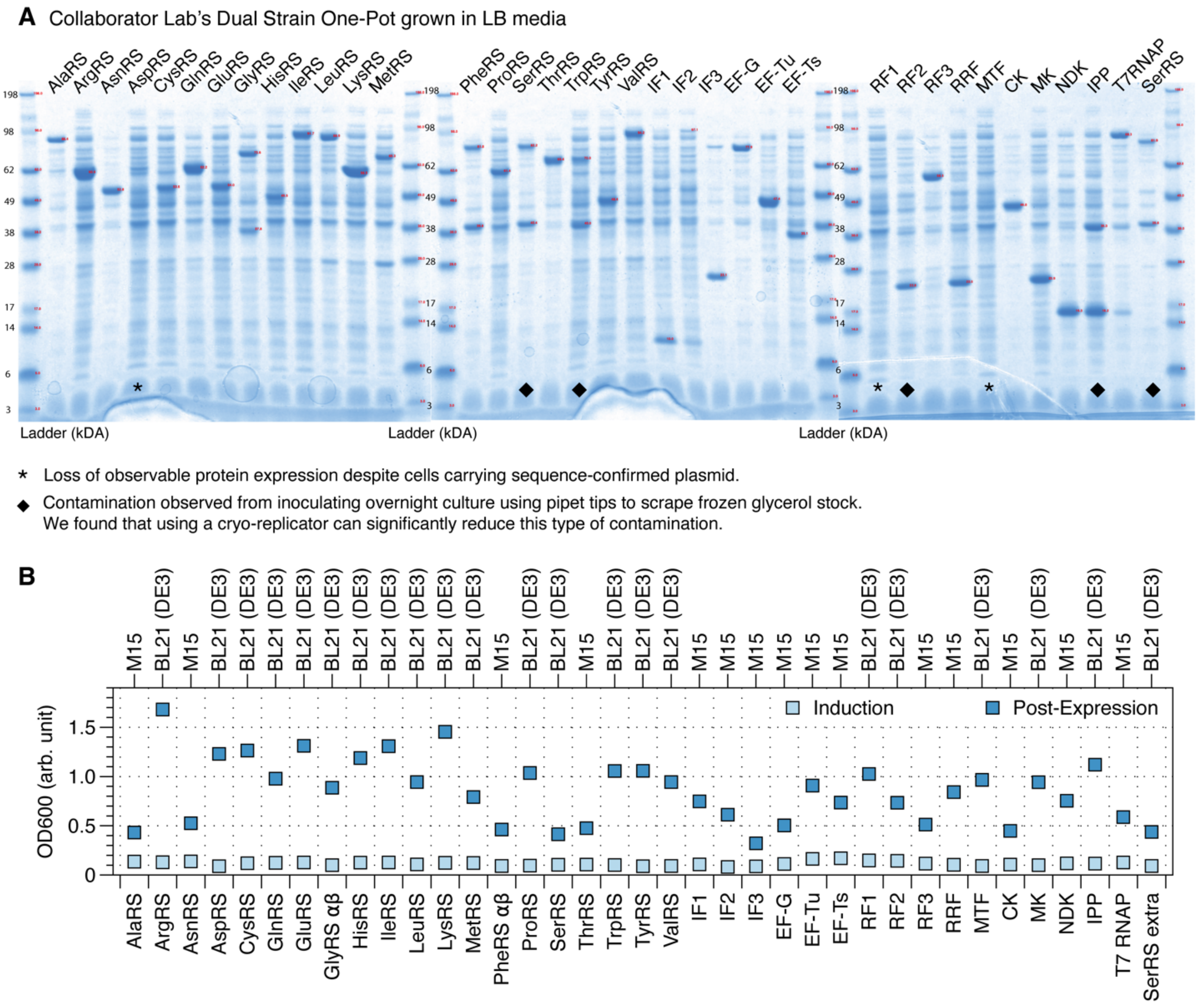


**Figure SI.III 2. (A)** Each PURE protein expression level was assessed on SDS-PAGE using cells directly obtained from a collaborator's lab. **(B)** Monoculture growth profile of *E. coli* M15/pREP4 and BL21(DE3) cells expressing each PURE protein. Light blue squares represent cells' $OD_{600}$ measured at the time of IPTG induction. Darker blue squares represent cells' $OD_{600}$ measured at 3 hours after protein expression. SerRS extra represents an extra culture of SerRS.

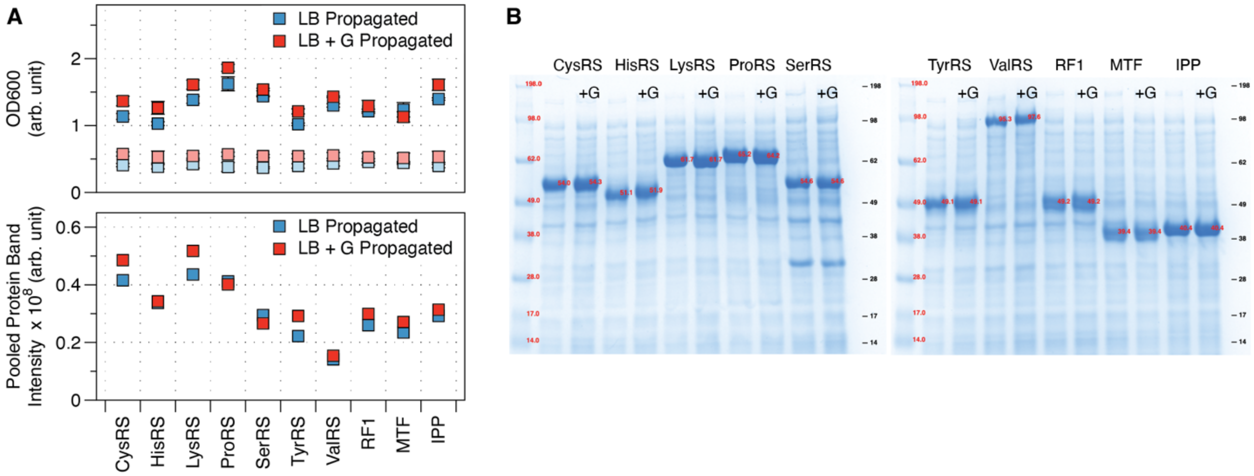


**Figure SI.III 3. (A)** Growth profile of fresh transformations of *E. coli* BL21(DE3) cells expressing 10 PURE proteins in LB and LB with 1% w/v glucose monocultures (top). Light blue and dark blue squares represent cells' $OD_{600}$ measured at the time of IPTG induction and at 3 hours after protein expression for cells grown in LB media, respectively. Light red and dark red squares represent cells' $OD_{600}$ measured at the time of IPTG induction and at 3 hours after protein expression for cells grown in LB media with 1% w/v glucose, respectively. Each square is the average of biological triplicates, and error bars represent standard deviations of biological triplicates. Biological triplicates of cells expressing One-Pot PURE proteins were combined and run on SDS-PAGE as pooled protein. Pooled protein band intensity (bottom) extracted from gel images are plotted. **(B)** SDS-PAGE gel of pooled protein expression from cells plated and grown in LB without or with 1% w/v glucose.

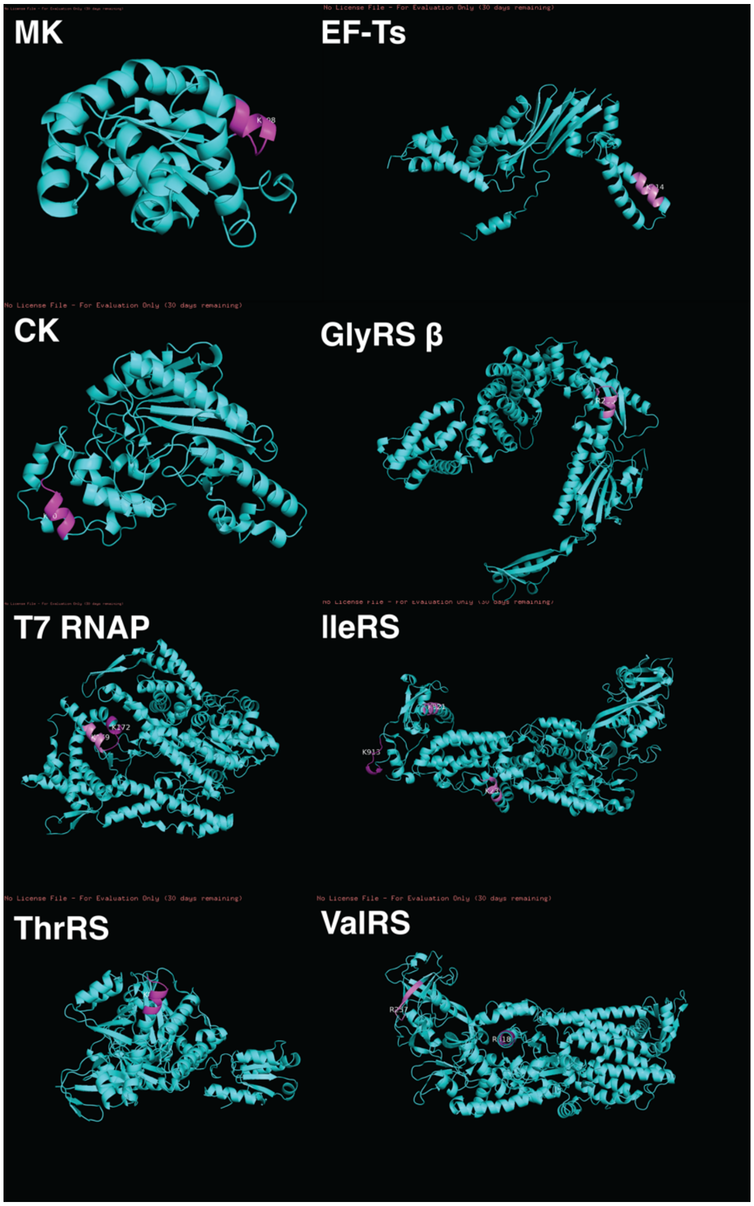


**Figure SI.III 4.** AlphaFold simulation of the eight PURE proteins containing peptides susceptible to OmpT proteolysis: myokinase (MK), elongation factor – thermal stable (EF-Ts), creatine kinase (CK), glycyl-tRNA synthetase – beta subunit (GlyRS β), T7 RNA polymerase (T7 RNAP), isoleucyl-tRNA synthetase (IleRS), threonyl-tRNA synthetase (ThrRS), valyl-tRNA synthetase (ValRS). Nearly all susceptible peptides (highlighted in purple) are exposed on the protein's surface, except for peptide WARMIMM (R517) for ValRS and EVNKVIEQ peptide (K821) for IleRS.

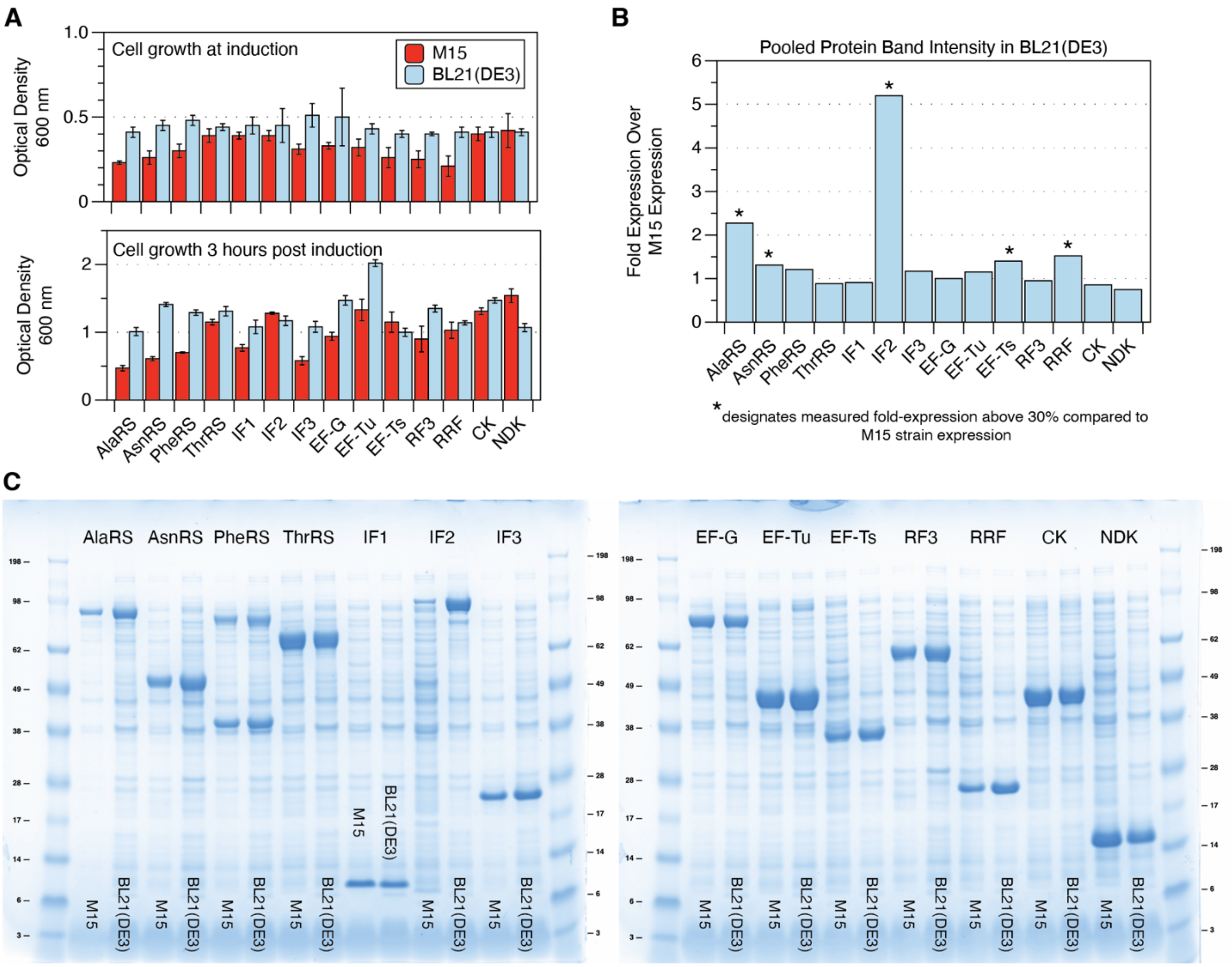


**Figure SI.III 5. (A)** Growth comparison of the same PURE protein expressed in *E. coli* M15/pREP4 and BL21(DE3) cells. BL21(DE3) strain expressing the same PURE protein exhibited better growth at the time of IPTG induction and 3 hrs after protein expression. Bar graphs represent the average of biological triplicates, and error bars represent the standard deviation of biological triplicates. **(B)** Comparison of protein expression levels in BL21(DE3) strain to M15/pREP4 strain using signal intensity extracted from SDS-PAGE gels. Five of 15 proteins, denoted by asterisks (*), exhibited at least 30% better expression when grown in BL21(DE3) than M15/pREP4. **(C)** SDS-PAGE gel showing individual protein bands pooled from biological triplicates, with expression levels from M15/pREP4 and BL21(DE3) juxtaposed.

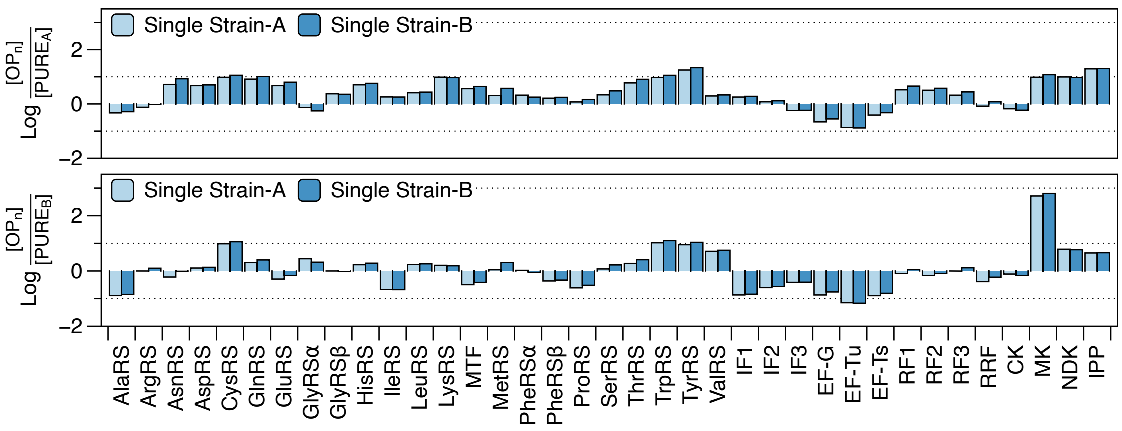


**Figure SI.III 6.** Comparison of the relative abundance of individual PURE proteins in the Single Strain One-Pot PURE and commercial PURE systems. The relative protein compositions are scaled to 2.5 mg/mL for Single Strain One-Pot PURE proteins in reaction and the manufacturers' recommended reaction composition for commercial PURE.

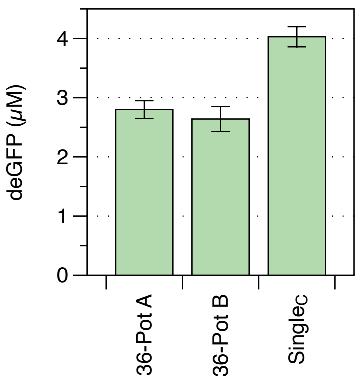


**Figure SI.III 7.** Productivity comparison of the collaborator-produced batches of Dual Strain 36-Pot A and B, and Single Strain One-Pot PURE ($Single_C$). PURE reactions were assembled at 5 mg/mL of PURE protein, using homemade energy mixes and 5 nM of $P_{T7}$-UTR1-deGFP plasmid. The bar graphs represent the average terminal protein yield of reaction triplicates, with error bars indicating the standard deviations of reaction triplicates.

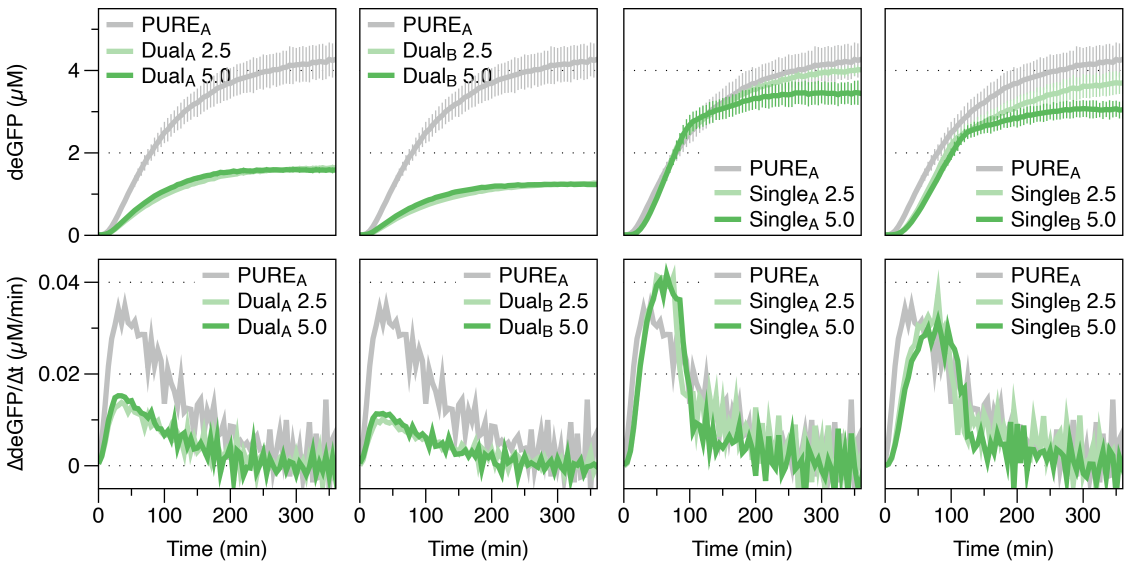


**Figure SI.III 8.** Kinetic time trace of deGFP production for Dual- and Single-Strain One-Pot PURE system in commercial $PURE_A$ energy mix (top). Plots represent the average of reaction triplicates, and error bars represent the standard deviation of reaction triplicates. The protein translation rate of Dual Strain and Single Strain One-Pot PURE reaction compared to the commercial $PURE_A$ reference reaction (bottom). Plots represent the average rate of reaction triplicates.

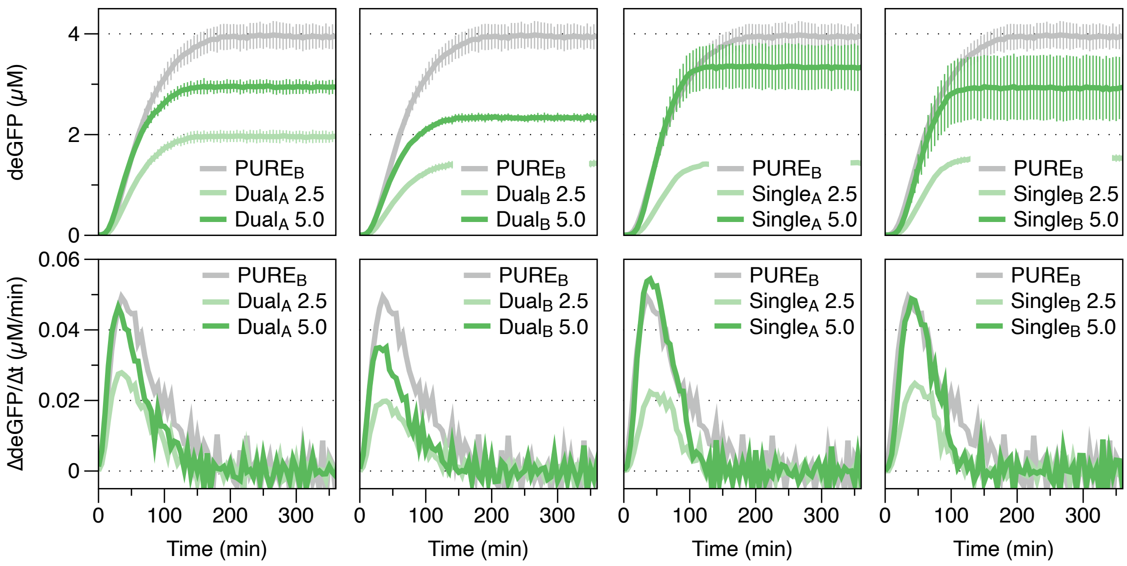


**Figure SI.III 9.** Kinetic time trace of deGFP production for Dual- and Single Strain One-Pot PURE system in commercial $PURE_B$ energy mix (top). Plots represent the average of reaction triplicates, and error bars represent the standard deviation of reaction triplicates. The protein translation rate of Dual Strain and Single Strain One-Pot PURE reaction compared to the commercial $PURE_B$ reference reaction (bottom). Plots represent the average rate of reaction triplicates.

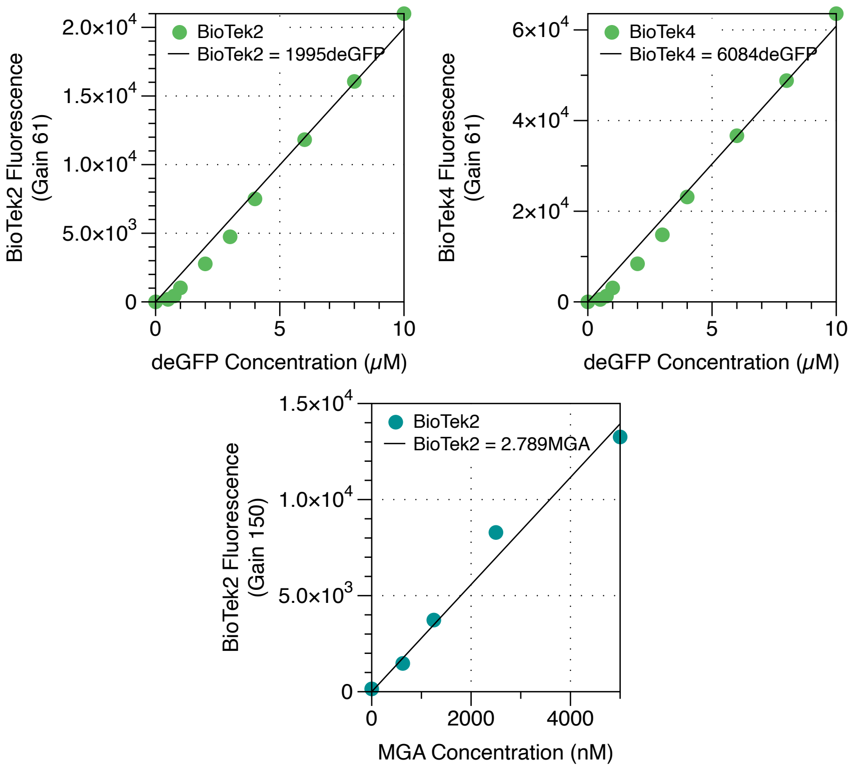


**Figure SI.III 10.** Plate reader fluorescence calibration. Purified deGFP proteins were prepared at varying concentrations by diluting in 1xPBS. Dilutions of deGFP proteins were added onto 384-well low-volume plates at 5 $\mu$L, and fluorescence was measured at 485/515 excitation/emission wavelengths (nm/nm) at a gain setting of 61. A linear line was fit to the data (top). Synthesized malachite green aptamer (MGA) sequence was prepared with 10 $\mu$M malachite green dye and 1 U/$\mu$L Rnase Inhibitor, Murine (NEB) in 1xPBS. Dilutions of MGA were added onto 384-well low-volume plates at 5 $\mu$L, and fluorescence was measured at 610/650 excitation/emission wavelengths (nm/nm) at a gain setting of 150. A linear line was fit to the data (bottom).
